# Groundfish biodiversity change in northeastern Pacific waters under projected warming and deoxygenation

**DOI:** 10.1101/2022.05.04.490690

**Authors:** Patrick L. Thompson, Jessica Nephin, Sarah C. Davies, Ashley E. Park, Devin A. Lyons, Christopher N. Rooper, M. Angelica Peña, James R. Christian, Karen L. Hunter, Emily Rubidge, Amber M. Holdsworth

## Abstract

In the coming decades, warming and deoxygenation of marine waters are anticipated to result in shifts in the distribution and abundance of fishes, with consequences for the diversity and composition of fish communities. Here, we combine fisheries independent trawl survey data spanning the west coast of the USA and Canada with high resolution regional ocean models to make projections of how 34 groundfish species will be impacted by changes in temperature and oxygen in British Columbia (B.C.) and Washington. In this region, species that are projected to decrease in occurrence are roughly balanced by those that are projected to increase, resulting in considerable compositional turnover. Many, but not all, species are projected to shift to deeper depths as conditions warm, but low oxygen will limit how deep they can go. Thus, biodiversity will likely decrease in the shallowest waters (< 100 m) where warming will be greatest, increase at mid depths (100—600 m) as shallow species shift deeper, and decrease at depths where oxygen is limited (> 600 m). These results highlight the critical importance of accounting for the joint role of temperature, oxygen, and depth when projecting the impacts of climate change on marine biodiversity.

## Introduction

In marine ecosystems, climate change is expected to result in warmer waters and reduced dissolved oxygen, both of which are likely to impact organismal performance, population growth rates, and viability (Prince *et al.* 2010; Deutsch *et al.* 2015). There is evidence that marine species are more sensitive to temperature changes than terrestrial species, and many species are already shifting their distributions to track changing conditions (Poloczanska *et al.* 2013; Pecl *et al.* 2017; Pinsky *et al.* 2021). Therefore, we expect to see considerable reorganization in the composition of marine communities as conditions continue to change (Bartley *et al.* 2019; Pinsky *et al.* 2020). Projections of how species and communities will respond to climate change over the coming decades are needed in order to develop management strategies for preserving biodiversity and ensuring the sustainability of fisheries and ecosystems (Wilson *et al.* 2020).

In marine environments, species are strongly influenced by temperature and dissolved oxygen (Prince *et al.* 2010; Brown & Thatje 2015; Deutsch *et al.* 2015; Clarke *et al.* 2021), both of which are projected to change in the coming decades (e.g., Holdsworth *et al.* 2021). When available, experimental measurements of species specific thermal tolerances and oxygen requirements, including how temperature influences oxygen requirements, can be used to assess how species distributions and abundances will shift as the oceanographic conditions change (i.e., via the metabolic index; Deutsch *et al.* 2015; Penn *et al.* 2018; Sunday *et al.* 2022). However, because such measurements are not available for most marine fish, this approach cannot be applied to assess how climate change will impact the overall biodiversity and composition of marine communities (but see Clarke *et al.* 2022 for a potential alternative approach).

Instead, species distribution models (SDMs) can be used to leverage current species–environment associations across space to estimate species specific temperature and oxygen responses (Guisan & Thuiller 2005). SDMs are increasingly being applied in marine environments to project how species will respond to future conditions. These models generally project that species’ ranges will shift polewards as conditions warm (García Molinos *et al.* 2016). Range shifts for species in west coast North American waters are projected to be particularly large, with many species’ range limits expected to change by more than 1000 km by the end of the century under high greenhouse gas emissions scenarios (Morley *et al.* 2018). This result is consistent with estimates that suggest that the range edges of marine species are generally, but not universally, changing with warming ocean water temperatures (Sunday *et al.* 2012; Fredston *et al.* 2021). In addition, there is evidence that some, but not all, marine species are shifting to deeper depths as conditions warm (Chaikin *et al.* 2022). In Canadian Pacific waters, there is evidence that changes in temperature and oxygen are impacting the distribution and densities of groundfish species (English *et al.* 2022), but that these changes have not yet resulted in major changes in the diversity and composition of the groundfish community (Thompson *et al.* 2022a).

The robustness of SDMs depends on properly accounting for, and distinguishing, the influence of multiple environmental variables on species distributions. Most SDM models to date do not account for oxygen (e.g., Cheung *et al.* 2009; Ready *et al.* 2010; Morley *et al.* 2018; Chaikin *et al.* 2022), and in some cases, rely on sea surface temperature when modeling species that live on the seafloor (e.g., Ready *et al.* 2010; García Molinos *et al.* 2016). In addition, demersal marine communities (e.g., groundfish) are highly depth structured because of physiological limits to temperature, hydrostatic pressure (Brown & Thatje 2014), and hypoxia tolerance (Brown & Thatje 2015; Essington *et al.* 2022), as well as light, competition, and food availability (Elith & Leathwick 2009). However, because depth, temperature, and oxygen are closely correlated in many regions, it can be challenging to distinguish the role of each of these variables in determining contemporary species distributions. Some models have addressed this challenge by assuming that depth distributions are entirely determined by temperature, because this allows for projected distributions to shift into deeper waters as conditions warm (Pinsky *et al.* 2013; Morley *et al.* 2018). However, if depth—and associated variables (e.g., light and pressure)—influences species ranges directly, as well as by influencing water temperature and dissolved oxygen, then the direct effect of temperature and dissolved oxygen on species ranges cannot be determined without also accounting for the independent influence of depth.

Although climate SDM studies often focus on range shift rates and distances, climate change will also impact species within their range boundaries (English *et al.* 2022). Projections that capture local scale changes in species’ occurrences and abundances are particularly needed for the implementation of climate resilient management strategies such as establishing networks of marine protected areas and fisheries management areas (Wilson *et al.* 2020; O’Regan *et al.* 2021). The recent development of regional ocean models that downscale climate projections at a high-resolution (e.g., Peña *et al.* 2019; Holdsworth *et al.* 2021) offers the opportunity to make projections at spatial resolutions (e.g., <3 km) that are relevant to marine spatial planning initiatives (Frazão Santos *et al.* 2018; Gissi *et al.* 2019; Friesen *et al.* 2021).

Here we provide projections of groundfish community change in British Columbia (B.C.), Canada and Washington, U.S.A. waters under future climate change scenarios (2046–2065) that account for the combined effects of temperature, dissolved oxygen, and seafloor depth at a spatial scale that is relevant to marine spatial planning and fisheries management. We do this by using SDMs to estimate how temperature, oxygen, and seafloor depth determine the current occurrence and distribution of 34 groundfish species across the west coast from California to Alaska (Figure 1a), based on fisheries independent trawl surveys. We then combine these estimated species responses with two regional ocean models that cover B.C. and Washington waters to project changes in species occurrences from a historical baseline of 1986–2005 to a future period in 2046–2065 under Representative Concentration Pathways (RCP) 4.5 and 8.5 (Figure 1). This time frame was selected because it is short enough to be relevant for informing policy decisions over the coming decades, while being long enough to ensure that changes in temperature and dissolved oxygen are unambiguously due to increased greenhouse gases. Estimating species’ responses to the environmental variables using the spatially extensive trawl survey dataset allows us to capture a much wider range of environmental conditions than are present in the region where we make our projections. In particular, the tight correlation between depth and temperature in B.C. and Washington waters is not present across the coast-wide dataset, and this allows us to separate the influence of these variables using correlational SDMs.

**Figure 1:**
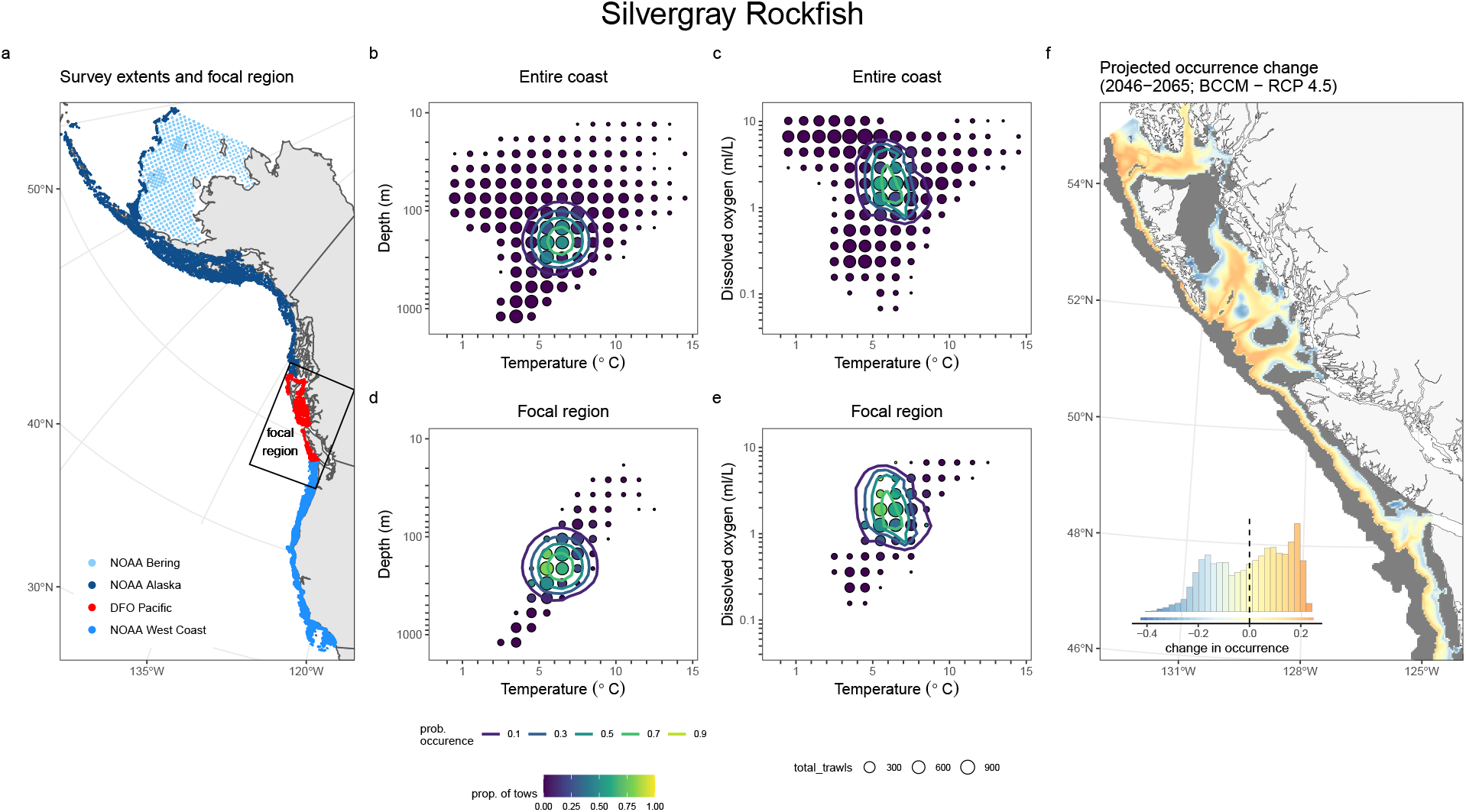
Overview of the SDM fitting and projection process using (Silvergray Rockfish, *Sebastes brevispinis*) as an example. Panel (a) shows the extent of the groundfish survey data, with each point representing a single trawl, coloured by survey. Focal region outlined in panel (a) corresponds to the extent of the regional ocean models that are used for the future projections. Panels c–e show the trawl data binned in environmental space for temperature by depth (b, d) and temperature by dissolved oxygen (c, e). The size of the circle shows the number of tows that fall into that bin, the color shows the proportion of those tows where the species was present. The SDM model fit is shown as contour lines of predicted occurrence. Panels b and c show the full coastwide dataset used to fit the SDMs, panels d and e show only the trawls from B.C. and Washington. Panel f shows the projected change in occurrence for each 3 km^2^ grid cell between the historical baseline (1986–2005) and the future projection (2046–2065), here based on the BCCM model and RCP 4.5. The inset histogram in panel (f) shows the distribution of values across all 3 km^2^ grid cells and provides a legend for the colours shown on the map. Grid cells where the probability of occurrence was projected to be below 0.1 in both the historical and future time periods periods are shown in grey and are not included in the histogram, as we do not consider these to be important habitat for the species. Plots for all species and projected changes from both climate models and both RCP scenarios are available in Appendix A.

## Methods

### Estimating species response curves

#### Fisheries independent trawl data

We estimated how the observed distribution of groundfish species is determined by temperature, dissolved oxygen, and seafloor depth using data from 31 239 fisheries independent scientific research trawls. This dataset spans the entire U.S. and Canadian west coast (Figure 1a) and includes both species presence and absence in the trawls. The Canadian portion of the coast is surveyed by Fisheries and Oceans Canada (DFO) with the Groundfish Synoptic Bottom Trawl Surveys which have been conducted since 2003 (DFO Pacific; 4218 trawls) (Sinclair *et al.* 2003; Anderson *et al.* 2019). The U.S. portions of the coast are surveyed by the National Oceanic and Atmospheric Administration (NOAA) and include the NWFSC West Coast Groundfish Bottom Trawl Survey (NOAA West Coast; 10 769 trawls), which has been conducted since 2003, and the Alaska Groundfish Bottom Trawl Surveys in the Aleutian Islands, the Gulf of Alaska, the Bering Sea upper continental slope, and the east and north Bering Sea shelf (Stauffer 2004; Keller *et al.* 2017). The Alaska Groundfish Bottom Trawl Surveys have been conducted since the 1980s, but our analysis only includes data from 2000 onward in order to align with the temporal extent of the DFO and NOAA West Coast surveys. Because surveys in the sub regions of Alaska use different protocols, our analysis considers them as two separate surveys: the surveys on the Bering Sea Shelf (NOAA Bering; 7365 trawls) and those in the Aleutian Islands, the Gulf of Alaska, and the Bering Sea upper continental slope (NOAA Alaska; 8887 trawls). We included data from all surveys up to 2019. We included all 77 groundfish species that were present in the overall trawl dataset that did not show obvious breakpoints in occurrence that corresponded with the boundaries of the three surveys. However, our final analysis included only the 34 species (listed in the relevant figures) for which the models had adequate forecasting ability, as outlined in the “assessing predictive accuracy” section below.

Temperature, depth, and dissolved oxygen for the observed data were obtained by CTD instrumentation and dissolved oxygen sensors deployed on the headrope of the trawls. Dissolved oxygen was available for 17.7% of the trawls. We filled in missing oxygen data using predictions from a Generalized Linear Mixed Effects Models fit with the sdmTMB package (Anderson *et al.* 2022a) that was also informed by an additional 7037 oxygen observations from the International Pacific Halibut Commission (IPHC) ocean profile data (Sadorus *et al.* 2016). Full details of the dissolved oxygen model are provided in the Supplementary Materials. This model was able to predict withheld data with an R^2^ of 0.952 (Figure S1, S2). The correlation between log depth and temperature was −0.904 within our focal region but was 3 × 10^−4^ across the full dataset. The correlation between log depth and log dissolved oxygen was −0.811 within our focal region and was −0.864 across the full dataset. The correlation between temperature and log dissolved oxygen was 0.742 within our focal region and was −0.221 across the full dataset.

#### Species distribution model

For each species, we modelled species occurrences in the coastwide trawl dataset using a Generalized Linear Model with a binomial distribution and a logit link function to map the linear predictors to the binary presence-absence data using the sdmTMB package (Anderson *et al.* 2022b) in R v.4.0.2 (R Core Team 2021). The fixed effects were temperature, log dissolved oxygen, log depth, and survey. We included quadratic terms for temperature and log depth to allow species occurrences to peak at intermediate values (Figure 1b–e). We fit a breakpoint function for log dissolved oxygen to reflect the fact oxygen is a limiting factor (Essington *et al.* 2022). That is, each species is expected to have an oxygen threshold, below which oxygen limitation is expected to reduce their probability of occurrence, but above this threshold species should not be sensitive to changes in oxygen concentrations (Fry 1971). A survey term specifying the data source—NOAA West Coast, NOAA Alaska, NOAA Bering, or DFO Pacific—was included as a catchability covariate to account for variation in detection probability across surveys due to differences in survey design and gear. We expected that these differences in detection probability should be small because our analysis was based on presence/absence data. The use of survey ID as a fixed effect in a presence/absence model has been demonstrated to sufficiently capture catchability differences between trawl and longline surveys (Thompson *et al.* 2022b), where differences in catchability are expected to be considerably larger than when comparing across surveys that all use similar trawling methods. We elected not to include spatial or spatiotemporal random fields in our models. While random fields offer an effective way of accounting for unmeasured environmental variables (Anderson *et al.* 2022b), they have the potential to absorb variation in species occurrences that is due to the environmental covariates, and thus result in an underestimate of species’ sensitivity to environmental change (Clayton *et al.* 1993; Marques *et al.* 2022).

In the majority of species, our models estimated reasonable dissolved oxygen responses–that is positive oxygen slopes, a breakpoint that fell within the range of observed oxygen conditions (i.e, < 10 mL L^−1^), and proper model convergence. However, for some of the species the breakpoint model estimated a negative slope above the breakpoint. We assume that this was not a real response to increasing oxygen but was due to the fact that depth and oxygen are correlated and so the model is erroneously attributing the fact that the species is not found in shallower waters (where oxygen tends to be higher) to oxygen. For species that did not meet this criteria, we elected to drop the oxygen response and model their occurrence based on temperature, depth, and survey.

#### Assessing forecasting accuracy

We assessed the forecasting accuracy of the SDM by comparing how well a model fit to only data from 2000–2010 could forecast species’ occurrences in trawls within our focal region for the period of 2011–2019. This assessment approximates the approach that we used in our projections (described below), but uses the latter half of our trawl data as testing data in order to estimate how well our models can predict future time periods that were not included in the training data. In particular, this post 2010 period includes the marine heatwave that occurred in the region from late 2013 to 2016 (Cavole *et al.* 2016), and so provides a test of how well the model can predict anomalous conditions without historical precedent. The final set of species included in this manuscript includes only those species that exceeded a threshold Tjur R^2^ (Tjur 2009) of 0.2 and an area under the curve (AUC; Pearce & Ferrier 2000) of 0.75 based on the temporal forecasting accuracy assessment. Tjur R^2^ quantifies the mean difference in forecasted occurrence between trawls where the focal species is present and trawls where it is absent (Tjur 2009) and so assesses how good the model is at distinguishing presence/absence patterns. Note that this temporal blocking of training and testing data was only used for assessing the predictive accuracy of the models; the models used to fit the SDMs that were used to make the projections for the 2045–2065 period (described in subsequent sections) included all trawls in the dataset.

### Projecting groundfish biodiversity changes in British Columbia and Washington waters

#### Regional ocean models

We based our groundfish biodiversity change projections on two regional models that downscale climate projections: the British Columbia Continental Margin model (BCCM; Peña *et al.* 2019) and the North-Eastern Pacific Canadian Ocean Ecosystem model (NEP36-CanOE; Holdsworth *et al.* 2021). The BCCM model is an implementation of the Regional Ocean Modeling System (ROMS; Haidvogel *et al.* 2008) at a 3 km horizontal resolution. The NEP36 model is an implementation of Nucleus for European Modeling of the Ocean (NEMO) numerical framework 3.6 (Madec 2016) at a variable grid spacing between 1.5 and 2.25 km. Both models used anomalies from the Canadian Earth System Model version 2 (Arora *et al.* 2011) at the open boundaries and the Canadian Regional Climate Model version 4 (CanRCM4; Scinocca *et al.* 2016) for the surface boundary conditions to downscale climate projections.

The BCCM model outputs were interpolated from a curvilinear to a regular 3 km grid using a thin plate spline using the fields (Nychka *et al.* 2017) and raster packages (Hijmans 2022). The NEP36 model outputs were interpolated to the same 3 km grid using a linear interpolation. We used a historical baseline of 1986 to 2005 because these years were present in the historical hindcast of the BCCM as well as the historical climatology of the NEP36. We used projected values from these models for 2046 to 2065 based on RCP 4.5 and 8.5. RCP 4.5 represents a scenario with moderate climate change mitigation and RCP 8.5 represents a no mitigation, worst case scenario (Moss *et al.* 2010; Arora *et al.* 2011; Hausfather & Peters 2020). We elected to use 20-year climatologies for the historical baseline and future scenarios to reduce the influence of year-to-year variation in climate (i.e., natural variability).

We used mean summer (April–September) near-bottom temperature and dissolved oxygen averaged across all years in the historical baseline and future projection periods (Figures S3 - S5). Historical temperatures from both models and dissolved oxygen from the BCCM model were comparable to those observed in the research trawl surveys, but oxygen concentrations from the NEP36 model were consistently high. Therefore, we bias corrected the NEP36 data based on the BCCM hindcast as a common baseline. We elected to use proportional change rather than absolute change for this bias correction to avoid negative projected oxygen concentrations. We did this by calculating the proportional change in oxygen and temperature between the historical and future projections for each 3 km^2^ grid cell as: *x_f_/x_h_*, where *x_f_* represents the future projected value and *x_h_* represents the historical value (Navarro-Racines *et al.* 2020). We then multiplied these proportional changes by the historical BCCM values to obtain future projections that were bias corrected to the BCCM baseline. We calculated proportional changes in temperature in Kelvin and then converted these back to Celsius to avoid issues of estimating proportional increases for values close to zero.

#### Species level projections

Using the models that we validated in our forecasting accuracy assessment, we projected the occurrence of each species in each 3 km^2^ grid cell for the historical baseline, as well as for two emissions scenarios, from each of the two regional ocean models. We substituted the in situ temperature, oxygen, and depth measurements from the trawl surveys with outputs from the regional oceanographic models. This substitution requires making the assumption that the in situ measurements are comparable with the model outputs. The model outputs represent mean summer (April to September) conditions over multiple years and so are less variable than the in situ measurements. However, a comparison of outputs from the BCCM model and the in situ measurements shows good agreement, with a correlation of 0.841 for temperature and 0.836 for dissolved oxygen (Figure S6). A similar comparison is not possible for the NEP36 model, because we do not have model outputs from after 2005. For our projections, we set our survey fixed effect to be DFO Pacific as this survey covers the majority of our focal region.

Projected occurrence change for each scenario and model was calculated as the historical projected occurrence subtracted from the future projected occurrence for each grid cell (Figure 1f). For each species, we excluded grid cells where the projected probability of occurrences fall below 0.1 in both the historical and future periods, as these are not considered to be important habitat. This allowed us to exclude areas where the projected change in occurrence is minimal because conditions are unsuitable in both time periods. Our findings remain qualitatively unchanged when we use a threshold of 0.3 (results not shown).

To estimate the degree to which each species could tolerate increases in temperature, decreases in oxygen, and shifts to deeper depths, we estimated a margin for change across all of the trawls in our focal region for which the species was present. This allowed us to assess which aspects of environmental change are most likely to have negative consequences for each species, within our focal region. This margin for change was estimated as the difference between the conditions in which they were observed (i.e., present in a trawl) in B.C. and Washington and a given threshold for each variable. This threshold was the estimated breakpoint for dissolved oxygen and the maximum temperature or depth where our SDM estimates that the probability of occurrence remains above 0.1. However, because the influence of one variable on occurrence probabilities depends also on the other two variables, we estimated these thresholds for temperature and depth based only on combinations of conditions observed in our coastwide training dataset. This ensured that thresholds were not based on unreasonable combinations of variables (e.g., high dissolved oxygen or temperature at depth). We also excluded conditions that were greater than 50 m of the maximum depth and greater than 1 °C of the maximum temperature where the focal species was observed to be present in the coastwide dataset. Once we had estimated the threshold values for each variable, we compared these with the observed conditions in all of the trawls in our focal region (i.e., B.C. and Washington), where the focal species was observed. The difference between the observed conditions, conditional on species presence, and the estimated threshold was then taken as the margin for change in that location and time. We then aggregated the margin for change across all trawls to obtain a distribution for each species, for each environmental variable, for our focal region.

We quantified projection uncertainty due to SDM parameter uncertainty as the average width of the 95% confidence interval for the probability of occurrence. For this we took the average value across all grid cells where the species was projected to be present with at least a probability of 0.1 in the future scenarios, averaged across both regional ocean models and both emissions scenarios. We quantified regional ocean model uncertainty as the absolute difference in projected occurrence in the future scenarios based on the two regional ocean models, averaged across all grid cells where the species were projected to be present with at least a probability of 0.1, and averaged across both emissions scenarios. We quantified scenario uncertainty as the absolute difference in projected occurrence between RCP 4.5 and 8.5, averaged across all grid cells where the species was projected to be present with at least a probability of 0.1, and averaged across both regional ocean models.

#### Community level projections

Projected species richness for each period, regional ocean model, and scenario was calculated as the summed probability of occurrences across all species. Projected species richness change was then calculated as the historical projected richness subtracted from the future projected richness within each 3 km^2^ grid cell.

## Results

### Forecast accuracy

Of the 77 species that we assessed for forecast accuracy, 34 had a Tjur R^2^ greater than our 0.2 threshold for inclusion (Figure S7). All species that met this threshold also had an AUC greater than 0.75. The mean and maximum Tjur R^2^ of the species that met our inclusion threshold was 0.43 and 0.83 respectively (Figure 2b). Models that included a subset of the environmental covariates generally had lower forecast accuracy (Figure S8), with the greatest loss of predictive accuracy occurring with the omission of depth, survey, and temperature. Models that did not include dissolved oxygen had similar forecast accuracy compared to models with dissolved oxygen (Figure S8). However, we opted to keep oxygen in our models because experimental evidence clearly shows that oxygen is an important determinant of species performance (Deutsch *et al.* 2015).

**Figure 2:**
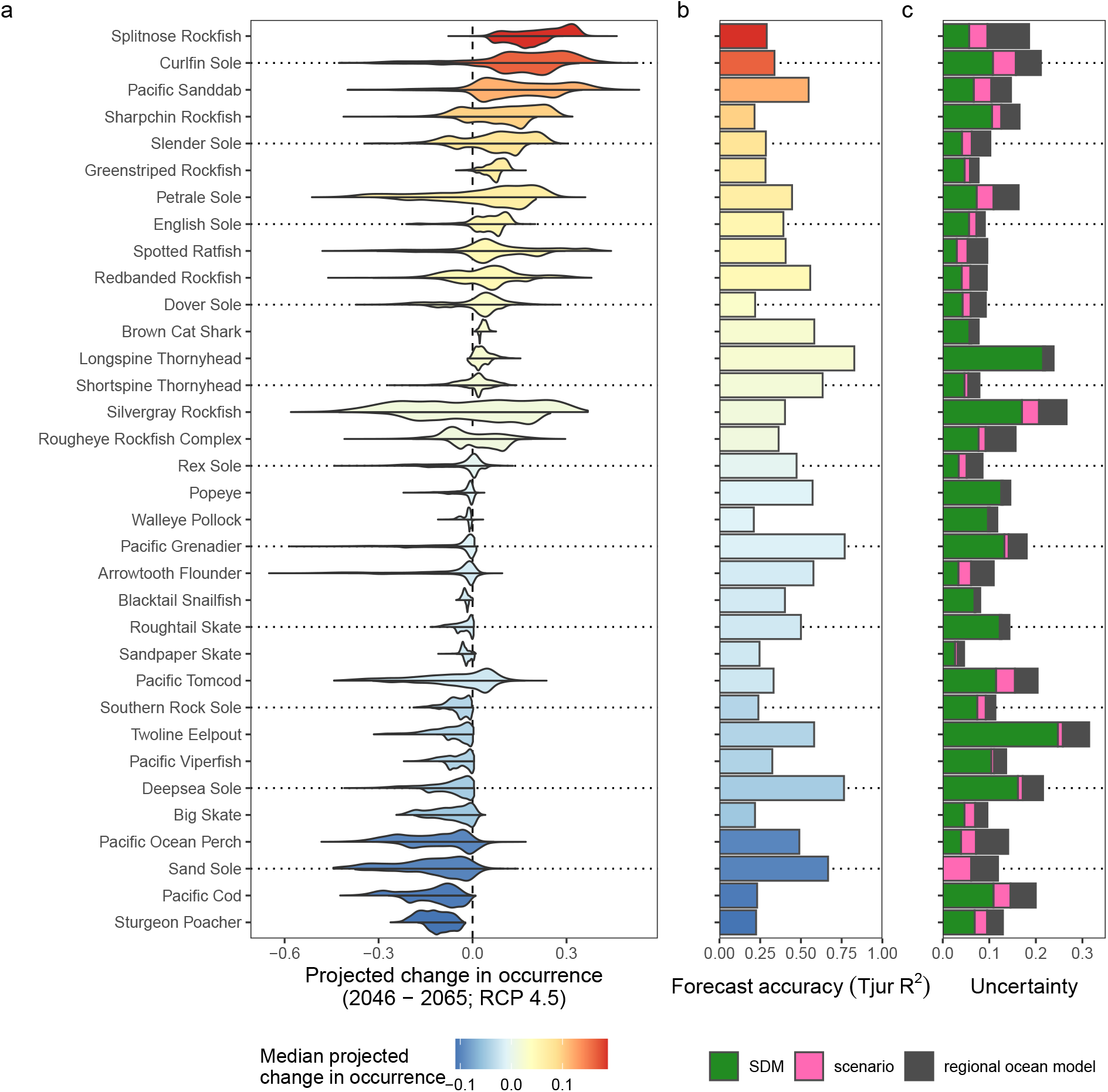
Distributions of projected change in occurrences across the 3 km^2^ grid cells in the study region for all species (a) based on a comparison of average conditions in 1986–2005 vs 2046–2065 under the RCP 4.5 emissions scenario (see Figure S10 for RCP 8.5). For each species, the upper half of the distribution shows projections based on the NEP36 model and the lower half shows projections based on the BCCM model. Grid cells where projected occurrences fall below 0.1 in both the historical and future periods are not considered to be habitat and changes in occurrence in these grid cells are excluded from the distributions. Species are ordered from most positive to most negative median change in projected occurrence. Panel (b) shows the forecast accuracy, measured as Tjur R^2^ for post 2010 data in the focal region using a model trained on all pre 2011 data. The color indicates the median projected change in occurrence across all 3 km^2^ grid cells in the region based on the BCCM RCP 4.5 scenario. Panel (c) shows the average estimated uncertainty of the projections across all 3 km^2^ grid cells where the probability of occurrence is greater than 0.1 in both the historical and future periods, separated by SDM uncertainty (95% confidence interval), emmissions scenario difference, and regional ocean model difference.

### Estimated environmental responses

Occurrence probabilities for all species were estimated to peak at intermediate depths (i.e., negative quadratic depth coefficient), but there was wide variation in the estimated depth ranges (i.e., variation in linear and quadratic depth coefficients; Figure S9). Our models estimated reasonable oxygen break points and slopes for 22 out of the 34 species. The estimated oxygen breakpoints varied from 0.154 to 2.366 mL L^−1^, with a median of 0.802 mL L^−1^ (Figure S9). Occurrence probabilities for all but 4 species were estimated to peak at intermediate temperatures (i.e., negative quadratic temperature coefficient), but there was wide variation in the estimated temperature ranges (i.e., variation in linear and quadratic temperature coefficients; Figure S9). The 4 species that were not estimated to have unimodal temperature responses were all species where occurrences were highest at the lowest temperatures (i.e, lowest linear temperature coefficients).

### Projected environmental change

Both regional ocean models projected increases in bottom temperature across the entire focal region, except for off the shelf at depths greater than 1400 m where increases are projected to be less than 0.01 °C (Figure S4). The NEP36 model projects a greater degree of warming (median 0.98 °C for RCP 4.5) compared to the BCCM (median 0.73 °C for RCP 4.5). Projected changes are greater under RCP 8.5 (median 1.21 °C for NEP36 and 0.92 °C for BCCM). The greatest increases (as high as 2.82 °C under RCP 8.5 in the NEP36 model) are projected in the shallowest waters to the east of Haida Gwaii.

Projected changes in dissolved oxygen differed more between the two regional models (Figure S5). The BCCM model projected oxygen losses in the shallower waters on the continental shelf (< 300 m; median −0.12 mL L^−1^ under RCP 4.5 and −0.16 mL L^−1^ under RCP 8.5), negligible changes in oxygen at mid depths (troughs in Queen Charlotte Sound and on the shelf slope), and increases at the deepest waters off the shelf (> 500 m; median 0.03 mL L^−1^ under RCP 4.5 and 0.03 mL L^−1^ under RCP 8.5). In contrast, the NEP36 model projected decreases across most of the region (median −0.25 mL L^−1^ under RCP 4.5 and −0.27 mL L^−1^ under RCP 8.5; maximum −0.63 mL L^−1^ under RCP 4.5 and −0.72 mL L^−1^ under RCP 8.5).

### Projected species’ occurrence change

Projected occurrence changes for Washington and B.C. varied considerably across species (Figure 2a). These projections were consistent across the two regional ocean models, although the projected changes based on the NEP36 model tended to be slightly larger compared to those based on the BCCM model (Figure 2a). Likewise, the projected changes in species occurrences were similar in models based on the RCP 4.5 and 8.5 emissions scenarios, with RCP 8.5 resulting in projected changes that were of slightly greater magnitude (Figure 2 vs. S10). Under the RCP 4.5 scenario with the BCCM (NEP36) model, 5 (8) species were projected to decrease in occurrence in at least 95% of the region where they were estimated to be present, 3 (3) species were projected to increase in occurrence in at least 95% of the region where they were estimated to be present, while the remaining 26 (23) species were projected to increase and decrease in prevalence in different parts of the region. The projections were similar for the RCP 8.5 scenario with 5 (10) species decreasing in occurrence, 3 (3) species increasing in occurrence, and 26 (22) showing variable changes in projected occurrence.

Quantifiable uncertainty in the probability of occurrence change varied across species, ranging from 0.046 for Sandpaper Skate to 0.545 for Sand Sole, with an average of 0.154 across all species (Figure 2c). Uncertainty in projected occurrences from SDM model parameter uncertainty resulted in the greatest variation in projected occurrences, with an average 95% confidence interval width across all grid cells (with a projected occurrence > 0.1) and species of 0.096. The next greatest source of uncertainty was due to variation in projected occurrences across the two regional ocean models, which had an average difference of 0.038 across all grid cells (with a projected occurrence > 0.1) and species. Emission scenario differences contributed the least uncertainty to our projections, with an averaged difference of 0.021 across all grid cells (with a projected occurrence > 0.1) and species.

### Species’ margin for change

There was considerable variation in how far species in B.C. and Washington waters are from our estimated maximum temperature thresholds (Figure 3a). Species such as Greenstriped Rockfish and English Sole were caught (>99% of trawls) in conditions that are at least 5 °C cooler than their estimated maximum temperature threshold. Others, such as Pacific Viperfish, Roughtail Skate, and Sand Sole were found at temperatures right up to, and sometimes exceeding their maximum temperature threshold. In contrast, groundfish in B.C. and Washington tended to extend all the way to their estimated maximum depth thresholds (Figure 3b; Wall-eye Pollock and Rex sole being notable exceptions). Observed species’ occurrences were common above the estimated oxygen breakpoints, but tended to decrease quickly at lower oxygen levels (Figure 3c). However, some species such as Walleye Pollock, Blacktail Snailfish, and Twoline Eelpout were predominantly found in conditions where oxygen was estimated to be limiting. In general, the species that were projected to increase in occurrence throughout B.C. and Washington are those that were far from their temperature thresholds (Figure 3a). Species that were projected to have mixed responses under future conditions are those that have wide depth ranges (Figure 3b). Species that were projected to decrease in occurrence throughout the region fall into two categories: 1) they are close to their maximum temperature thresholds and occupy a relatively narrow range of depths (e.g., Sand Sole, Sturgeon Poacher), 2) they are already found in conditions with oxygen concentrations near or below their estimated oxygen breakpoint (e.g., Pacific Cod, Pacific Ocean Perch).

**Figure 3:**
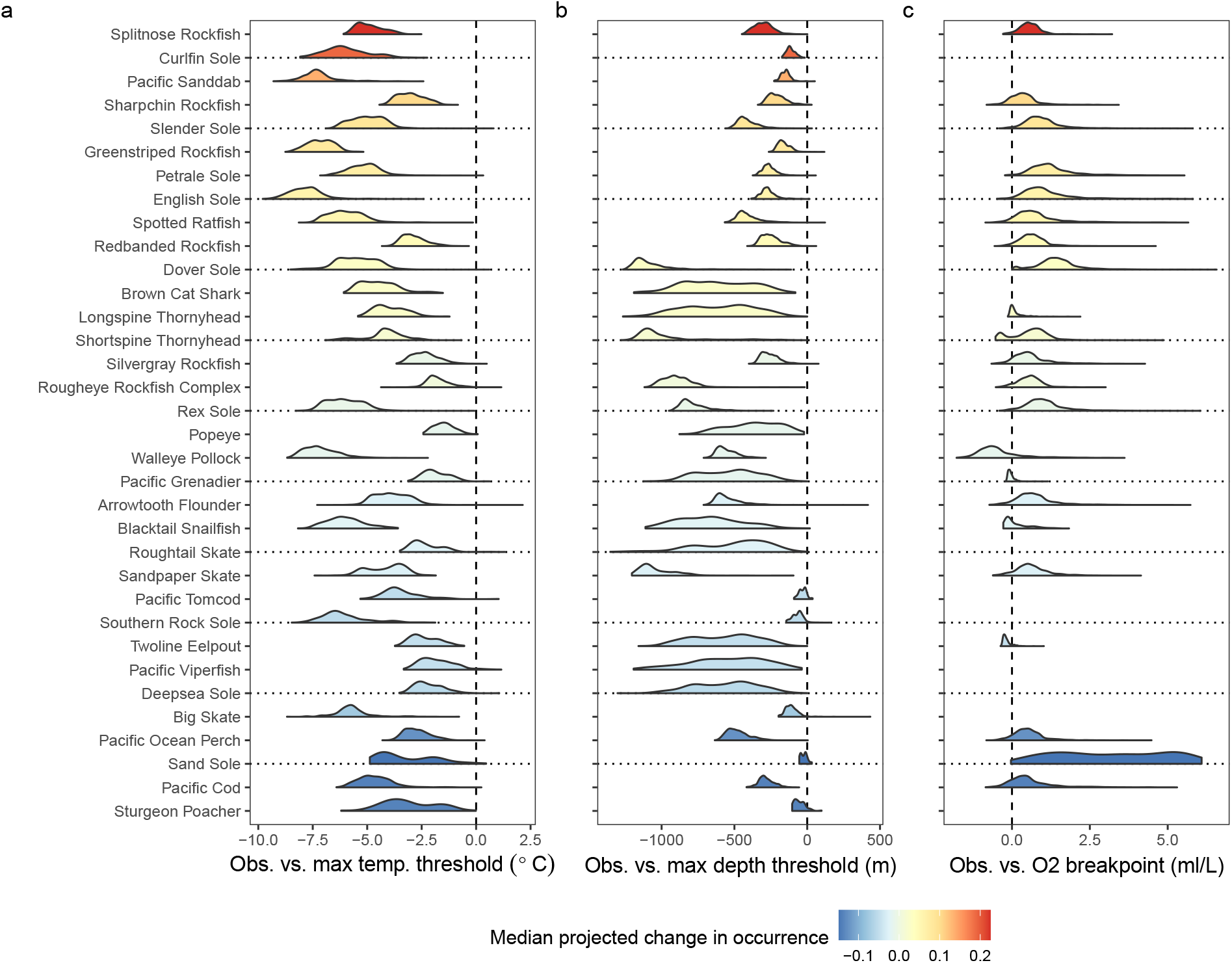
Species’ margin for change in temperature (a), depth (b), and dissolved oxygen (c). Margins were calculated as the observed environmental value in each trawl where a species was present minus the threshold, breakpoint for dissolved oxygen, value for that variable estimated using the SDM (see methods for details). Threshold values are for high temperature and depth and for low dissolved oxygen concentrations. Distributions represent the margins for all trawls where a species was present. The distance between the distribution and the dashed 0 line (left for temperature and depth, right for oxygen) indicates the degree to which the species is estimated to be able to tolerate warmer, deeper, or lower oxygen conditions respectively. Species that do not have distributions for oxygen margin change are those that did not have oxygen included in the model. As for Figure 2, the color indicates the median projected change in occurrence between 1986–2005 vs 2046–2065 for the RCP 4.5 scenario in the BCCM model across all 3 km^2^ grid cells in the region and species are ordered from most positive to most negative change.

#### Projected species richness change

Projected species richness changes varied spatially across our focal region, ranging from −4.2 to 2.6, with a median of 0 across both regional ocean models and both emissions scenarios (Figures 4a,c, S11a,c). These projected richness changes varied strongly with depth (Figures 4b,d, S11b,d). Decreases were projected in the shallowest waters (< ~100 m) where warming is projected to be greatest, and in deep, low oxygen waters (> ~600 m) where further reductions in oxygen are projected to be detrimental for many species. In contrast, species richness is projected to increase at mid depths (~100–600 m) as species shift to deeper depths to deal with warming or shift to shallower depths to avoid hypoxic conditions. Projected changes were greater in magnitude based on the NEP36 model, particularly at depth where the NEP36 model projects oxygen loss while the BCCM projects oxygen increases over this time period (Figure S5). Projected richness changes were similar, but of great magnitude, for the RCP 8.5 emissions scenarios (Figure S11).

**Figure 4:**
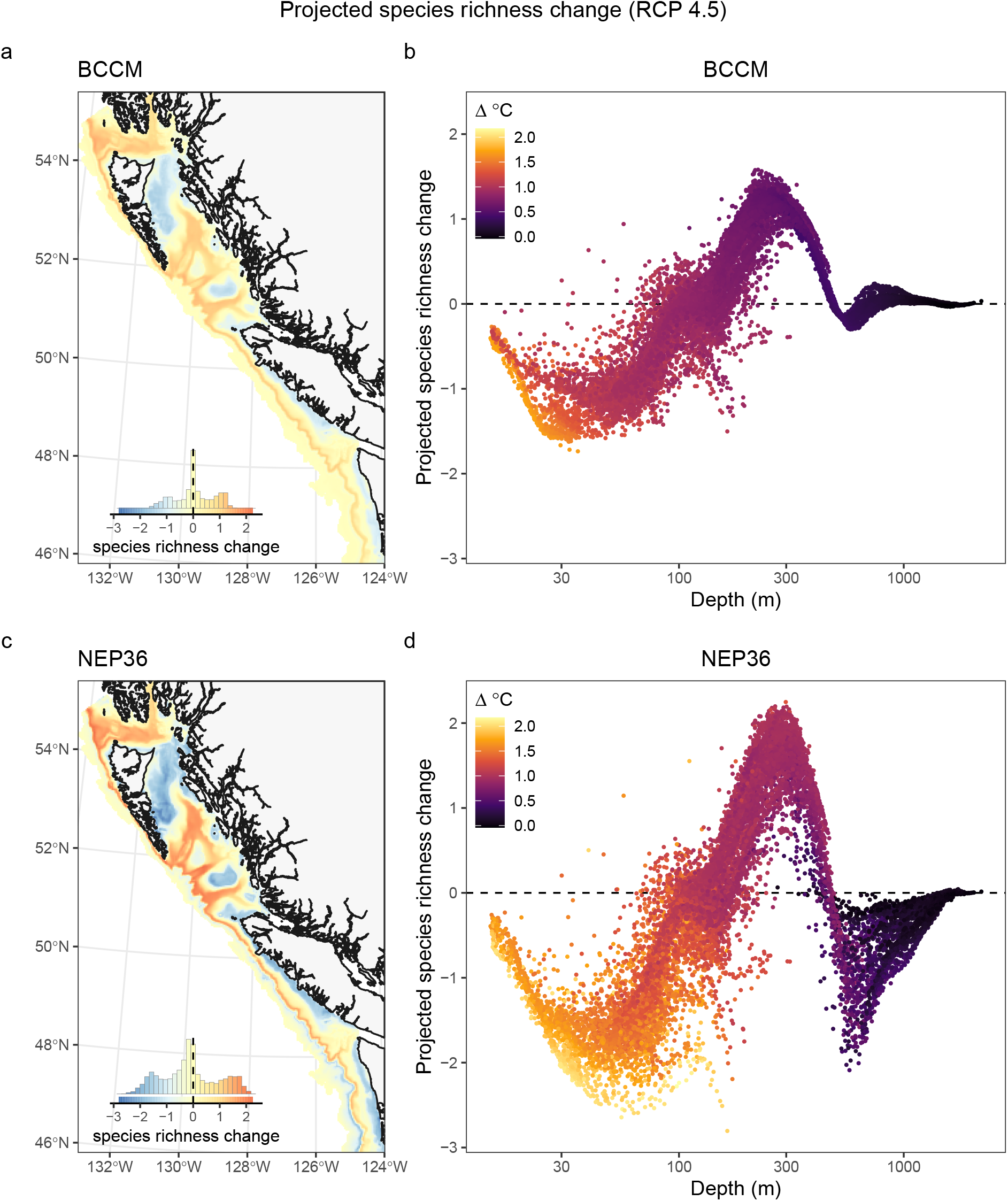
Projected species richness change (a, c) mapped across the study region between historical (1986–2005) and future (2046–2065, RCP 4.5) and as a function of seafloor depth (b, d). Panels (a, b) show projections based on the BCCM model. Panels c and d show projections based on the NEP36 model. The inset histograms in panels (a, c) show the distribution of values across all 3 km^2^ grid cells and provide a legend for the colours shown on the maps. Each point in panels (b, d) represents a single 3 km^2^ grid cell in panels (a, c) coloured by the projected temperature change. See Figure S11 for results based on RCP 8.5.

## Discussion

Our analysis suggests that projected warming and deoxygenation is likely to cause a reorganization of the groundfish community in B.C. and Washington by 2046 to 2065. We estimate that species will differ in their responses to climate change; some species will benefit from warmer conditions and increase in occurrence, others will shift to deeper waters as conditions warm, but those that cannot tolerate deeper depths or lower oxygen are expected to decrease in occurrence. While species that are projected to decline are roughly balanced by species that are projected to increase, the overall impact on the community is expected to depend on depth. At depths of less than 100 m where warming is projected to be greatest, we expect to see a decline in species richness. Species richness is also projected to decline at depths below 600 m because of oxygen loss. However, there is uncertainty in how much richness will decline due to oxygen loss as our two regional ocean models differ in their projections of oxygen change at these depths. At mid depths, between 100 and 600 m, we expect species richness to increase as warming drives shallow species deeper and deoxygenation in deeper waters pushes species shallower. The increase in species richness is projected to be greatest at depths around 300 m as below this hypoxic conditions (Diaz & Rosenberg 2008) may prevent further species range shifts. While the magnitude of these projected changes to the groundfish community varies depending on the regional ocean model and the emissions scenario used, the overall pattern of change is generally consistent in all cases. This result suggests that our projections for how the groundfish community will reorganize should be qualitatively robust to uncertainty in how climate change will play out over the coming decades.

These findings demonstrate the importance of accounting for the influence of depth and dissolved oxygen when projecting how climate change will affect species in marine environments. Based on our projections, it is not possible to determine how a species will respond to warming based only on its temperature response curve because species vary in the degree to which their tolerance to depth and low oxygen constrain their ability to shift to deeper depths as conditions warm (Figure 3). Therefore, SDMs which do not account for depth and dissolved oxygen (e.g., Morley *et al.* 2018) may underestimate the sensitivity of marine organisms to climate change because they assume that species can always shift to deeper waters when conditions warm. In contrast, models that rely only on sea surface temperatures (e.g., García Molinos *et al.* 2016) assume that species cannot shift in depth to deal with warming, thus they are likely to overestimate the sensitivity of groundfish to climate change. The fact that dissolved oxygen and depth were correlated, even in our coastwide dataset, posed a challenge for accurately estimating the oxygen responses in our models. For a subset of species, our models estimated that occurrence decreased with increasing oxygen, which is unlikely given the fact that oxygen is understood to be a limiting factor (Fry 1971). This indicates that, for these species, the models are erroneously attributing the fact that the species is not found in shallower oxygenated waters to oxygen rather than some other variable that is correlated with depth. Indeed, models without oxygen responses tended to have similar forecast accuracy as those with oxygen responses, indicating that the depth response is able to capture the oxygen response, at least over the time period where we have trawl data. Nevertheless, because oxygen limitation is understood to have strong and detrimental impacts on fish performance (Fry 1971; Deutsch *et al.* 2015), we elected to keep oxygen in our SDMs whenever it was possible to estimate a reasonable parameters for the oxygen response.

Of course, additional variables that we have not accounted for may further constrain species distributions as well as their responses to warming and oxygenation. For example, projected ocean acidification (Peña *et al.* 2019; Holdsworth *et al.* 2021) may impact fish populations (Branch *et al.* 2013; Haigh *et al.* 2015), but SDMs are not well suited to projecting these because we lack occurrence data across relevant pH ranges. Species distributions may also be constrained to certain seafloor substrate types (Pirtle *et al.* 2019) and including substrate in our models may result in more accurate projections (e.g., Morley *et al.* 2018). However, the lack of comprehensive substrate data for the entire extent of the groundfish surveys used in our analysis precluded us from including this variable in our models. It is likely that the inclusion of additional variables would have improved model performance, and may have increased the number of species that met our inclusion criteria based on the temporal forecasting assessment. However, many of the unobserved variables also change with depth so, as implemented here, depth may be a proxy for these processes. A previous study of groundfish in B.C. estimated that the addition of oceanographic variables, including substrate, currents, and primary production, to a model that already included depth only resulted in a minor increase in the accuracy of predictions (Thompson *et al.* 2022a).

By combining spatially extensive scientific survey data with regional ocean models, we were able to develop robust models that are informed by broad spatial and environmental gradients but make projections at a relatively high resolution. If we had trained our SDMs on data only from our focal region we would not be including the full range of environmental conditions that define the distributions of these species. Furthermore, the high correlation between depth, temperature, and oxygen in B.C. and Washington would have prevented us from distinguishing their unique influence on species’ distributions (Thompson *et al.* 2022a). While some species in our models have ranges that extend beyond the broad spatial extent and environmental range sampled by the surveys, we expect that this had negligible impact on our projections because our focal region is situated in the middle of the geographic and environmental space sampled by the surveys. Reassuringly, the future environmental conditions that are projected by our regional models fall within the range of conditions present in our survey data, except in the few warmest grid cells where future conditions are projected to exceed our observational range by less than 1 °C (Figure S12). Thus, we are largely avoiding extrapolating into novel climate conditions, which is a key source of uncertainty in many SDM projections (Veloz *et al.* 2012). In contrast, if we had extended our projections to cover the full spatial extent of the survey data we would have had to use low resolution outputs from global climate models, which have not been downscaled to incorporate the complex bathymetry of the continental shelf (Hewitson & Crane 1996; Giorgi *et al.* 2009). Alternatively, we could have stitched together multiple regional ocean models (e.g., Veneziani *et al.* 2009; Hermann *et al.* 2019; Kearney *et al.* 2020) but this would have required additional assumptions because models differ in their underlying mechanics, data, and assumptions. As our goal was to make projections that could be used for ongoing management initiatives in Canadian Pacific waters, this approach was not necessary.

Projecting biodiversity responses to climate change involves considerable uncertainty (Urban *et al.* 2016) and our approach allows us to quantify some aspects of this. Of the uncertainty that we could quantify, roughly half was due to uncertainty in the parameters in our SDMs and the remainder was due to regional ocean model uncertainty or scenario uncertainty. This amount of uncertainty in the SDM parameters is typical (Brodie *et al.* 2022), stemming from the fact that contemporary species distributions are also influenced by other factors that we have not included in our model. In addition, although oxygen demand is understood to vary with temperature (Deutsch *et al.* 2015), limitations in the implementation of breakpoint models prevented us from estimating a temperature dependent oxygen breakpoint. However, although somewhat unrealistic, this limitation is unlikely to have greatly increased the uncertainty in our SDM parameters because low oxygen concentrations occurred almost exclusively at depths where temperature variation and projected change was small.

To reduce uncertainty due to year-to-year variation in climate, our model projections are based on 20-year climatologies with a future period that is far enough in the future to ensure that changes are unambiguously due to greenhouse gases (Hawkins & Sutton 2009; Holdsworth *et al.* 2021). We have made projections based on two different emissions scenarios, and two different regional ocean models that are both downscaled from the same global model, the second generation Canadian Earth System model (CanESM2), using different downscaling techniques. While the BCCM model was run inter-annually and then averaged to produce the climatologies (Peña *et al.* 2019), the NEP36 model used atmospheric climatologies with augmented winds to force the ocean model and produce representative climatologies (Holdsworth *et al.* 2021). Comparing these regional projections provides an estimate of the uncertainty across different regional downscaling models and methods. We find that the projected impacts of climate change on the groundfish community are more sensitive to the differences in the regional ocean models than they are to the emissions scenarios used. However, these differences are in magnitude (changes tend to be larger based on NEP36 compared to the BCCM) rather than in direction, with both models resulting in similar overall patterns of biodiversity change and turnover for the groundfish community. Over the sixty year time period (1986–2005 vs. 2046–2065) used in our study, our projections suggest that groundfish community changes are similar regardless of the scenario used which is not surprising given the fact that the two emissions scenarios do not appreciably differ until later in the 21st century (Arora *et al.* 2011). Ideally, we would also quantify uncertainty due to differences between global climate models and use the ensembles from each of the models (e.g., Frölicher *et al.* 2016; Morley *et al.* 2018), however, regional models driven by other global models have not yet been developed for this region. We expect that projections based on different global climate models would show greater variation than we see between our regional ocean models, but they would produce similar overall patterns in projected groundfish community change unless they project vastly different temperature and oxygen changes.

One aspect of projection uncertainty that we cannot quantify is that which is due to eco-evolutionary processes (Urban *et al.* 2016). Interactions between species, movement across the seafloor (e.g., dispersal, migration, foraging Guzman *et al.* 2019), and evolution all influence current species distributions as well as the way that species will respond when conditions change (Urban *et al.* 2012; Norberg *et al.* 2012; Thompson & Fronhofer 2019). However, SDMs implicitly, and unrealistically, assume that the influence of interactions between species will not change as species shift their distributions and respond to future environmental conditions, that species are not dispersal limited, and that they will not adapt or acclimatize to environmental conditions. Our projections suggest that warming and deoxygenation will cause species to shift their depth ranges, causing more overlap in ranges at mid depths. How this increase in species richness at these depths (and the reductions of species richness in shallower and deeper waters) alters food web dynamics is a major source of uncertainty (Thompson & Gonzalez 2017; Bartley *et al.* 2019). For example, our projections assume that prey exist in sufficient abundance at those new depths, and that the species will be able to establish in the face of novel competitors and predators (Gilman *et al.* 2010). Likewise, any degree of acclimation or adaption to environmental change would cause us to overestimate changes in areas where occurrences are projected to decrease (Aitken *et al.* 2008). Despite these unavoidable sources of uncertainty, we can use our understanding of eco-evolutionary community dynamics to identify areas where our projections are likely to have more and less uncertainty. That is, eco-evolutionary dynamics lead to the greatest uncertainty at range edges where they influence whether an established species will be lost as well as whether a new species can establish (Brown & Thatje 2014; Alexander *et al.* 2015; Urban *et al.* 2016). Conversely, eco-evolutionary processes should result in less uncertainty in areas that are projected to remain as suitable habitat for a species. These areas that are projected to remain suitable offer a “no regrets” conservation opportunity as their protection is likely to benefit the species, both now and in the future.

The species in our analysis are all under some degree of fishing pressure—either as the target of fisheries or as bycatch (Anderson *et al.* 2019). Climate change will make the management of these species more challenging and is likely to threaten the sustainability of some fisheries (Duplisea *et al.* 2021). Estimated species’ sensitivity to climate change is a key knowledge gap in current groundfish stock assessments in this region. While changes in biomass are likely to differ from the changes in occurrence that we have projected here (Thompson *et al.* 2022a), we expect occurrence changes to be informative of the sensitivity of species to climate change. Thus, our results can be used to integrate climate change risk into stock assessments and ecosystem-based fisheries management in this region (Duplisea *et al.* 2021; Roux *et al.* 2022). For many species, this may require accounting for climate driven shifts in species distributions and biomass across international boundaries (Palacios-Abrantes *et al.* 2022).

With this analysis, we have demonstrated how spatially extensive scientific survey data can be combined with high resolution regional ocean models to provide projections of groundfish community changes at a scale that is relevant to developing spatial management strategies to preserve marine biodiversity and sustainable fisheries. Furthermore, we have demonstrated that our overall projected patterns of biodiversity change are consistent regardless of the level of emissions, and across the two different regional ocean models that are available for our region. However, had we chosen a more distant future time period, the sensitivity to emission scenario would have been greater. These projections can inform ongoing management initiatives to ensure that they account for anticipated impacts of climate change. For example, Canada is in the process of establishing a network of marine protected areas (MPAs) and refuges to provide long-term conservation of biodiversity (Environment and Climate Change Canada 2021). While climate change adaptation principles are largely lacking from Canadian MPA management plans to date (O’Regan *et al.* 2021), our projections offer an opportunity to ensure that new protected areas in B.C. are situated in areas that will benefit groundfish and other species, both now and under future climates.

## Acknowledgements

Thanks to all participants in the DFO Future Habitats working group and associated workshop for feedback and discussions that contributed to this manuscript. We also thank the DFO and NOAA groundfish research trawl survey teams and DFO and NOAA for making this data available. Special thanks to Jerry Hoff at the Alaska Fisheries Science Center for implementing O2 collections on the bottom trawl surveys and providing this data. We thank the Canadian Centre for Climate Modelling and Analysis for CanESM and CanRCM data, the data originators who contributed data to the gridded data products used to define open ocean boundary conditions for the regional model simulations, and the teams who assembled those products. Thank you to the IPHC for providing oxygen data that was used to fit the oxygen model.

## Data availability

Groundfish Survey Data Data from the DFO Groundfish Synoptic Bottom Trawl Surveys are available at: https://open.canada.ca/data/en/dataset/a278d1af-d567-4964-a109-ae1e84cbd24a

Data from the NOAA U.S. West Coast Groundfish Bottom Trawl Survey are available at: https://www.webapps.nwfsc.noaa.gov/data/map

Data from the NOAA Alaska Groundfish Bottom Trawl Surveys are available at: https://www.fisheries.noaa.gov/alaska/commercial-fishing/alaska-groundfish-bottom-trawl-survey-data

Regional Ocean Model Data Monthly averages for the NEP36 model are available at: https://open.canada.ca/data/en/dataset/a203a06d-9c1f-4bb1-a908-fc52912ff658

Historical climatologies from the MCCM are available at: https://open.canada.ca/data/en/dataset/1084a522-c2dd-47a0-81ac-d4810ce29056

Code All code used for this analysis is available at: https://gitlab.com/dfo-msea/cc_sensitivity_predict

## Supplemental Methods

### Dissolved Oxygen Model

We opted to use a phenomenological model of dissolved oxygen rather than use values from an oceanographic model because we could not find an oceanographic model that spanned the entire spatial extent of the trawl dataset at a sufficient spatial resolution to match to the missing observations. Instead, we estimated the missing oxygen observations using a Generalized Linear Mixed Effects Models fit with the sdmTMB package (Anderson *et al.* 2022b). This model predicted oxygen as a function of near bottom log pressure, temperature, exponential potential density (*σ_θ_*), and month. A spatial random field was also used to capture spatial variation that was not associated with these fixed effects (details below). Potential density (*σ_θ_*) was calculated from salinity and temperature values using the gsw R package (Kelley *et al.* 2021) with a reference pressure of 0 dbar. Near bottom measurements of salinity, temperature, and pressure were sourced from ocean profile data that were collected during trawl and longline fisheries surveys along the Canadian and US west coasts. Data were sourced from the International Pacific Halibut Commission (IPHC) ocean profile data (Sadorus *et al.* 2016) from 2009 to 2015 and all groundfish trawls where oxygen was measured. This resulted in a final dataset of 12581 observations.

We modelled log dissolved oxygen using a Gaussian observation model:

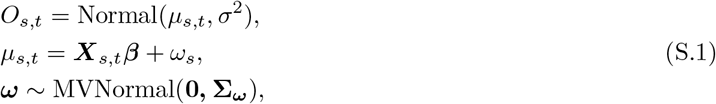

where *O_s,t_* is log dissolved oxygen at location *s* in year *t*, *μ* represents the mean, and *σ* represents the observation error standard deviation. The parameter *X_s,t_* represents the vector of predictors (temperature, pressure, potential density, month) and *β* represents the corresponding vector of coefficients. The parameter *ω_s_* represents the spatial random effects. These were assumed to be drawn from Gaussian Markov random fields with a covariance matrix Σ_*ω*_ that was constrained by Matérn covariance functions (Cressie & Wikle 2011). Oxygen solubility is influenced by temperature and pressure and a non-linear correlation between dissolved oxygen concentration and potential density has been documented in this region (Stefánsson & Richards 1964). Thus we included each of these variables as predictors using plate regression splines. Dissolved oxygen is also understood to fluctuate over the course of the year and so we included month as a predictor using a cyclic cubic spline. The spatial component of the random fields was modelled using a triangulated mesh with vertices selected based on a maximum distance threshold of 15km cut off. We trained the dissolved oxygen model on 70% of the observations and tested it on the remaining 30% hold out data.

## Supplemental Figures

**Figure S1:**
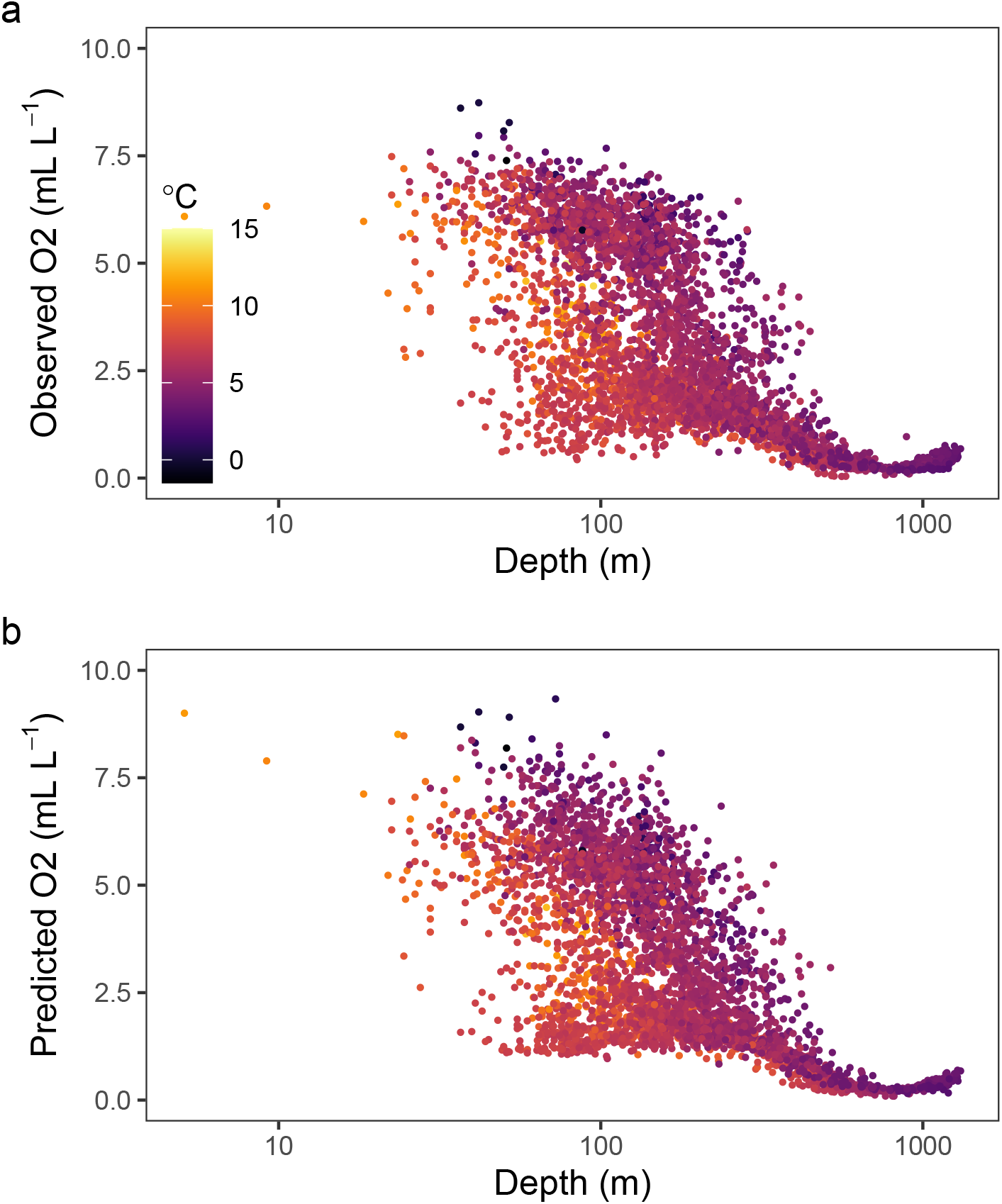
Observed (a) and predicted (b) dissolved oxygen as a function of depth in data withheld from the model.

**Figure S2:**
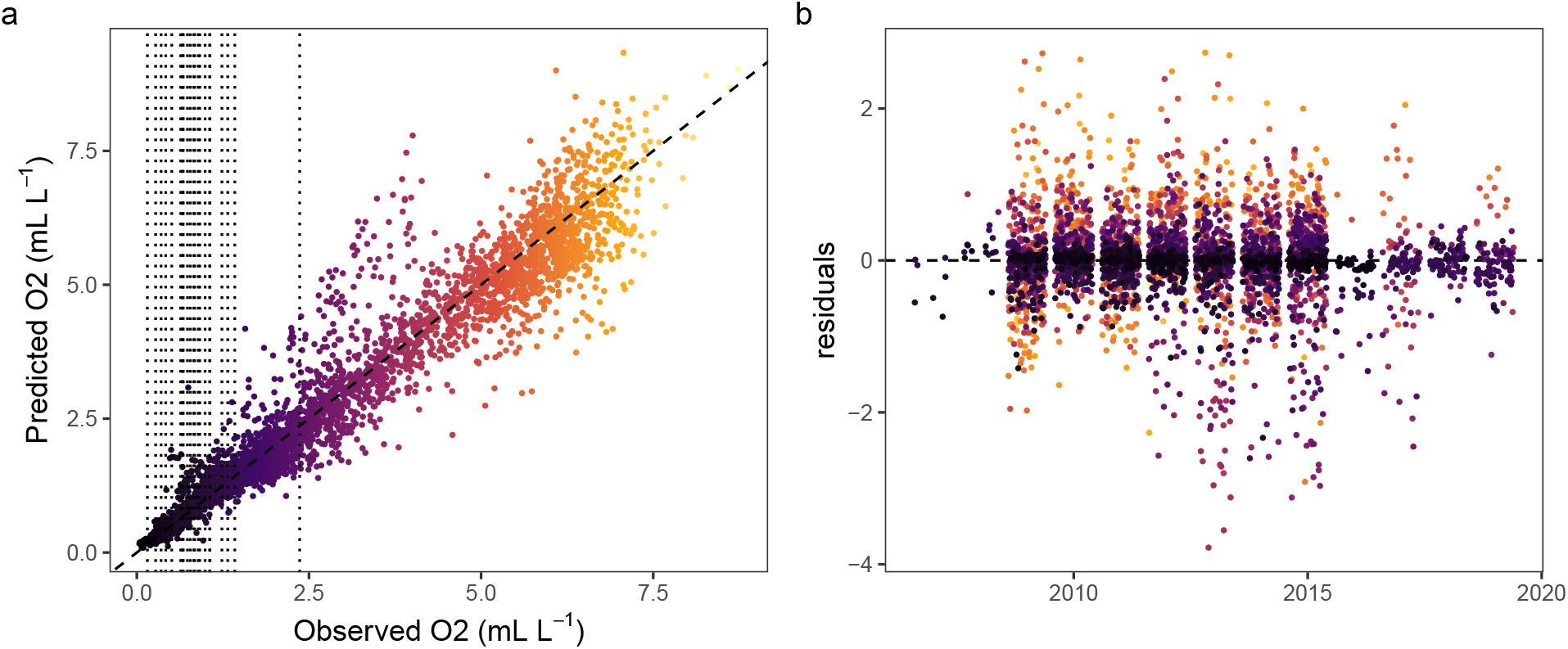
Comparison of observed vs. predicted dissolved oxygen concentrations for data that was witheld from the model. Panel (a) shows a scatterplot of the observed vs. predicted values. The vertical dotted lines show the estimated oxygen breakpoints for each species to show the range of oxygen values where oxygen is estimated to limit species’ occurrence. Panel b shows the residuals (observed - predicted) by year. Points in both panels are coloured by their observed dissolved oxygen concentration with the x-axis of panel a acting as a colour legend.

**Figure S3:**
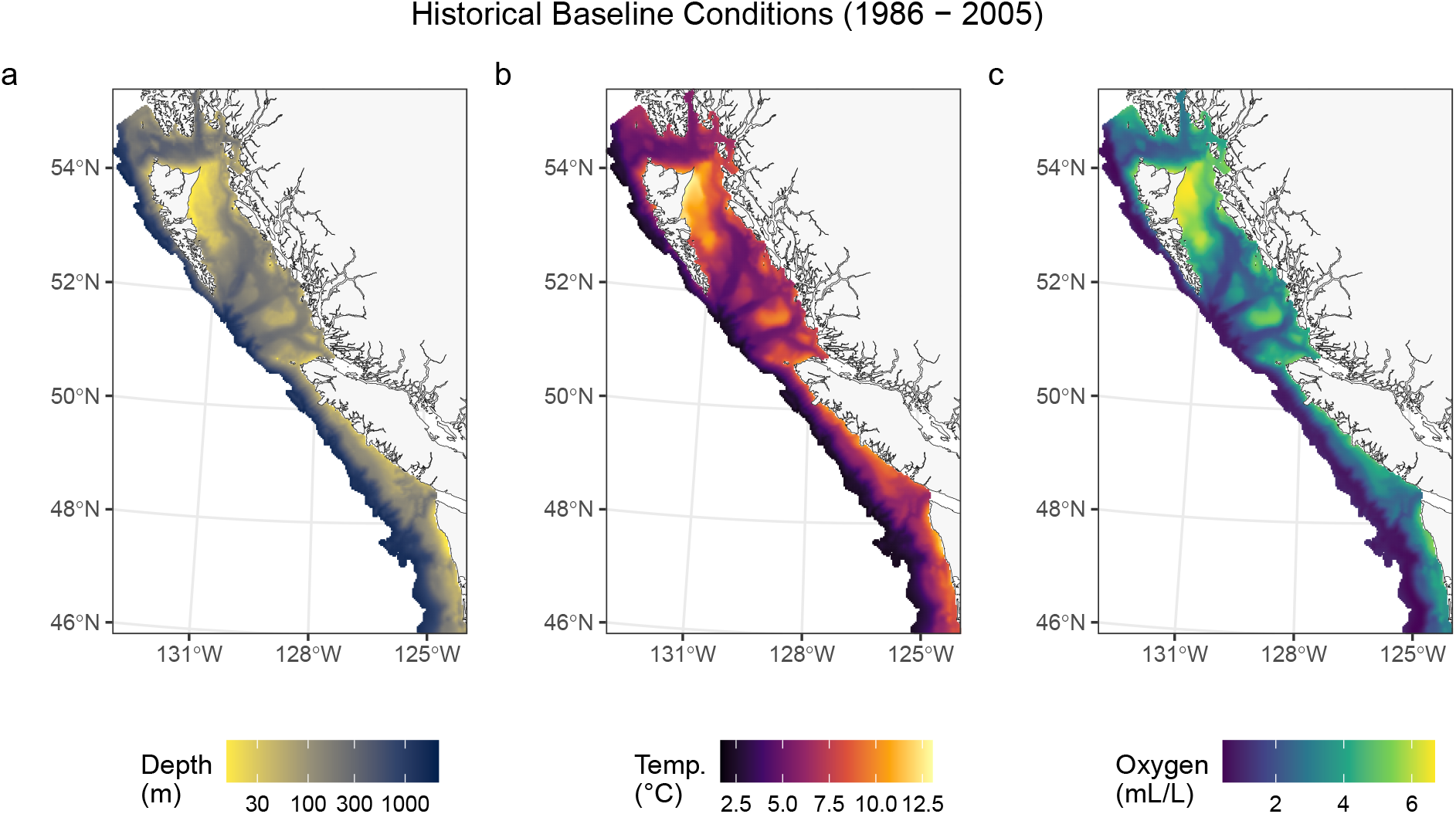
Maps of the historical baseline conditions (1986 - 2005) for seafloor depth (a), near seafloor temperature (b) and near seafloor dissolved oxygen (c).

**Figure S4:**
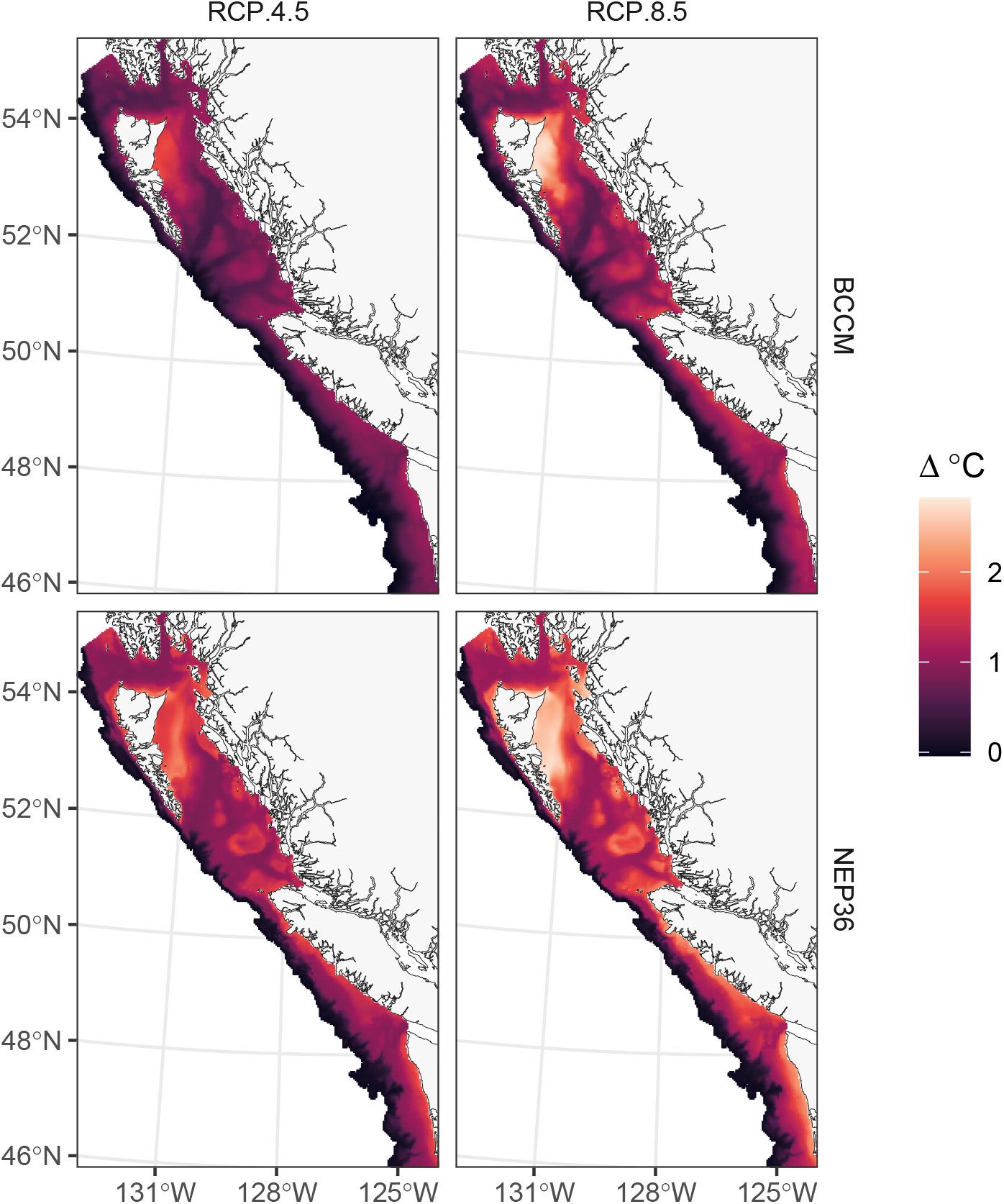
Maps of near seafloor temperature change in the two regional ocean models (rows) and the two RCP scenarios (columns) from the 1986–2006 baseline to 2046–2065.

**Figure S5:**
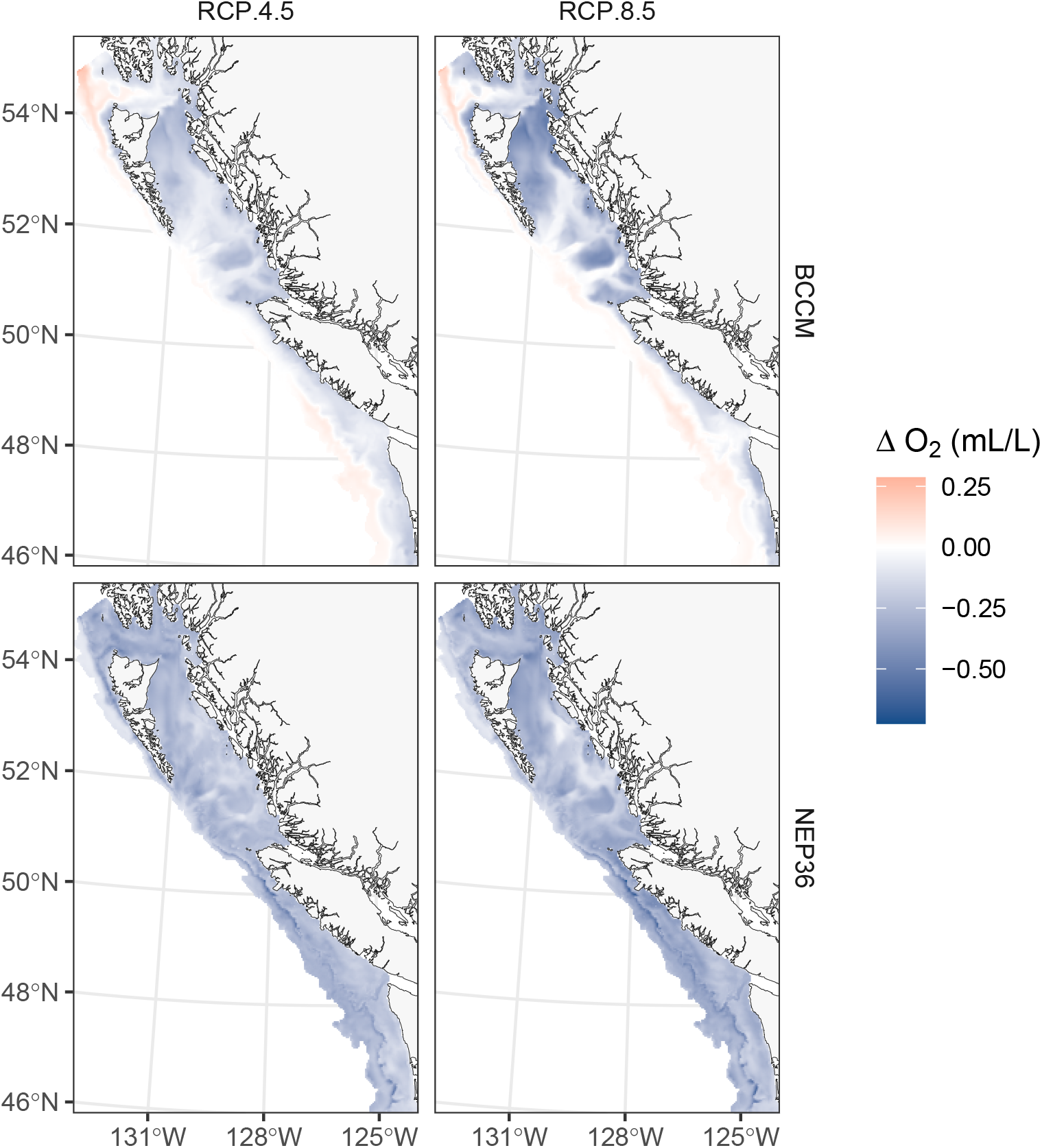
Maps of near seafloor oxygen change in the two regional ocean models (rows) and the two RCP scenarios (columns) from the 1986–2006 baseline to 2046–2065.

**Figure S6:**
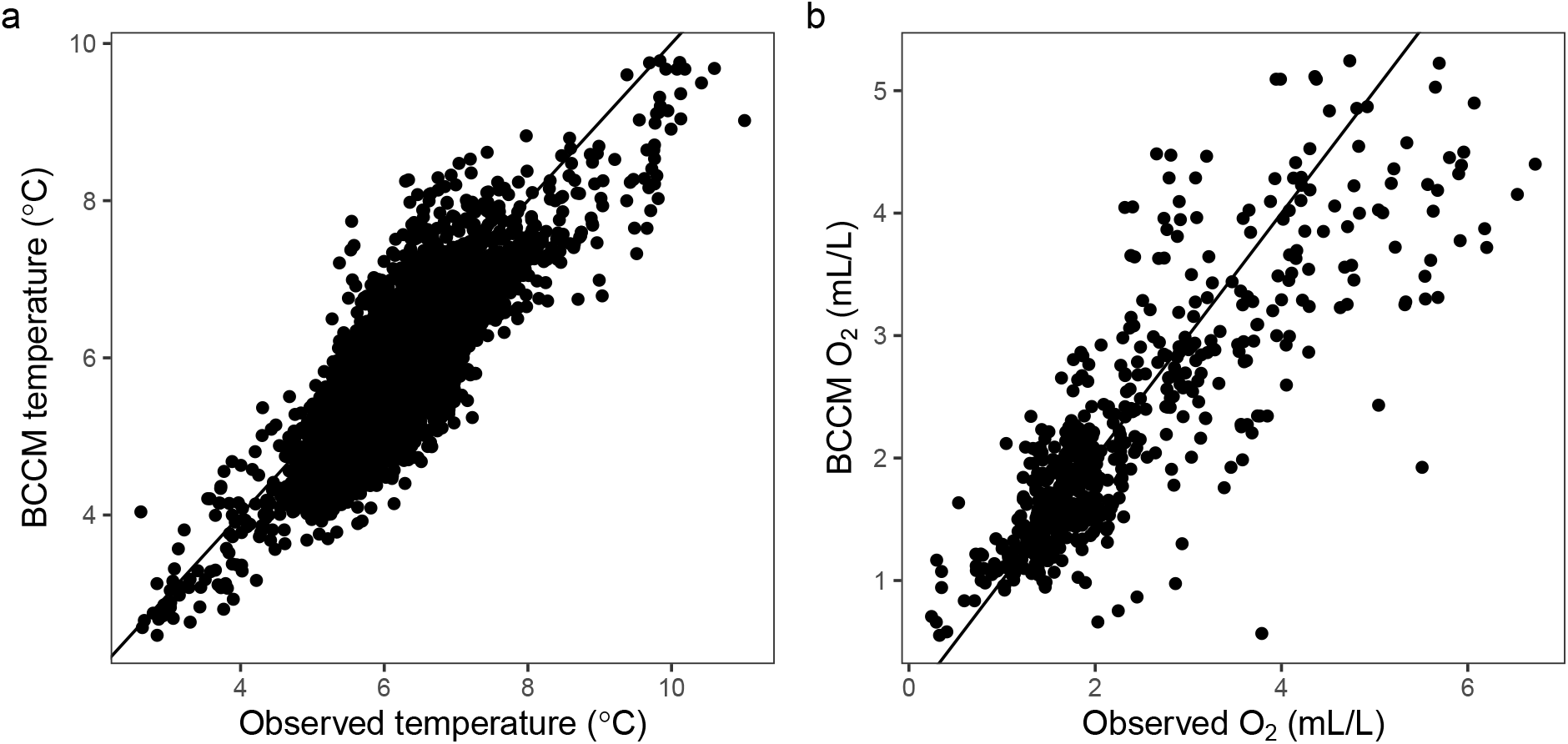
Comparison of observed and modelled temperature (a) and dissolved oxygen (b). Observations are from sensors on the trawls, modelled data are yearly mean summer (April to September) outputs from the BCCM. Data points are matched based on the year in which the trawl was taken whether the center of the trawl fell within the 3 km^2^ grid of the BCCM model.

**Figure S7:**
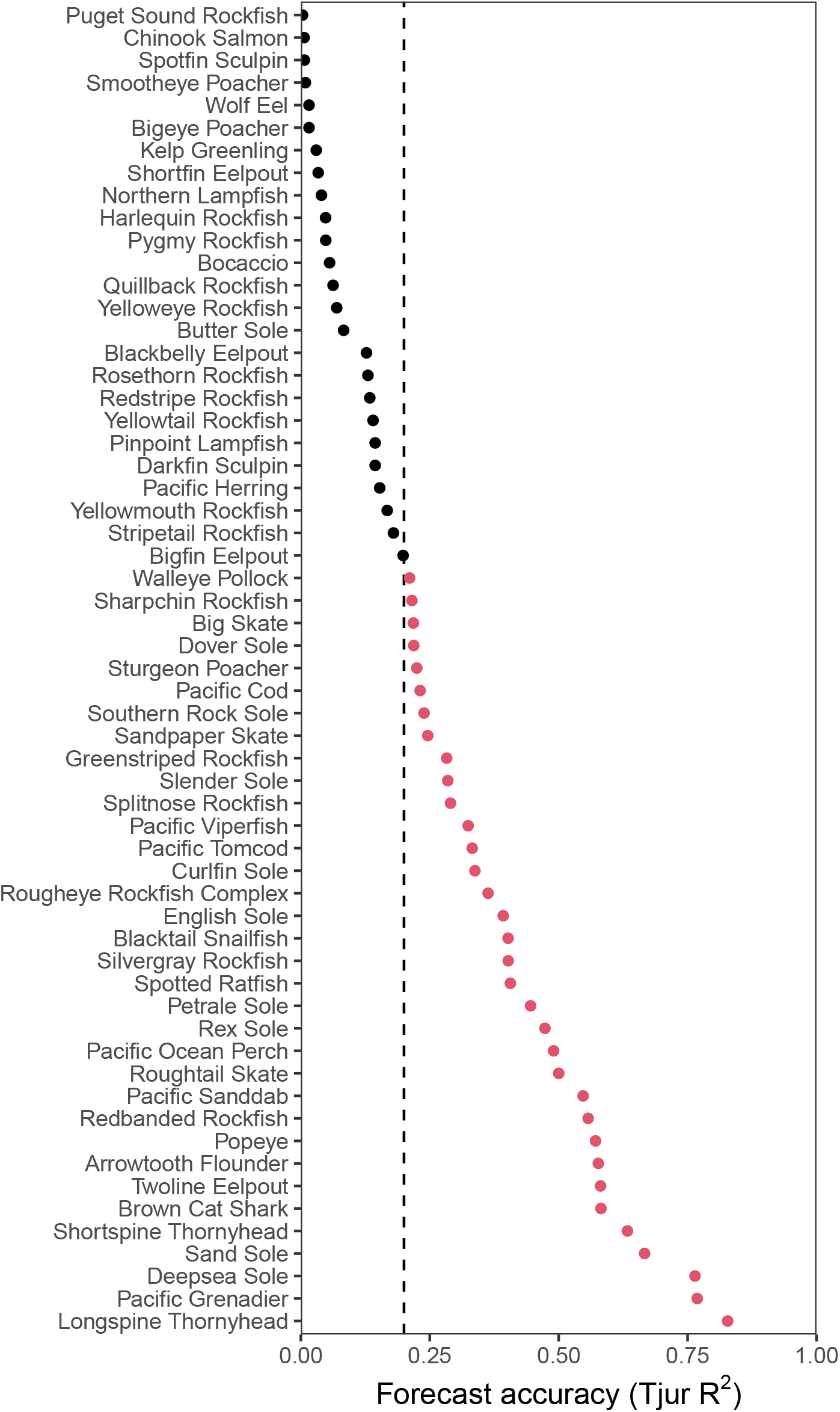
Model forecast accuracy assessment showing the Tjur R^2^ for each species for predictions made for post 2010 data research trawl data from B.C. and Washington based on a model trained on coastwide data prior to 2011. The vertical dashed line shows the 0.2 Tjur R^2^ cut off for inclusion in the subsequent analysis. Species that meet this threshold are shown in red, species that do not are shown in black. Only species with an forecasting accuracy area under the curve (AUC) of 0.75 or greater are shown.

**Figure S8:**
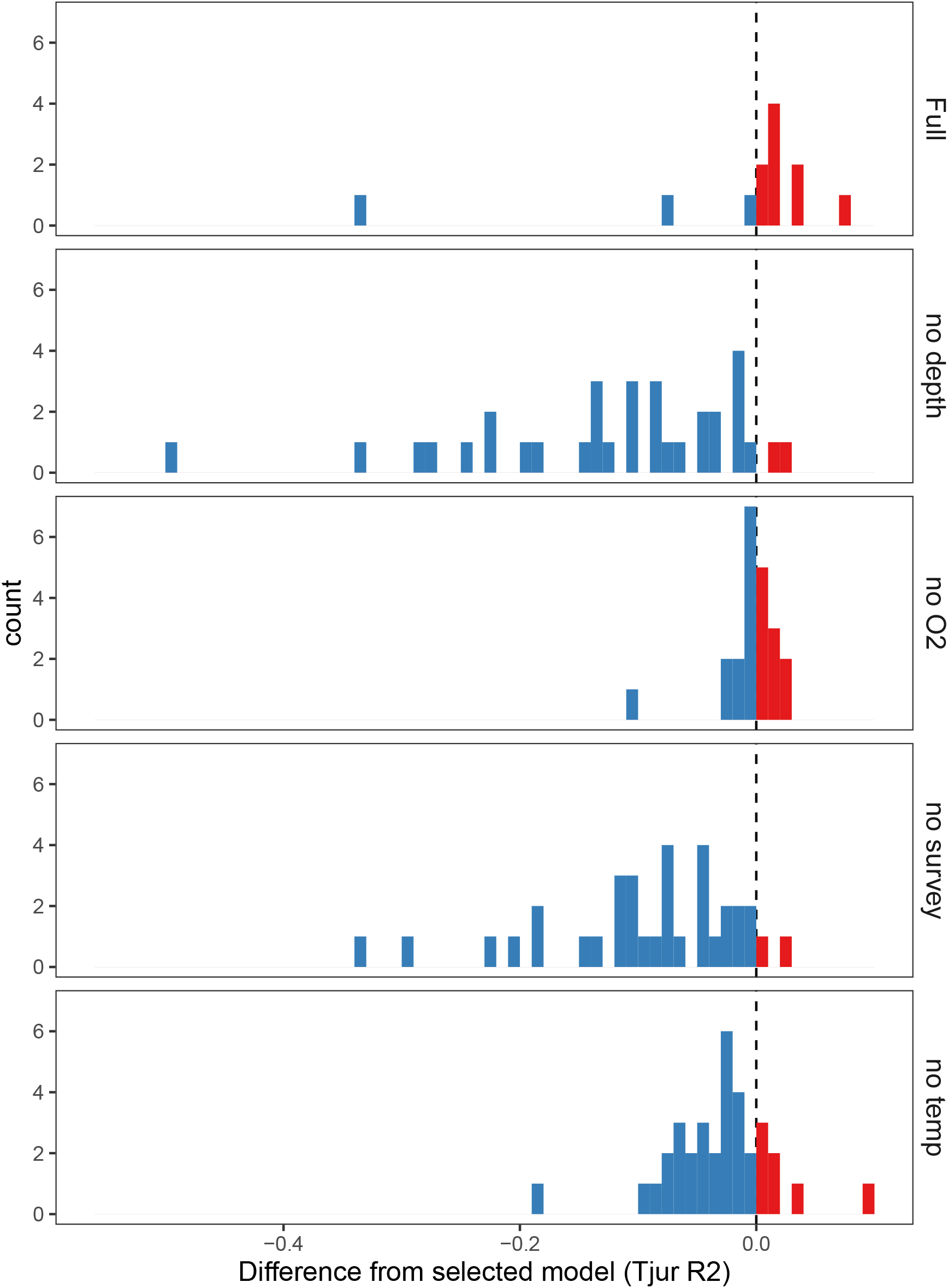
Comparison of models with different subsets of the fixed effects with our final selected model for each species based on forecast accuracy (Tjur’s R^2^). Negative values (blue) indicate that the model had lower forecast accuracy compared to the selected model, positive values (red) indicate that the model had higher forecast accuracy compared to the selected model. Each species contributes a single count in each histogram. Species shown in the full model panel are those for which the full model resulted in unreasonable oxygen response estimates and so the selected model was the model without salinity.

**Figure S9:**
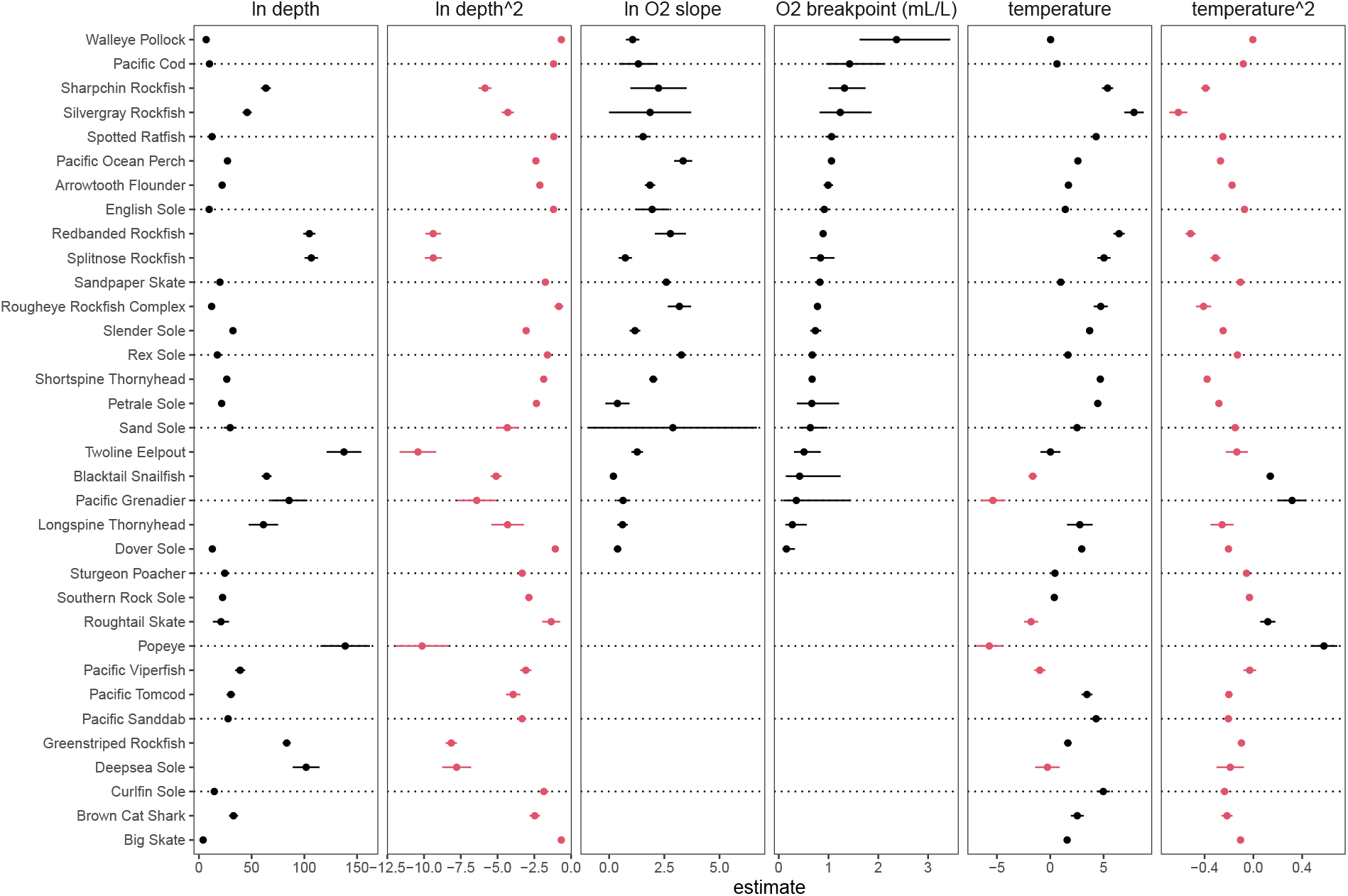
Estimated model coefficients and standard errors for depth, dissolved oxygen, and temperature. Depth and oxygen slope coefficients are based on log transformed variables but the oxygen breakpoint has been transformed so that it is expressed as mL/L. Postive coefficients are coloured black, negative coefficients are coloured red. The error bars show the 95% confidence interval. Species are ordered based on the estimated oxygen breakpoint. Species for which a reasonable oxygen breakpoint could not be estimated were modelled without oxygen, as indicated by the lack of estimated coefficients for oxygen.

**Figure S10:**
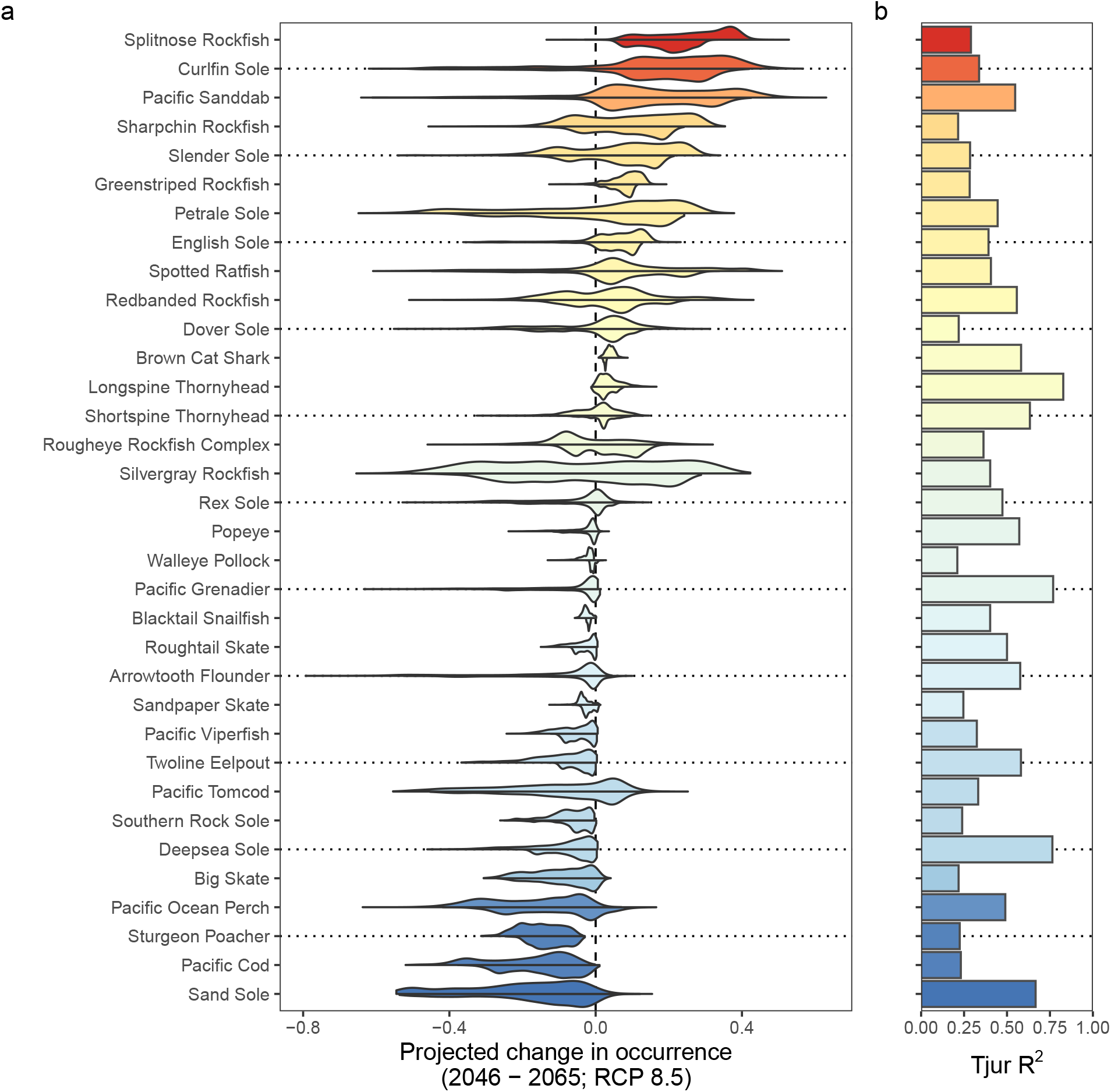
Distributions of projected change in occurrences across the 3 km^2^ grid cells in the study region for all species (a) based on a comparison of average conditions in 1986–2005 vs 2046–2065 under the RCP 8.5 emissions scenario. For each species, the upper half of the distribution shows projections based on the NEP36 model and the lower half shows projections based on the BCCM. Grid cells where projected occurrences fall below 0.1 in both the historical and future periods are not considered to be core habitat and changes in occurrence in these grid cells are excluded from the distributions. Panel (b) shows the Tjur R^2^ predictive power based on the temporal forecasting assessment. The color indicates the median projected change in occurrence across all 3 km^2^ grid cells in the region based on the BCCM RCP 8.5 scenario. Species are ordered from most positive to most negative change.

**Figure S11:**
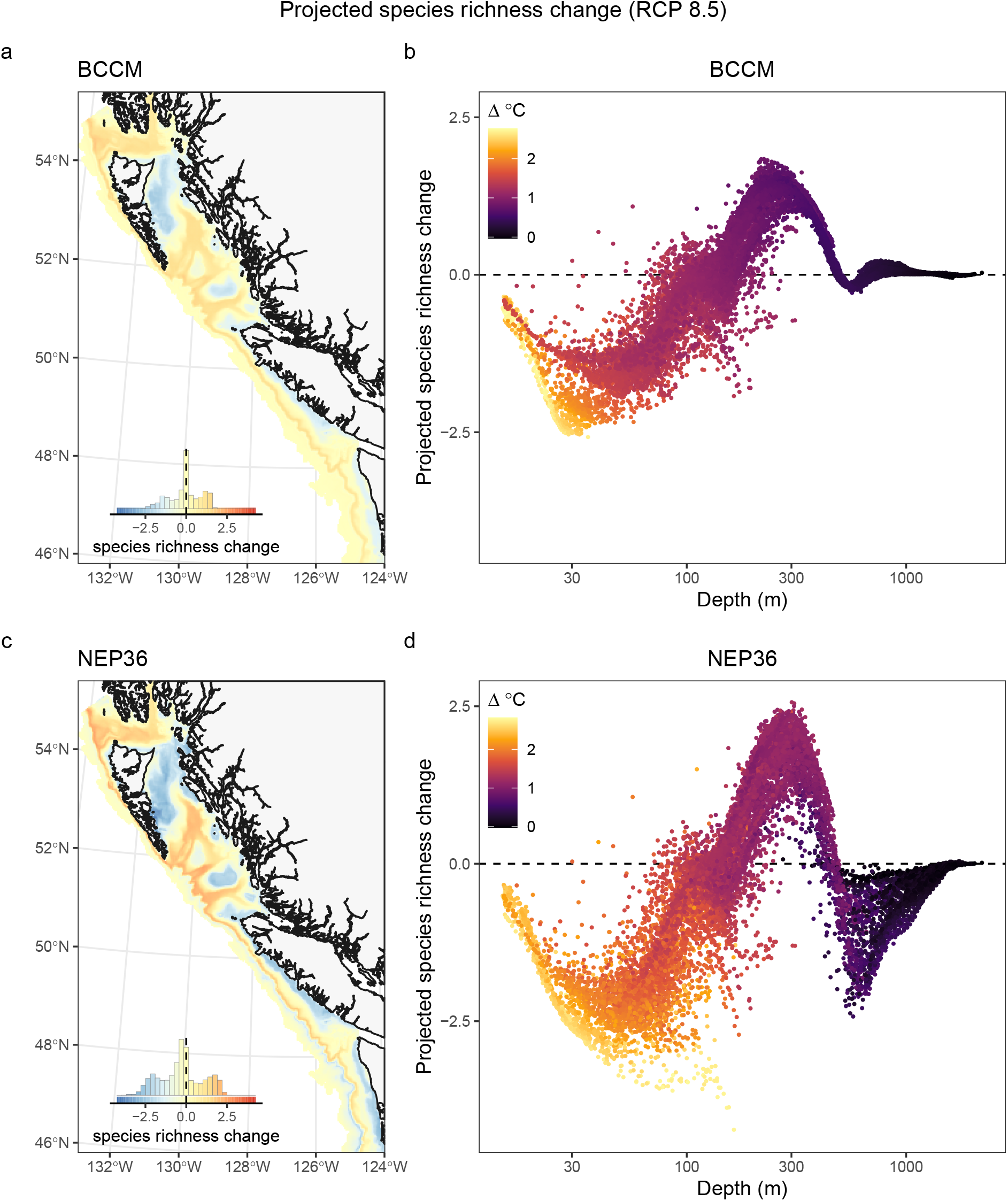
Projected species richness change (a, c) mapped across the study region between historical (1986–2005) and future (2046–2065, RCP 8.5) and as a function of seafloor depth (b, d). Panels (a, b) show projections based on the BCCM model. Panels c and d show projections based on the NEP36 model. The inset histograms in panels (a, c) show the distribution of values across all 3 km^2^ grid cells and provide a legend for the colours shown on the maps. Each point in panels (b, d) represents a single 3 km^2^ grid cell in panels (a, c) coloured by the projected temperature change. See Figure 4 for results based on RCP 4.5.

**Figure S12:**
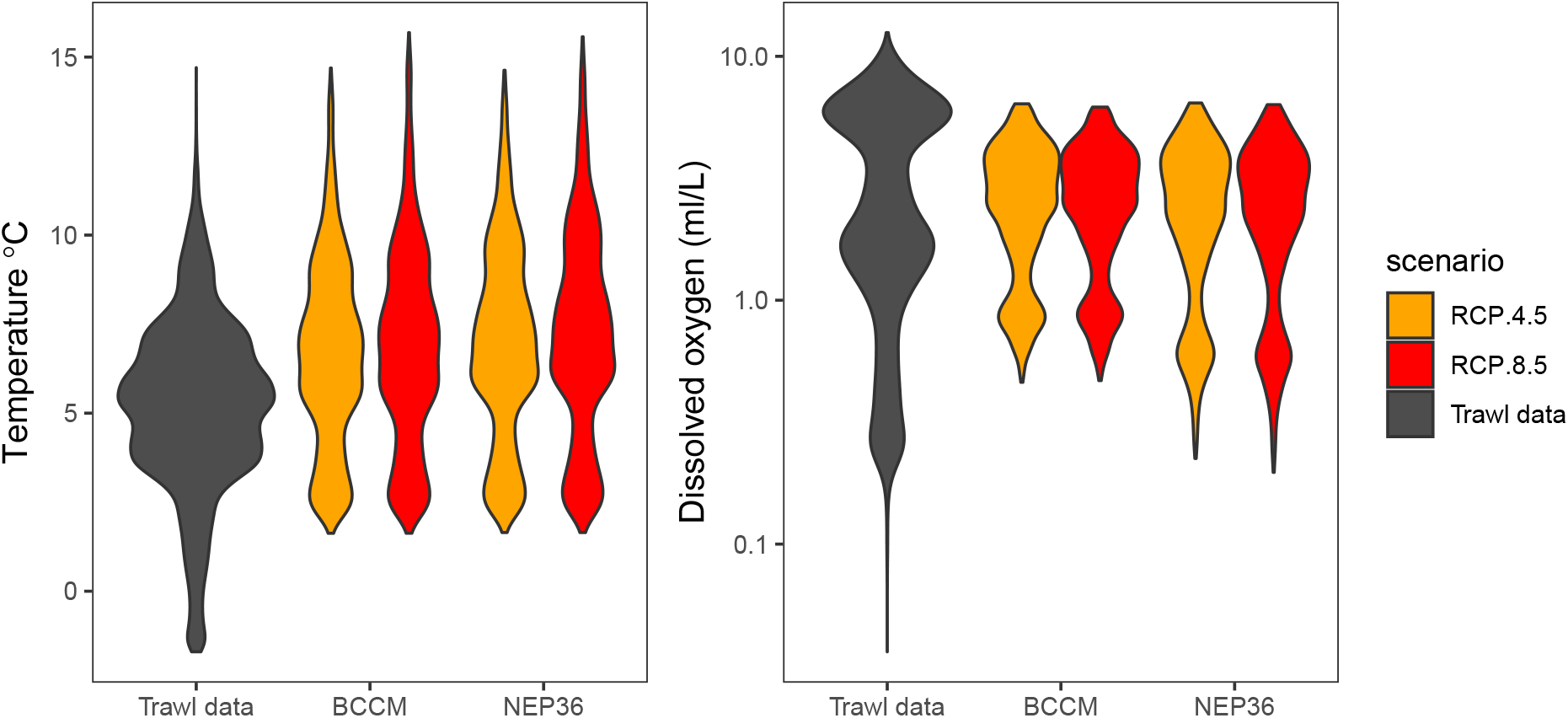
Comparison of the distribution of temperature (a) and dissolved oxygen (b) in the full trawl dataset and the future projected conditions in the two regional ocean models and RCP scenarios.

## References

Aitken, S.N., Yeaman, S., Holliday, J.A., Wang, T. & Curtis-McLane, S. (2008). Adaptation, migration or extirpation: Climate change outcomes for tree populations: Climate change outcomes for tree populations. Evolutionary Applications, 1, 95–111. Retrieved October 19, 2021, from https://onlinelibrary.wiley.com/doi/10.1111/j.1752-4571.2007.00013.x

Alexander, J.M., Diez, J.M. & Levine, J.M. (2015). Novel competitors shape species’ responses to climate change. Nature, 525, 515–518. Retrieved from http://eutils.ncbi.nlm.nih.gov/entrez/eutils/elink.fcgi?dbfrom=pubmed&id=26374998&retmode=ref&cmd=prlinks

Anderson, C., S., Ward, J., E., English, A., P., Barnett & K., L.A. (2022a). sdmTMB: An r package for fast, flexible, and user-friendly generalized linear mixed effects models with spatial and spatiotemporal random fields. bioRxiv, 2022.03.24.485545. Retrieved from https://doi.org/10.1101/2022.03.24.485545

Anderson, S.C., Keppel, E.A. & Edwards, A.M. (2019). A reproducible data synopsis for over 100 species of British Columbia groundfsh. DFO Can. Sci. Advis. Sec. Res. Doc., 041, vii + 321 p.

Anderson, S.C., Ward, E.J., English, P.A. & Barnett, L.A.K. (2022b). sdmTMB: An R package for fast, flexible, and user-friendly generalized linear mixed effects models with spatial and spatiotemporal random fields. bioRxiv, 2022.03.24.485545. Retrieved April 1, 2022, from http://biorxiv.org/lookup/doi/10.1101/2022.03.24.485545

Arora, V.K., Scinocca, J.F., Boer, G.J., Christian, J.R., Denman, K.L., Flato, G.M., Kharin, V.V., Lee, W.G. & Merryfield, W.J. (2011). Carbon emission limits required to satisfy future representative concentration pathways of greenhouse gases: ALLOWABLE FUTURE CARBON EMISSIONS. Geophysical Research Letters, 38, n/a–n/a. Retrieved January 24, 2022, from http://doi.wiley.com/10.1029/2010GL046270

Bartley, T.J., McCann, K.S., Bieg, C., Cazelles, K., Granados, M., Guzzo, M.M., MacDougall, A.S., Tunney, T.D. & McMeans, B.C. (2019). Food web rewiring in a changing world. Nature Ecology and Evolution, 3, 345–354. Retrieved from http://www.nature.com/articles/s41559-018-0772-3

Branch, T.A., DeJoseph, B.M., Ray, L.J. & Wagner, C.A. (2013). Impacts of ocean acidification on marine seafood. Trends in Ecology & Evolution, 28, 178–186. Retrieved January 14, 2022, from https://linkinghub.elsevier.com/retrieve/pii/S0169534712002625

Brodie, S., Smith, J.A., Muhling, B.A., Barnett, L.A.K., Carroll, G., Fiedler, P., Bograd, S.J., Hazen, E.L., Jacox, M.G., Andrews, K.S., Barnes, C.L., Crozier, L.G., Fiechter, J., Fredston, A., Haltuch, M.A., Harvey, C.J., Holmes, E., Karp, M.A., Liu, O.R., Malick, M.J., Pozo Buil, M., Richerson, K., Rooper, C.N., Samhouri, J., Seary, R., Selden, R.L., Thompson, A.R., Tommasi, D., Ward, E.J. & Kaplan, I.C. (2022). Recommendations for quantifying and reducing uncertainty in climate projections of species distributions. Global Change Biology, gcb.16371. Retrieved September 1, 2022, from https://onlinelibrary.wiley.com/doi/10.1111/gcb.16371

Brown, A. & Thatje, S. (2014). Explaining bathymetric diversity patterns in marine benthic invertebrates and demersal fishes: Physiological contributions to adaptation of life at depth. Biological Reviews, 89, 406–426. Retrieved August 27, 2021, from https://onlinelibrary.wiley.com/doi/10.1111/brv.12061

Brown, A. & Thatje, S. (2015). The effects of changing climate on faunal depth distributions determine winners and losers. Global Change Biology, 21, 173–180. Retrieved April 7, 2021, from http://doi.wiley.com/10.1111/gcb.12680

Cavole, L., Demko, A., Diner, R., Giddings, A., Koester, I., Pagniello, C., Paulsen, M.-L., Ramírez-Valdez, A., Schwenck, S., Yen, N., Zill, M. & Franks, P. (2016). Biological impacts of the 2013–2015 warm-water anomaly in the northeast pacific: Winners, losers, and the future. Oceanography (Washington D.C.), 29.

Chaikin, S., Dubiner, S. & Belmaker, J. (2022). Cold-water species deepen to escape warm water temperatures. Global Ecology and Biogeography, 31, 75–88. Retrieved from https://onlinelibrary.wiley.com/doi/abs/10.1111/geb.13414

Cheung, W.W.L., Lam, V.W.Y., Sarmiento, J.L., Kearney, K., Watson, R. & Pauly, D. (2009). Projecting global marine biodiversity impacts under climate change scenarios. Fish and Fisheries, 10, 235–251. Retrieved January 28, 2021, from http://doi.wiley.com/10.1111/j.1467-2979.2008.00315.x

Clarke, T.M., Frölicher, T., Reygondeau, G., Villalobos-Rojas, F., Wabnitz, C.C.C., Wehrtmann, I.S. & Cheung, W.W.L. (2022). Temperature and oxygen supply shape the demersal community in a tropical Oxygen Minimum Zone. Environmental Biology of Fishes. Retrieved October 7, 2022, from https://link.springer.com/10.1007/s10641-022-01256-2

Clarke, T.M., Wabnitz, C.C.C., Striegel, S., Frölicher, T.L., Reygondeau, G. & Cheung, W.W.L. (2021). Aerobic growth index (AGI): An index to understand the impacts of ocean warming and deoxygenation on global marine fisheries resources. Progress in Oceanography, 195, 102588. Retrieved October 7, 2022, from https://linkinghub.elsevier.com/retrieve/pii/S0079661121000756

Clayton, D.G., Bernardinelli, L. & Montomoli, C. (1993). Spatial Correlation in Ecological Analysis. International Journal of Epidemiology, 22, 1193–1202. Retrieved September 20, 2022, from https://academic.oup.com/ije/article-lookup/doi/10.1093/ije/22.6.1193

Cressie, N.A.C. & Wikle, C.K. (2011). Statistics for Spatio-Temporal Data. Wiley, Hoboken, N.J.

Deutsch, C., Ferrel, A., Seibel, B., Portner, H.-O. & Huey, R.B. (2015). Climate change tightens a metabolic constraint on marine habitats. Science, 348, 1132–1135. Retrieved August 4, 2021, from https://www.sciencemag.org/lookup/doi/10.1126/science.aaa1605

Diaz, R.J. & Rosenberg, R. (2008). Spreading Dead Zones and Consequences for Marine Ecosystems. Science, 321, 926–929. Retrieved March 9, 2022, from https://www.science.org/doi/10.1126/science.1156401

Duplisea, D.E., Roux, M.-J., Hunter, K.L. & Rice, J. (2021). Fish harvesting advice under climate change: A risk-equivalent empirical approach (A. Belgrano, Ed.). PLOS ONE, 16, e0239503. Retrieved January 17, 2022, from https://dx.plos.org/10.1371/journal.pone.0239503

Elith, J. & Leathwick, J.R. (2009). Species Distribution Models: Ecological Explanation and Prediction Across Space and Time. Annual Review of Ecology, Evolution, and Systematics, 40, 677–697. Retrieved September 3, 2021, from http://www.annualreviews.org/doi/10.1146/annurev.ecolsys.110308.120159

English, P.A., Ward, E.J., Rooper, C.N., Forrest, R.E., Rogers, L.A., Hunter, K.L., Edwards, A.M., Connors, B.M. & Anderson, S.C. (2022). Contrasting climate velocity impacts in warm and cool locations show that effects of marine warming are worse in already warmer temperate waters. Fish and Fisheries, 17.

Environment and Climate Change Canada. (2021). Canadian Environmental Sustainability Indicators: Canada’s conserved areas. Retrieved March 1, 2022, from https://www.canada.ca/en/environment-climate-change/services/environmental-indicators/conserved-areas.html

Essington, T.E., Anderson, S.C., Barnett, L.A.K., Berger, H.M., Siedlecki, S.A. & Ward, E.J. (2022). Advancing statistical models to reveal the effect of dissolved oxygen on the spatial distribution of marine taxa using thresholds and a physiologically based index. Ecography, 2022. Retrieved August 30, 2022, from https://onlinelibrary.wiley.com/doi/10.1111/ecog.06249

Frazão Santos, C., Agardy, T., Andrade, F., Crowder, L.B., Ehler, C.N. & Orbach, M.K. (2018). Major challenges in developing marine spatial planning. Marine Policy, S0308597X18306213. Retrieved March 1, 2021, from https://linkinghub.elsevier.com/retrieve/pii/S0308597X18306213

Fredston, A., Pinsky, M., Selden, R.L., Szuwalski, C., Thorson, J.T., Gaines, S.D. & Halpern, B.S. (2021). Range edges of North American marine species are tracking temperature over decades. Global Change Biology, 27, 3145–3156. Retrieved August 4, 2021, from https://onlinelibrary.wiley.com/doi/10.1111/gcb.15614

Friesen, S.K., Rubidge, E., Martone, R., Hunter, K.L., Peña, M.A. & Ban, N.C. (2021). Effects of changing ocean temperatures on ecological connectivity among marine protected areas in northern British Columbia. Ocean & Coastal Management, 211, 105776. Retrieved March 3, 2022, from https://linkinghub.elsevier.com/retrieve/pii/S0964569121002593

Frölicher, T.L., Rodgers, K.B., Stock, C.A. & Cheung, W.W.L. (2016). Sources of uncertainties in 21st century projections of potential ocean ecosystem stressors: UNCERTAINTIES IN STRESSOR PROJECTIONS. Global Biogeochemical Cycles, 30, 1224–1243. Retrieved January 24, 2022, from http://doi.wiley.com/10.1002/2015GB005338

Fry, F.E.J. (1971). The Effect of Environmental Factors on the Physiology of Fish. Fish Physiology, pp. 1–98. Elsevier. Retrieved September 27, 2022, from https://linkinghub.elsevier.com/retrieve/pii/S1546509808601466

García Molinos, J., Halpern, B.S., Schoeman, D.S., Brown, C.J., Kiessling, W., Moore, P.J., Pandolfi, J.M., Poloczanska, E.S., Richardson, A.J. & Burrows, M.T. (2016). Climate velocity and the future global redistribution of marine biodiversity. Nature Climate Change, 6, 83–88. Retrieved August 4, 2021, from http://www.nature.com/articles/nclimate2769

Gilman, S.E., Urban, M.C., Tewksbury, J., Gilchrist, G.W. & Holt, R.D. (2010). A framework for community interactions under climate change. Trends in Ecology & Evolution, 25, 325–331. Retrieved from http://linkinghub.elsevier.com/retrieve/pii/S0169534710000613

Giorgi, F., Jones, C. & Asrar, G.R. (2009). Addressing climate information needs at the regional level: The CORDEX framework. World Meteorological Organization (WMO) Bulletin, 53, 175.

Gissi, E., Fraschetti, S. & Micheli, F. (2019). Incorporating change in marine spatial planning: A review. Environmental Science & Policy, 92, 191–200. Retrieved September 1, 2021, from https://linkinghub.elsevier.com/retrieve/pii/S1462901118308517

Guisan, A. & Thuiller, W. (2005). Predicting species distribution: Offering more than simple habitat models. Ecology Letters, 8, 993–1009. Retrieved from http://doi.wiley.com/10.1111/j.1461-0248.2005.00792.x

Guzman, L.M., Germain, R.M., Forbes, C., Straus, S., O’Connor, M.I., Gravel, D., Srivastava, D.S. & Thompson, P.L. (2019). Towards a multi-trophic extension of metacommunity ecology (U. Brose, Ed.). Ecology Letters, 22, 19–33. Retrieved from https://onlinelibrary.wiley.com/doi/abs/10.1111/ele.13162

Haidvogel, D.B., Arango, H., Budgell, W.P., Cornuelle, B.D., Curchitser, E., Di Lorenzo, E., Fennel, K., Geyer, W.R., Hermann, A.J., Lanerolle, L., Levin, J., McWilliams, J.C., Miller, A.J., Moore, A.M., Powell, T.M., Shchepetkin, A.F., Sherwood, C.R., Signell, R.P., Warner, J.C. & Wilkin, J. (2008). Ocean forecasting in terrain-following coordinates: Formulation and skill assessment of the Regional Ocean Modeling System. Journal of Computational Physics, 227, 3595–3624. Retrieved May 20, 2021, from https://linkinghub.elsevier.com/retrieve/pii/S0021999107002549

Haigh, R., Ianson, D., Holt, C.A., Neate, H.E. & Edwards, A.M. (2015). Effects of Ocean Acidification on Temperate Coastal Marine Ecosystems and Fisheries in the Northeast Pacific (V. Thiyagarajan (Rajan), Ed.). PLOS ONE, 10, e0117533. Retrieved April 1, 2022, from https://dx.plos.org/10.1371/journal.pone.0117533

Hausfather, Z. & Peters, G.P. (2020). Emissions – the “business as usual” story is misleading. Nature, 577, 618–620. Retrieved September 8, 2021, from http://www.nature.com/articles/d41586-020-00177-3

Hawkins, E. & Sutton, R. (2009). The Potential to Narrow Uncertainty in Regional Climate Predictions. Bulletin of the American Meteorological Society, 90, 1095–1108. Retrieved January 24, 2022, from https://journals.ametsoc.org/doi/10.1175/2009BAMS2607.1

Hermann, A.J., Gibson, G.A., Cheng, W., Ortiz, I., Aydin, K., Wang, M., Hollowed, A.B. & Holsman, K.K. (2019). Projected biophysical conditions of the Bering Sea to 2100 under multiple emission scenarios (S. Sathyendranath, Ed.). ICES Journal of Marine Science, fsz043. Retrieved March 4, 2022, from https://academic.oup.com/icesjms/advance-article/doi/10.1093/icesjms/fsz043/5477847

Hewitson, B.C. & Crane, R.G. (1996). Climate downscaling: Techniques and application. Climate Research, 7, 85–95.

Hijmans, R.J. (2022). Raster: Geographic data analysis and modeling. Retrieved from https://CRAN.R-project.org/package=raster

Holdsworth, A.M., Zhai, L., Lu, Y. & Christian, J.R. (2021). Future Changes in Oceanography and Biogeochemistry Along the Canadian Pacific Continental Margin. Frontiers in Marine Science, 8, 602991. Retrieved August 27, 2021, from https://www.frontiersin.org/articles/10.3389/fmars.2021.602991/full

Kearney, K., Hermann, A., Cheng, W., Ortiz, I. & Aydin, K. (2020). A coupled pelagic#x2013;benthic#x2013;sympagic biogeochemical model for the Bering Sea: Documentation and validation of the BESTNPZ model (V2019.08.23) within a high-resolution regional ocean model. Geoscientific Model Development, 13, 597–650. Retrieved from https://gmd.copernicus.org/articles/13/597/2020/

Keller, A.A., Wallace, J.R. & Methot, R.D. (2017). The Northwest Fisheries Science Center’s West Coast Groundfish Bottom Trawl Survey : History, design, and description. U.S. Department of Commerce, NOAA Technical Memorandum, **NMFS-NWFSC-136**. Retrieved September 2, 2021, from https://repository.library.noaa.gov/view/noaa/14179

Kelley, D., Richards, C. & SCOR/IAPSO, W. (2021). Gsw: Gibbs sea water functions. Retrieved from https://CRAN.R-project.org/package=gsw

Madec, G. (2016). NEMO ocean engine. Notes du Pôle de modélisation de l’Institut Pierre-Simon Laplace (IPSL), 27, 412.

Marques, I., Kneib, T. & Klein, N. (2022). Mitigating spatial confounding by explicitly correlating Gaussian random fields. Environmetrics, 33. Retrieved September 20, 2022, from https://onlinelibrary.wiley.com/doi/10.1002/env.2727

Morley, J.W., Selden, R.L., Latour, R.J., Frölicher, T.L., Seagraves, R.J. & Pinsky, M.L. (2018). Projecting shifts in thermal habitat for 686 species on the North American continental shelf (B.R. MacKenzie, Ed.). PLOS ONE, 13, e0196127. Retrieved December 29, 2020, from https://dx.plos.org/10.1371/journal.pone.0196127

Moss, R.H., Edmonds, J.A., Hibbard, K.A., Manning, M.R., Rose, S.K., van Vuuren, D.P., Carter, T.R., Emori, S., Kainuma, M., Kram, T., Meehl, G.A., Mitchell, J.F.B., Nakicenovic, N., Riahi, K., Smith, S.J., Stouffer, R.J., Thomson, A.M., Weyant, J.P. & Wilbanks, T.J. (2010). The next generation of scenarios for climate change research and assessment. Nature, 463, 747–756. Retrieved March 3, 2022, from http://www.nature.com/articles/nature08823

Navarro-Racines, C., Tarapues, J., Thornton, P., Jarvis, A. & Ramirez-Villegas, J. (2020). High-resolution and bias-corrected CMIP5 projections for climate change impact assessments. Scientific Data, 7, 7. Retrieved January 5, 2022, from http://www.nature.com/articles/s41597-019-0343-8

Norberg, J., Urban, M.C., Vellend, M., Klausmeier, C.A. & Loeuille, N. (2012). Eco-evolutionary responses of biodiversity to climate change. Nature Climate Change, 2, 747–751. Retrieved from http://dx.doi.org/10.1038/nclimate1588

Nychka, D., Furrer, R., Paige, J. & Sain, S. (2017). Fields: Tools for spatial data. Retrieved from https://github.com/NCAR/Fields

O’Regan, S.M., Archer, S.K., Friesen, S.K. & Hunter, K.L. (2021). A Global Assessment of Climate Change Adaptation in Marine Protected Area Management Plans. Frontiers in Marine Science, 8, 16.

Palacios-Abrantes, J., Frölicher, T.L., Reygondeau, G., Sumaila, U.R., Tagliabue, A., Wabnitz, C.C.C. & Cheung, W.W.L. (2022). Timing and magnitude of climate-driven range shifts in transboundary fish stocks challenge their management. Global Change Biology, 28, 2312–2326. Retrieved March 4, 2022, from https://onlinelibrary.wiley.com/doi/10.1111/gcb.16058

Pearce, J. & Ferrier, S. (2000). Evaluating the predictive performance of habitat models developed using logistic regression. Ecological Modelling, 133, 225–245. Retrieved July 29, 2020, from http://www.sciencedirect.com/science/article/pii/S0304380000003227

Pecl, G.T., Araújo, M.B., Bell, J.D., Blanchard, J., Bonebrake, T.C., Chen, I.-C., Clark, T.D., Colwell, R.K., Danielsen, F., Evengård, B., Falconi, L., Ferrier, S., Frusher, S., Garcia, R.A., Griffis, R.B., Hobday, A.J., Janion-Scheepers, C., Jarzyna, M.A., Jennings, S., Lenoir, J., Linnetved, H.I., Martin, V.Y., McCormack, P.C., McDonald, J., Mitchell, N.J., Mustonen, T., Pandolfi, J.M., Pettorelli, N., Popova, E., Robinson, S.A., Scheffers, B.R., Shaw, J.D., Sorte, C.J.B., Strugnell, J.M., Sunday, J.M., Tuanmu, M.-N., Vergés, A., Villanueva, C., Wernberg, T., Wapstra, E. & Williams, S.E. (2017). Biodiversity redistribution under climate change: Impacts on ecosystems and human well-being. Science, 355, eaai9214. Retrieved February 25, 2022, from https://www.science.org/doi/10.1126/science.aai9214

Peña, M.A., Fine, I. & Callendar, W. (2019). Interannual variability in primary production and shelf-offshore transport of nutrients along the northeast Pacific Ocean margin. Deep Sea Research Part II: Topical Studies in Oceanography, 169–170, 104637. Retrieved July 14, 2020, from http://www.sciencedirect.com/science/article/pii/S0967064519300220

Penn, J.L., Deutsch, C., Payne, J.L. & Sperling, E.A. (2018). Temperature-dependent hypoxia explains biogeography and severity of end-Permian marine mass extinction. Science, 362, eaat1327. Retrieved August 26, 2022, from https://www.science.org/doi/10.1126/science.aat1327

Pinsky, M.L., Fenichel, E., Fogarty, M., Levin, S., McCay, B., St. Martin, K., Selden, R.L. & Young, T. (2021). Fish and fisheries in hot water: What is happening and how do we adapt? Population Ecology, 63, 17–26. Retrieved August 4, 2021, from https://onlinelibrary.wiley.com/doi/10.1002/1438-390X.12050

Pinsky, M.L., Selden, R.L. & Kitchel, Z.J. (2020). Climate-Driven Shifts in Marine Species Ranges: Scaling from Organisms to Communities. Annual Review of Marine Science, 12, 153–179. Retrieved August 4, 2021, from https://www.annualreviews.org/doi/10.1146/annurev-marine-010419-010916

Pinsky, M.L., Worm, B., Fogarty, M.J., Sarmiento, J.L. & Levin, S.A. (2013). Marine Taxa Track Local Climate Velocities. Science, 341, 1239–1242. Retrieved December 29, 2020, from https://www.sciencemag.org/lookup/doi/10.1126/science.1239352

Pirtle, J.L., Shotwell, S.K., Zimmermann, M., Reid, J.A. & Golden, N. (2019). Habitat suitability models for groundfish in the Gulf of Alaska. Deep Sea Research Part II: Topical Studies in Oceanography, 165, 303–321. Retrieved January 14, 2022, from https://linkinghub.elsevier.com/retrieve/pii/S0967064517304332

Poloczanska, E.S., Brown, C.J., Sydeman, W.J., Kiessling, W., Schoeman, D.S., Moore, P.J., Brander, K., Bruno, J.F., Buckley, L.B., Burrows, M.T., Duarte, C.M., Halpern, B.S., Holding, J., Kappel, C.V., O’Connor, M.I., Pandolfi, J.M., Parmesan, C., Schwing, F., Thompson, S.A. & Richardson, A.J. (2013). Global imprint of climate change on marine life. Nature Climate Change, 3, 919–925. Retrieved August 4, 2021, from http://www.nature.com/articles/nclimate1958

Prince, E.D., Luo, J., Phillip Goodyear, C., Hoolihan, J.P., Snodgrass, D., Orbesen, E.S., Serafy, J.E., Ortiz, M. & Schirripa, M.J. (2010). Ocean scale hypoxia-based habitat compression of Atlantic istiophorid billfishes: Hypoxia-based habitat compression of billfishes. Fisheries Oceanography, 19, 448–462. Retrieved August 4, 2021, from https://onlinelibrary.wiley.com/doi/10.1111/j.1365-2419.2010.00556.x

R Core Team. (2021). R: A language and environment for statistical computing. R Foundation for Statistical Computing, Vienna, Austria. Retrieved from https://www.R-project.org/

Ready, J., Kaschner, K., South, A.B., Eastwood, P.D., Rees, T., Rius, J., Agbayani, E., Kullander, S. & Froese, R. (2010). Predicting the distributions of marine organisms at the global scale. Ecological Modelling, 221, 467–478. Retrieved April 20, 2022, from https://linkinghub.elsevier.com/retrieve/pii/S030438000900711X

Roux, M.-J., Duplisea, D.E., Hunter, K.L. & Rice, J. (2022). Consistent Risk Management in a Changing World: Risk Equivalence in Fisheries and Other Human Activities Affecting Marine Resources and Ecosystems. Frontiers in Climate, 3, 781559. Retrieved March 3, 2022, from https://www.frontiersin.org/articles/10.3389/fclim.2021.781559/full

Sadorus, L., Walker, J. & Sullivan, M. (2016). IPHC oceanographic data collection program 2000-2014. International Pacific Halibut Commission, Seattle, Washington.

Scinocca, J.F., Kharin, V.V., Jiao, Y., Qian, M.W., Lazare, M., Solheim, L., Flato, G.M., Biner, S., Desgagne, M. & Dugas, B. (2016). Coordinated Global and Regional Climate Modeling*. Journal of Climate, 29, 17–35. Retrieved March 3, 2022, from http://journals.ametsoc.org/doi/10.1175/JCLI-D-15-0161.1

Sinclair, A., Schnute, J., Haigh, R., Starr, P., Stanley, R., Fargo, J. & Workman, G. (2003). Feasibility of multispecies groundfish bottom trawl surveys on the BC coast. DFO Can. Sci. Advis. Sec. Res. Doc., 049. Retrieved from http://waves-vagues.dfo-mpo.gc.ca/Library/274196.pdf

Stauffer, G. (2004). NOAA Protocols for Groundfish Bottom Trawl Surveys of the Nation’s Fishery Resources. U.S. Dep. Commerce, NOAA Tech. Memo., **NMFS-F/SPO-65**, 205.

Stefánsson, U. & Richards, F.A. (1964). Distributions of dissolved oxygen, density, and nutrients off the Washington and Oregon coasts. Deep Sea Research and Oceanographic Abstracts, 11, 355–380. Retrieved March 4, 2022, from https://linkinghub.elsevier.com/retrieve/pii/0011747164905340

Sunday, J.M., Bates, A.E. & Dulvy, N.K. (2012). Thermal tolerance and the global redistribution of animals. Nature Climate Change, 2, 686–690. Retrieved from http://dx.doi.org/10.1038/nclimate1539

Sunday, J.M., Howard, E., Siedlecki, S., Pilcher, D.J., Deutsch, C., MacCready, P., Newton, J. & Klinger, T. (2022). Biological sensitivities to high-resolution climate change projections in the California current marine ecosystem. Global Change Biology, gcb.16317. Retrieved August 23, 2022, from https://onlinelibrary.wiley.com/doi/10.1111/gcb.16317

Thompson, P.L., Anderson, S.C., Nephin, J., Haagarty, D., Peña, M.A., English, P.A., Gale, K.S.P. & Rubidge, E. (2022a). Disentangling the impacts of environmental change and commercial fishing on demersal fish biodiversity in a northeast Pacific ecosystem. Marine Ecology Progress Series, 689, 137–154.

Thompson, P.L., Anderson, S.C., Nephin, J., Robb, C.K., Proudfoot, B., Park, A.E., Haggarty, D.R. & Rubidge, E. (2022b). Integrating trawl and longline surveys across British Columbia improves groundfish distribution predictions. Canadian Journal of Fisheries and Aquatic Sciences, **In press.**

Thompson, P.L. & Fronhofer, E.A. (2019). The conflict between adaptation and dispersal for maintaining biodiversity in changing environments. Proceedings of the National Academy of Sciences, 116, 21061–21067. Retrieved from http://www.pnas.org/lookup/doi/10.1073/pnas.1911796116

Thompson, P.L. & Gonzalez, A. (2017). Dispersal governs the reorganization of ecological networks under environmental change. Nature Ecology and Evolution, 1, 0162. Retrieved from http://www.nature.com/articles/s41559-017-0162

Tjur, T. (2009). Coefficients of Determination in Logistic Regression Models—A New Proposal: The Coefficient of Discrimination. The American Statistician, 63, 366–372. Retrieved July 29, 2020, from https://amstat.tandfonline.com/doi/abs/10.1198/tast.2009.08210

Urban, M.C., Bocedi, G., Hendry, A., Mihoub, J.B., Peer, G., Singer, A., Bridle, J.R., Crozier, L.G., De Meester, L., Godsoe, W., Gonzalez, A., Hellmann, J.J., Holt, R.D., Huth, A., Johst, K., Krug, C.B., Leadley, P.W., Palmer, S.C.F., Pantel, J.H., Schmitz, A., Zollner, P.A. & Travis, J.M.J. (2016). Improving the forecast for biodiversity under climate change. Science, 353, aad8466–aad8466. Retrieved from http://www.sciencemag.org/cgi/doi/10.1126/science.aad8466

Urban, M.C., Tewksbury, J.J. & Sheldon, K.S. (2012). On a collision course: Competition and dispersal differences create no-analogue communities and cause extinctions during climate change. Proceedings of the Royal Society B-Biological Sciences, 279, 2072–2080. Retrieved from http://rspb.royalsocietypublishing.org/cgi/doi/10.1098/rspb.2011.2367

Veloz, S.D., Williams, J.W., Blois, J.L., He, F., Otto-Bliesner, B. & Liu, Z. (2012). No-analog climates and shifting realized niches during the late quaternary: Implications for 21st-century predictions by species distribution models. Global Change Biology, 18, 1698–1713. Retrieved January 17, 2022, from https://onlinelibrary.wiley.com/doi/10.1111/j.1365-2486.2011.02635.x

Veneziani, M., Edwards, C.A., Doyle, J.D. & Foley, D. (2009). A central California coastal ocean modeling study: 1. Forward model and the influence of realistic versus climatological forcing. Journal of Geophysical Research, 114, C04015. Retrieved March 4, 2022, from http://doi.wiley.com/10.1029/2008JC004774

Wilson, K.L., Tittensor, D.P., Worm, B. & Lotze, H.K. (2020). Incorporating climate change adaptation into marine protected area planning. Global Change Biology, 26, 3251–3267. Retrieved January 15, 2021, from https://onlinelibrary.wiley.com/doi/abs/10.1111/gcb.15094

