## Supplemental for "Groundfish biodiversity change in northeastern Pacific waters under projected warming and deoxygenation"

Appendix A

Patrick L. Thompson, Jessica Nephin, Sarah Davies, Ashley Park, Devin Lyons, Christopher N. Rooper, M. Angelica Peña, James Christian, Karen Hunter, Emily Rubidge, Amber M. Holdsworth

#### Single species supplemental figures

The following text outlines the elements of the Supplemental Figures. Each figure corresponds to a single species, labeled at the top of the figure.

**Panels a–d** show modelled (contour lines) and observed (coloured points) occurrences based on seafloor depth and temperature (a,c) or temperature and dissolved oxygen (b, d). Panels a and b show observed data from the entire coast, panels c and d show the subset of this data from the focal region where future projections were made. The coloured circles represent gridded values across this parameter space, with the size of each circle showing the number of corresponding trawls, and the colour showing the proportion of trawls where the species was present. The SDM modelled responses are shown as contour lines over this environmental space, with each line corresponding to a given probability of occurrence as indicated by its colour. These SDM modelled responses were fit on the entire coastwide dataset (a–b), but we include these contour lines in panels c and d to show how these SDM modelled fits correspond to the environmental space and observations in the focal region. The model response curves for seafloor depth and temperature are produced by making predictions using the fitted SDM model across a regular grid (30 x 30) of environmental conditions from the minimum to the maximum depth (log) and temperature values in the coastwide dataset, excluding combinations that fell outside of the observed environmental space. Dissolved oxygen for each depth, temperature combination was obtained by binning trawls within this grid and calculating the median value. The model response curves for seafloor temperature and oxygen (b, d) were produced in the same way, but with depth instead of oxygen estimated from the binned data.

**Panels e–h** shows projected changes in occurrence between 1986–2005 and 2046–2065. These are based on applying the fitted SDM model to seafloor depth, temperature, and dissolved oxygen at a 3 km resolution from a hindcast (1986–2005) and future projection (2046–2065) of the BCCM (e, g; Peña *et al.* 2019) and the NEP36 model (f, h; Holdsworth *et al.* 2021). Panels e and f show projections based on the RCP 4.5 emissions scenario, panels g and h show projections based on the RCP 8.5 emissions scenario. The map shows the difference between the hindcast and future projected mean occurrences, with blue shades indicating increased occurrence and red and orange shades indicating decreases in occurrence. We assume that areas with an occurrence probability of 0.1 or greater are suitable habitat. Grid cells where both the hindcast and the future projection fall below this threshold are coloured grey. The inset histogram serves as a colour legend and shows the distribution of projected occurrence change values across all 3 km grid cells in the focal region.

### Arrowtooth Flounder

*Atheresthes stomias*

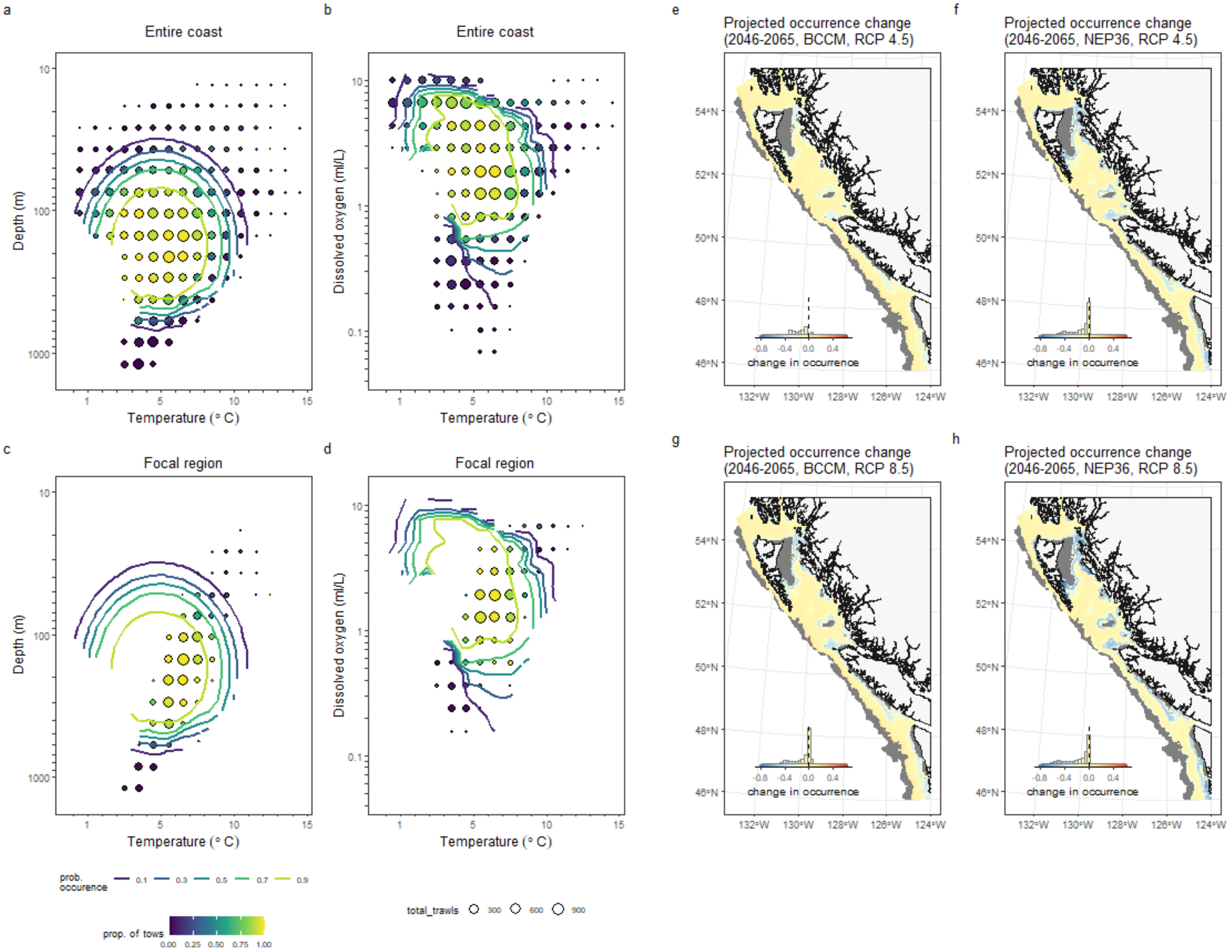


2

#### Big Skate
*Beringraja binoculata*

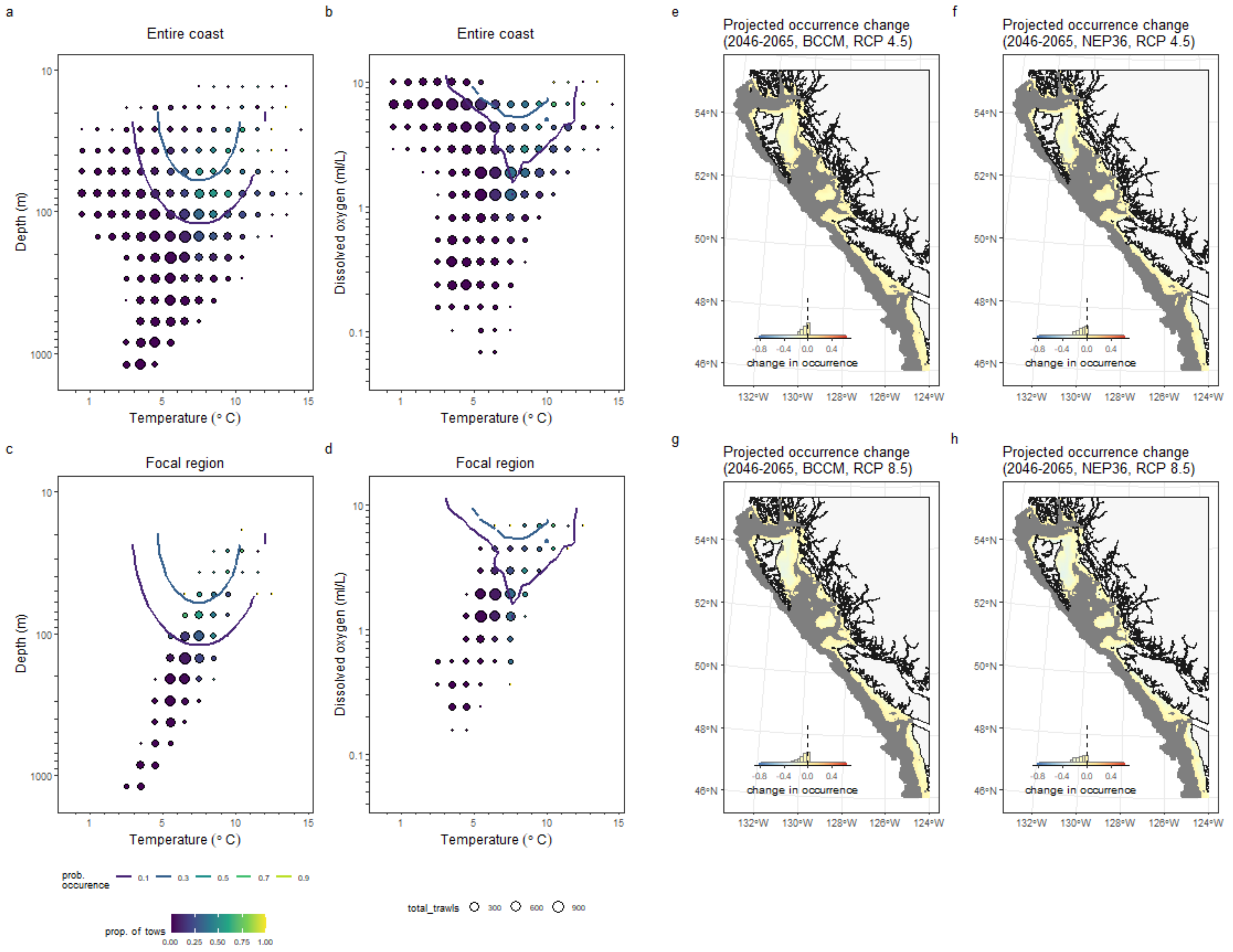


3

### Blacktail Snailfish
*Careproctus melanurus*

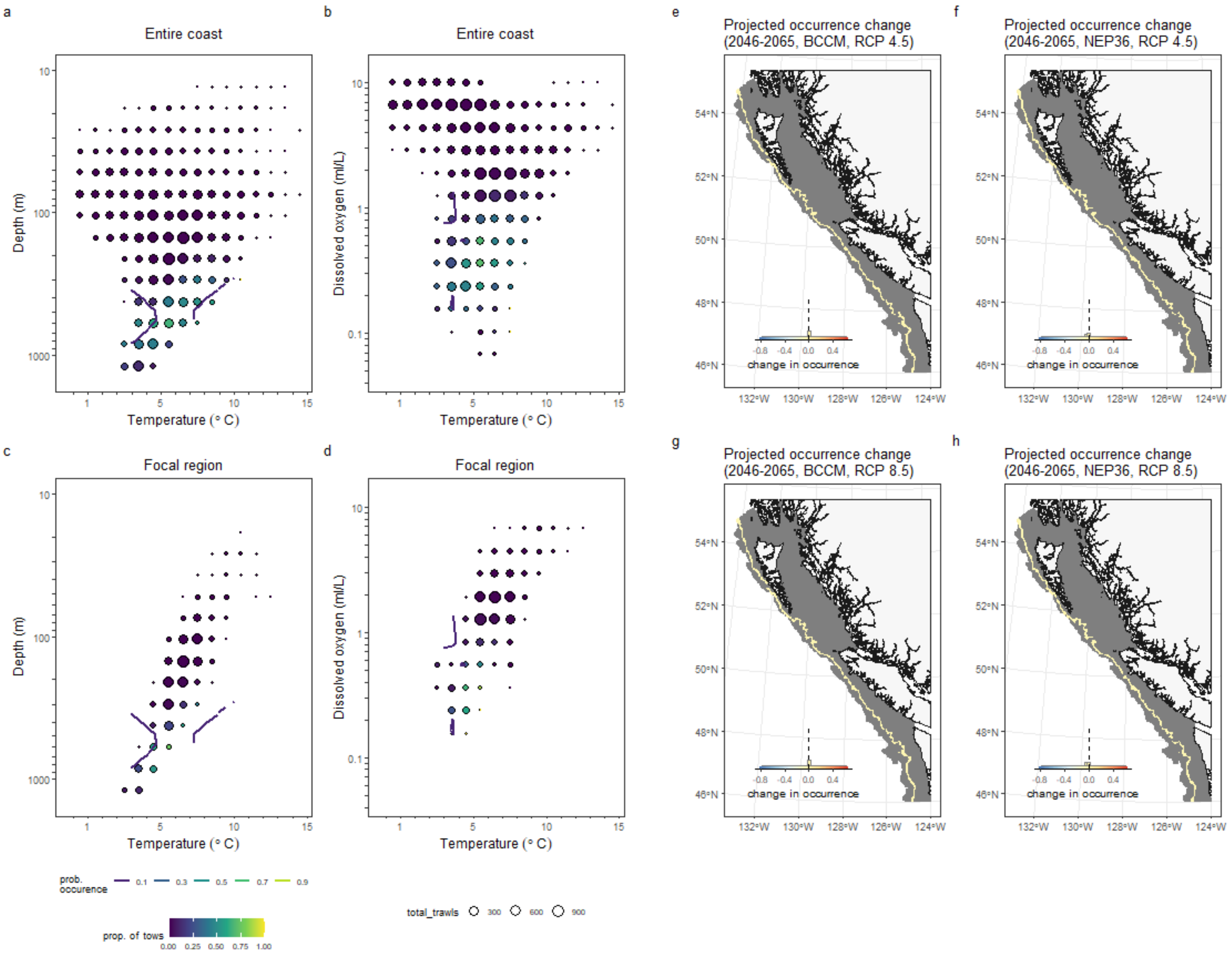

### Brown Cat Shark

*Apristurus brunneus*

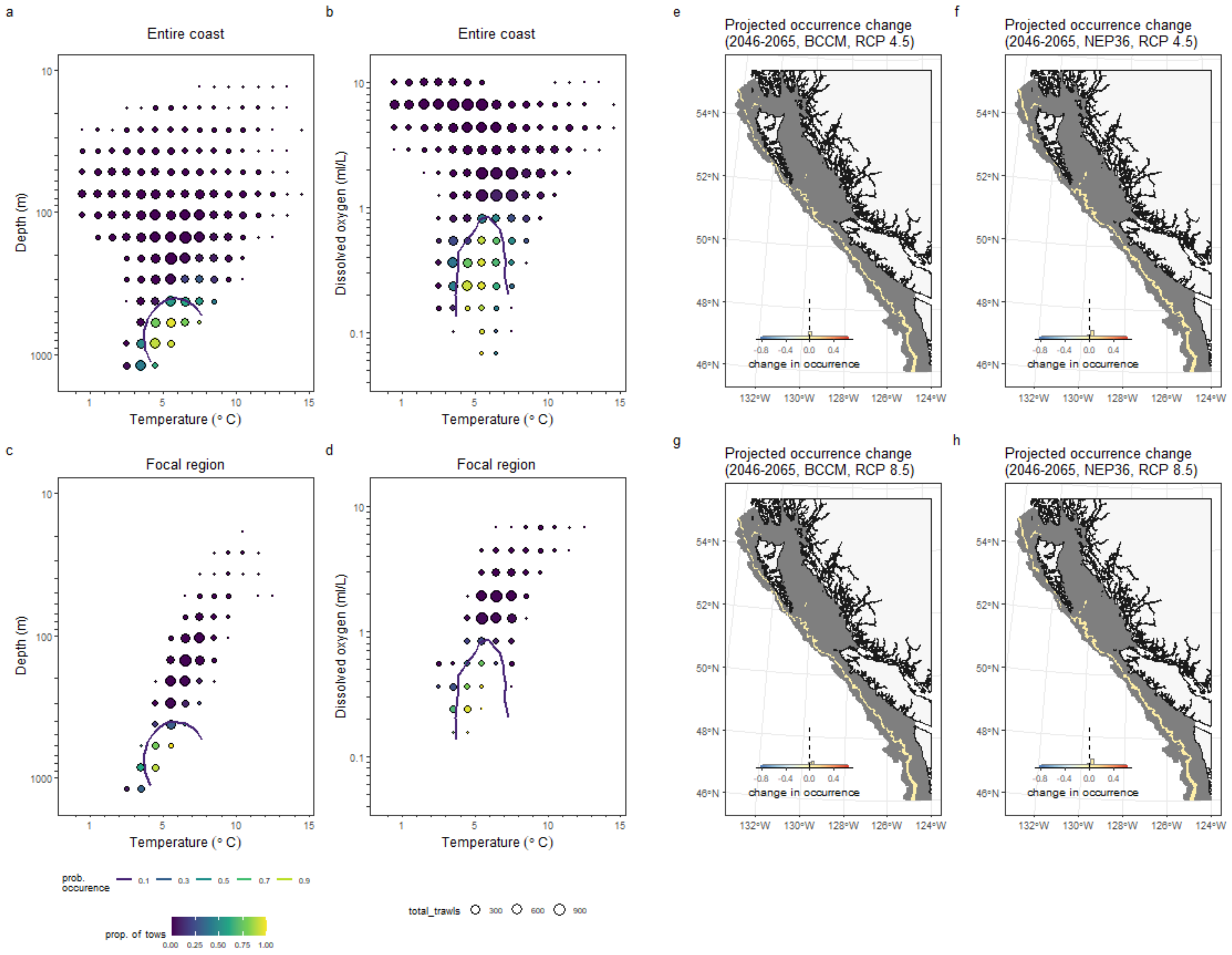

### Curlfin Sole

*Pleuronichthys decurrens*

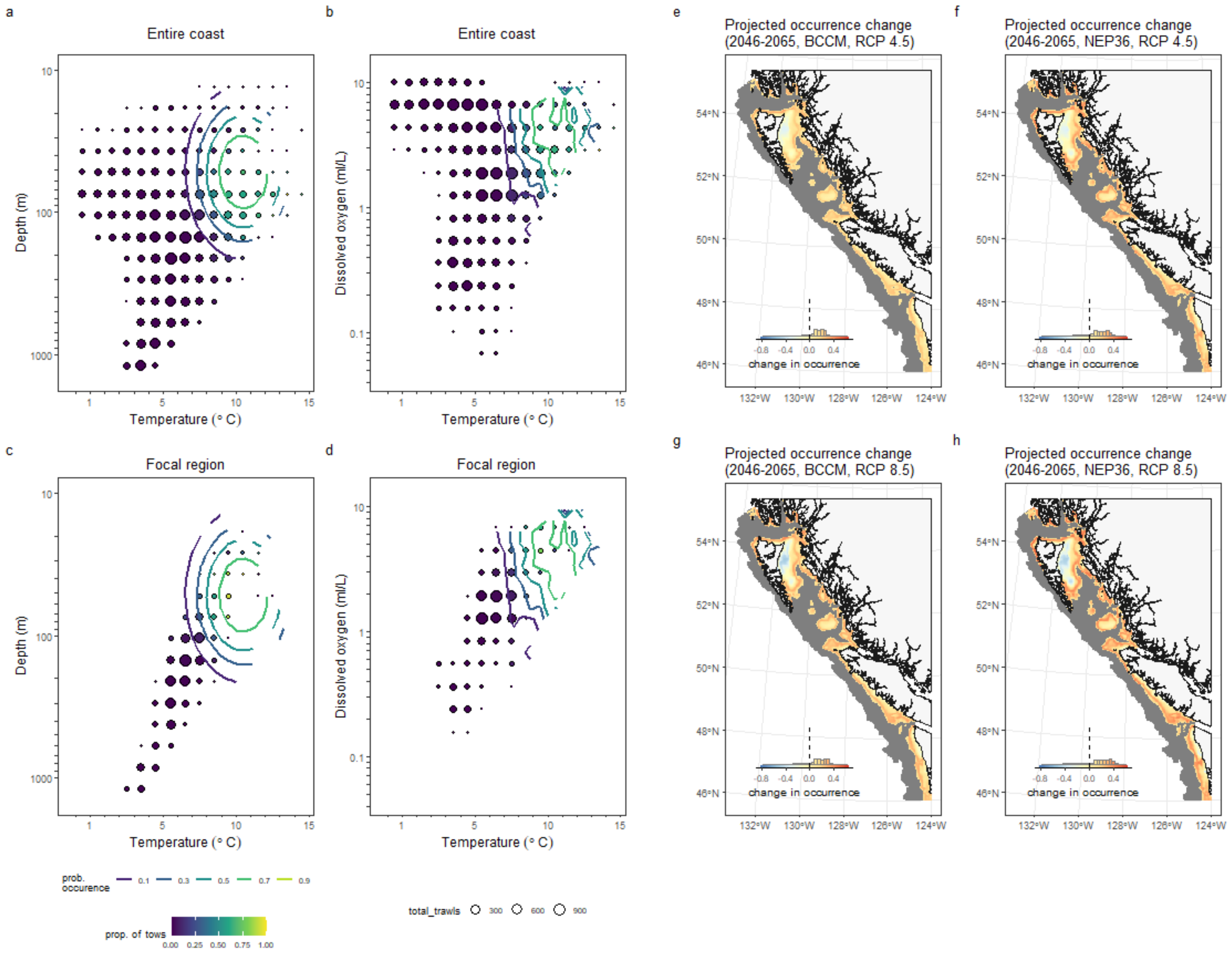

### Deepsea Sole

*Embassichthys bathybius*

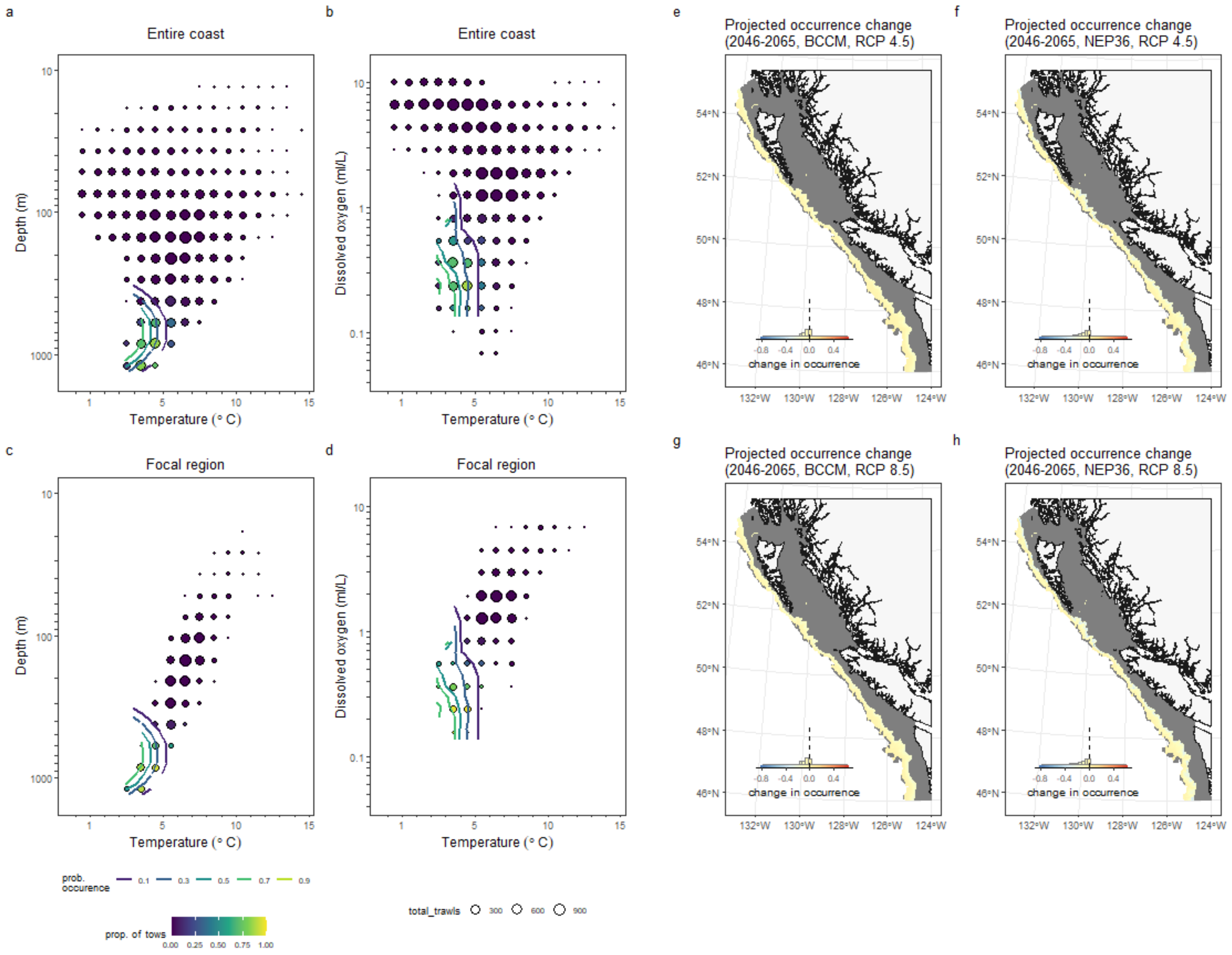

### Dover Sole

*Microstomus pacificus*

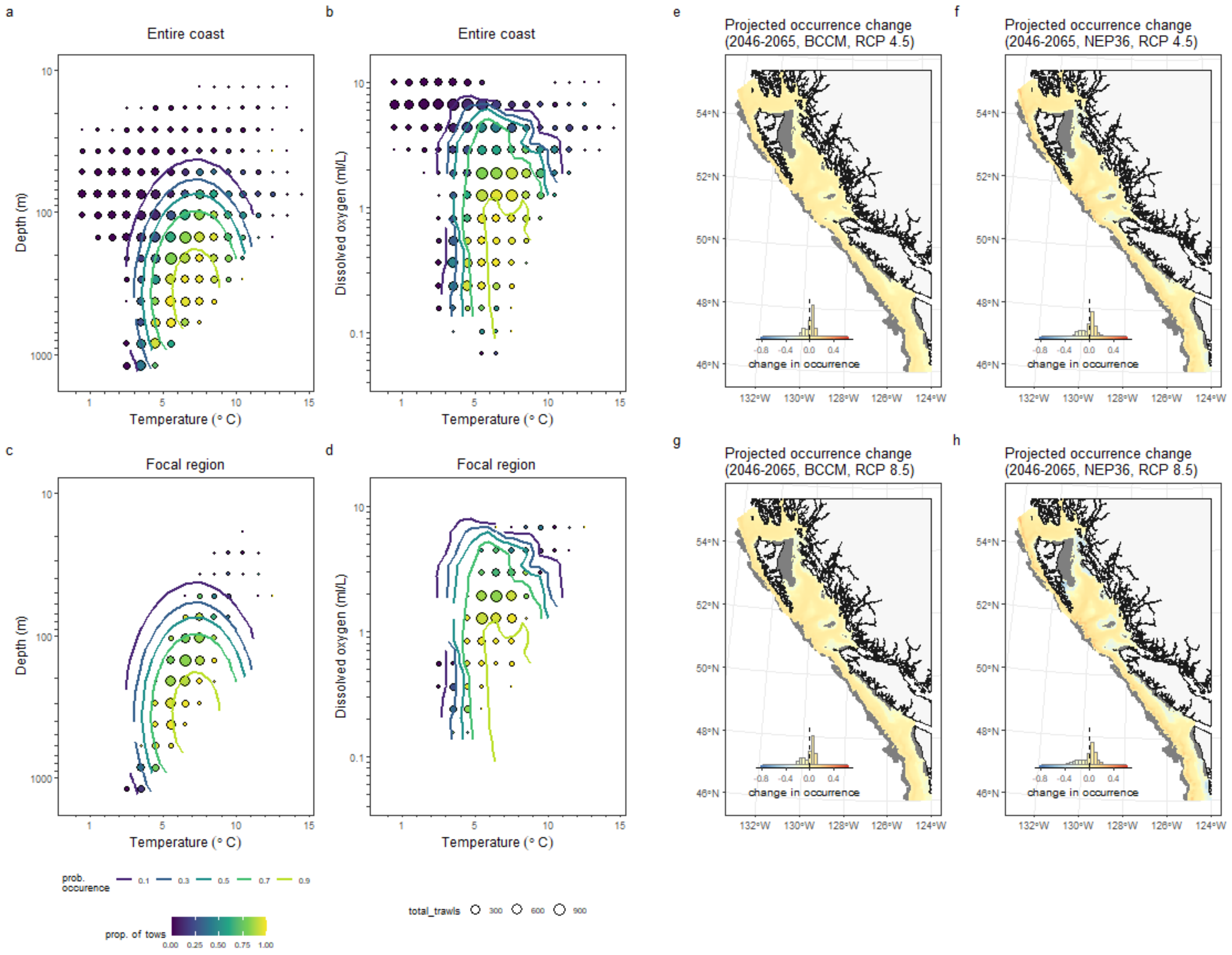

### English Sole
*Parophrys vetulus*

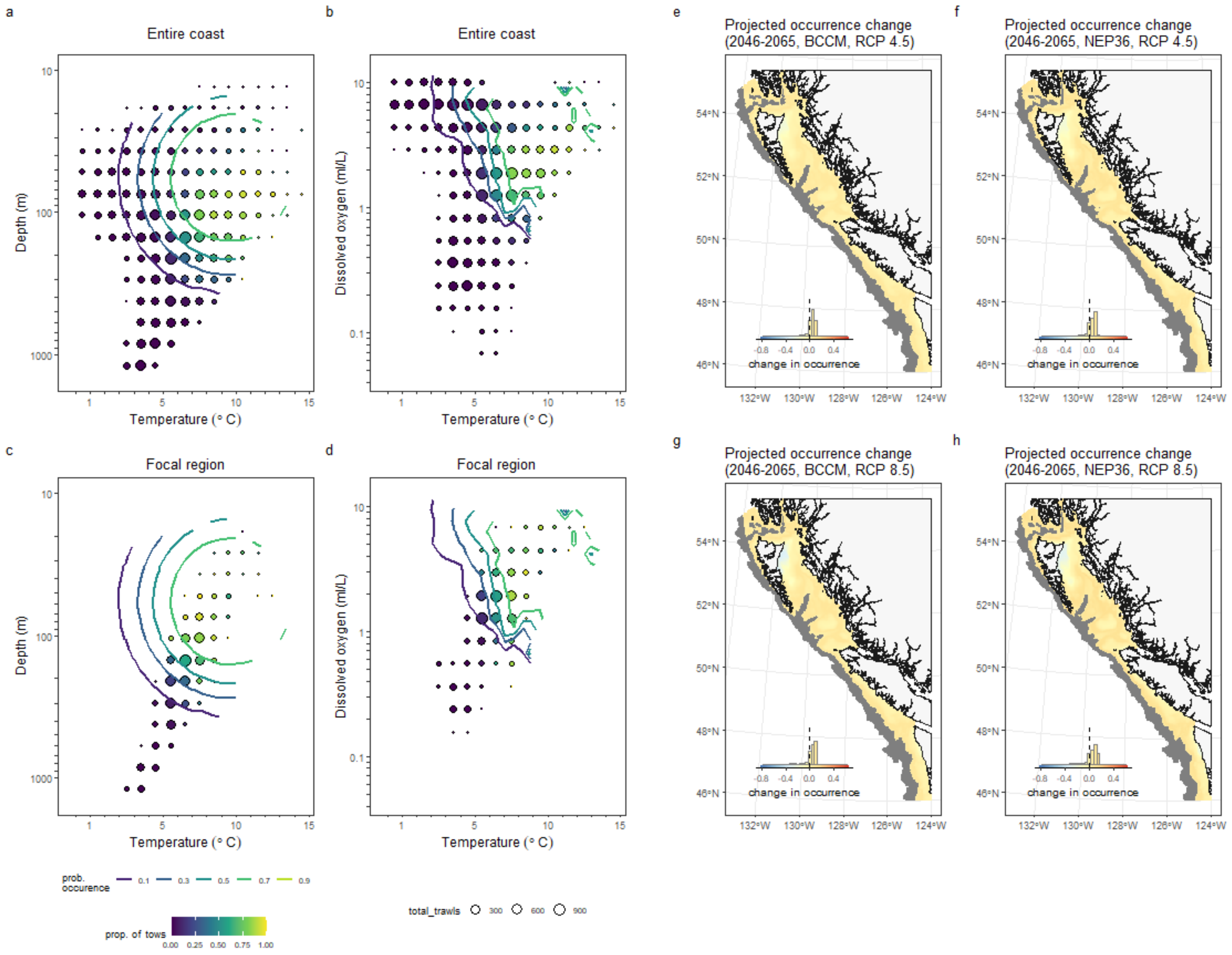

#### Greenstriped Rockfish
*Sebastes elongatus*

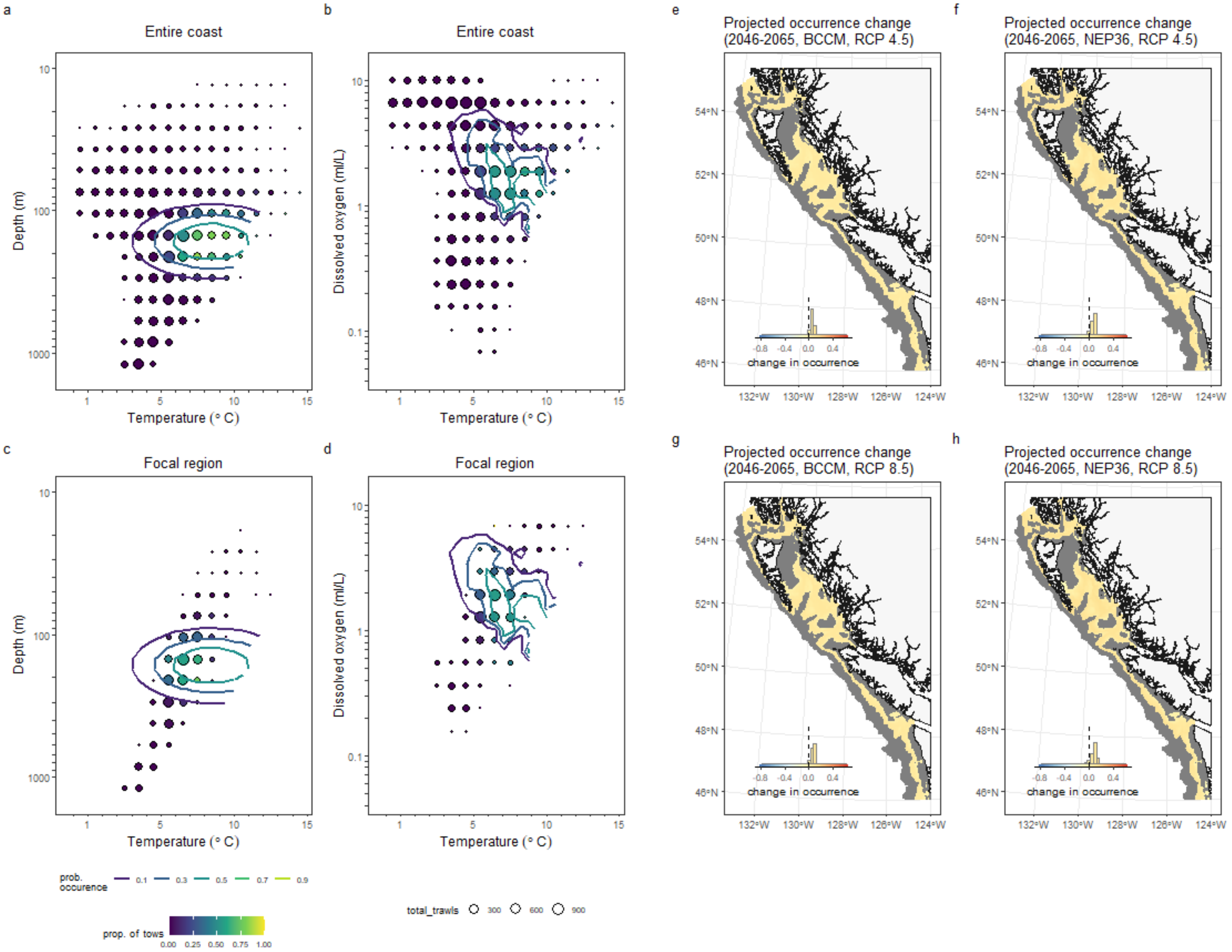

#### Longspine Thornyhead
*Sebastolobus altivelis*

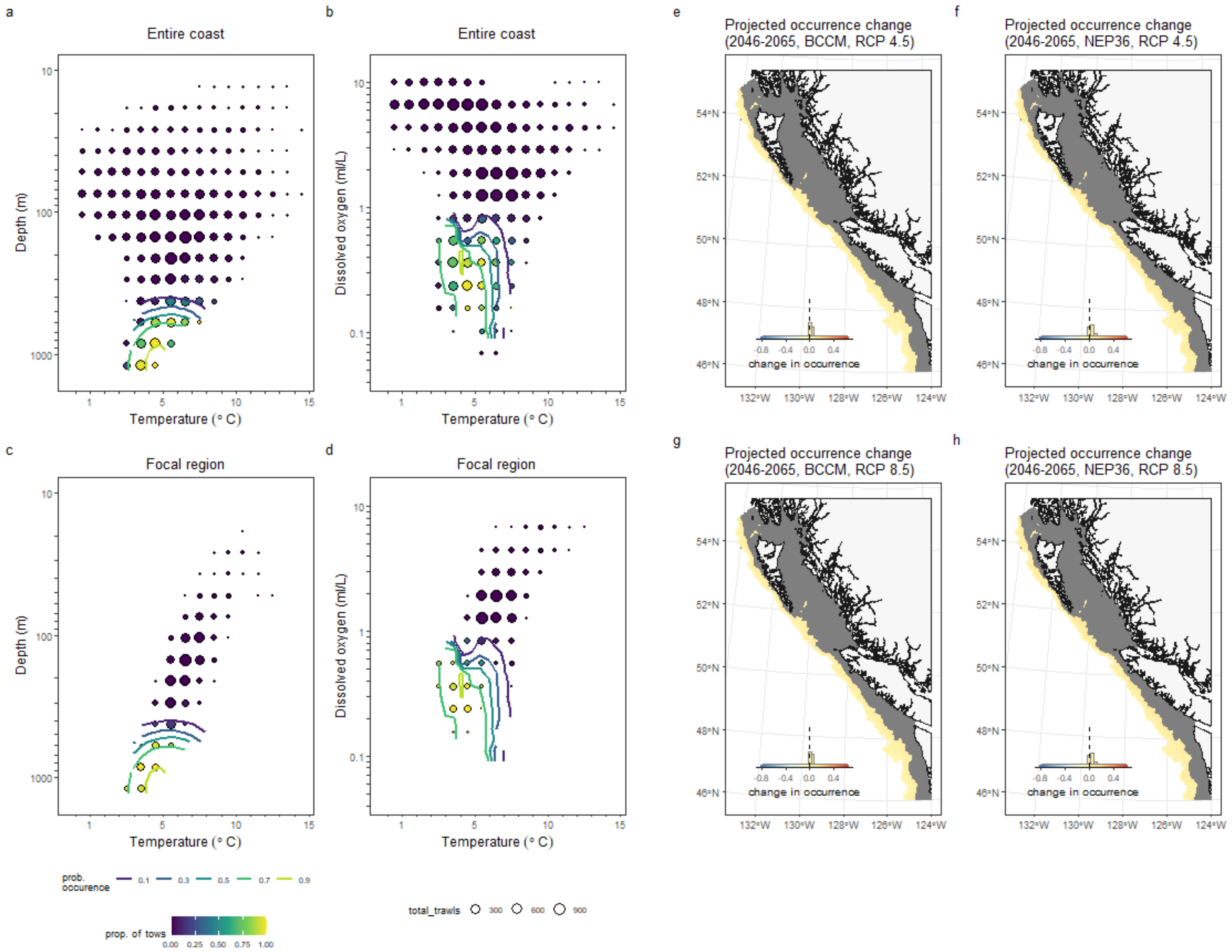

### Pacific Cod
*Gadus macrocephalus*

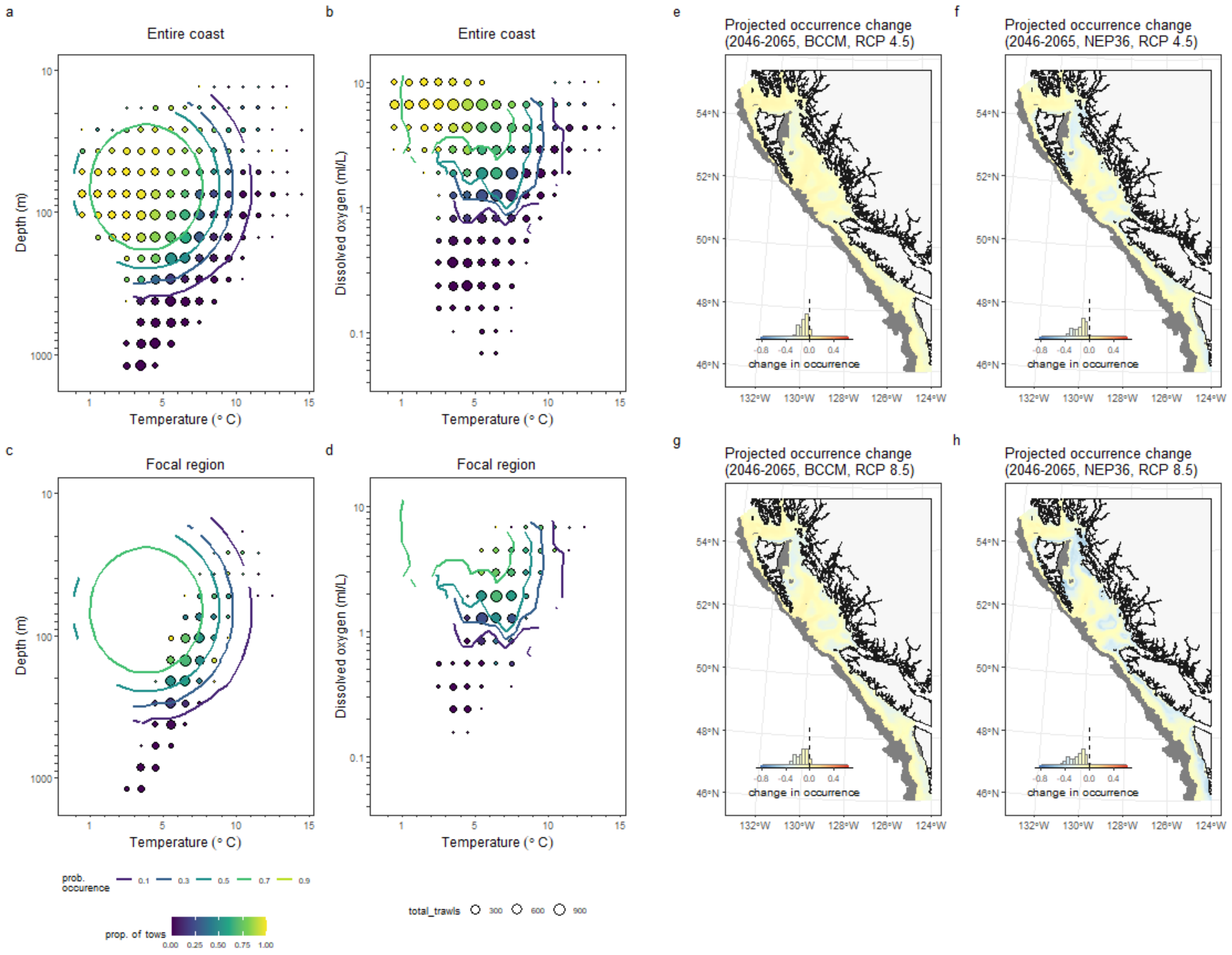

#### Pacific Grenadier

*Coryphaenoides acrolepis*

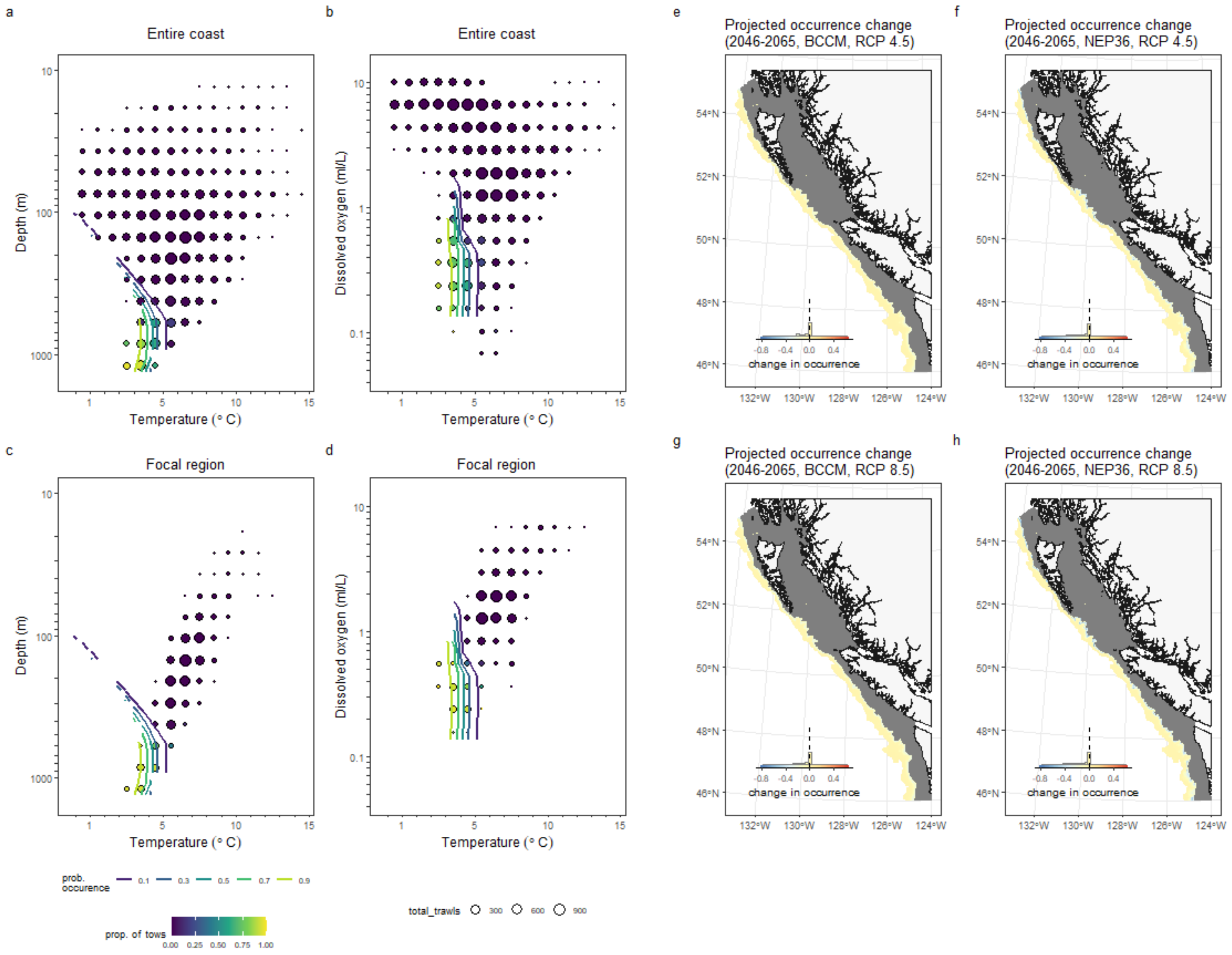

### Pacific Ocean Perch

*Sebastes alutus*

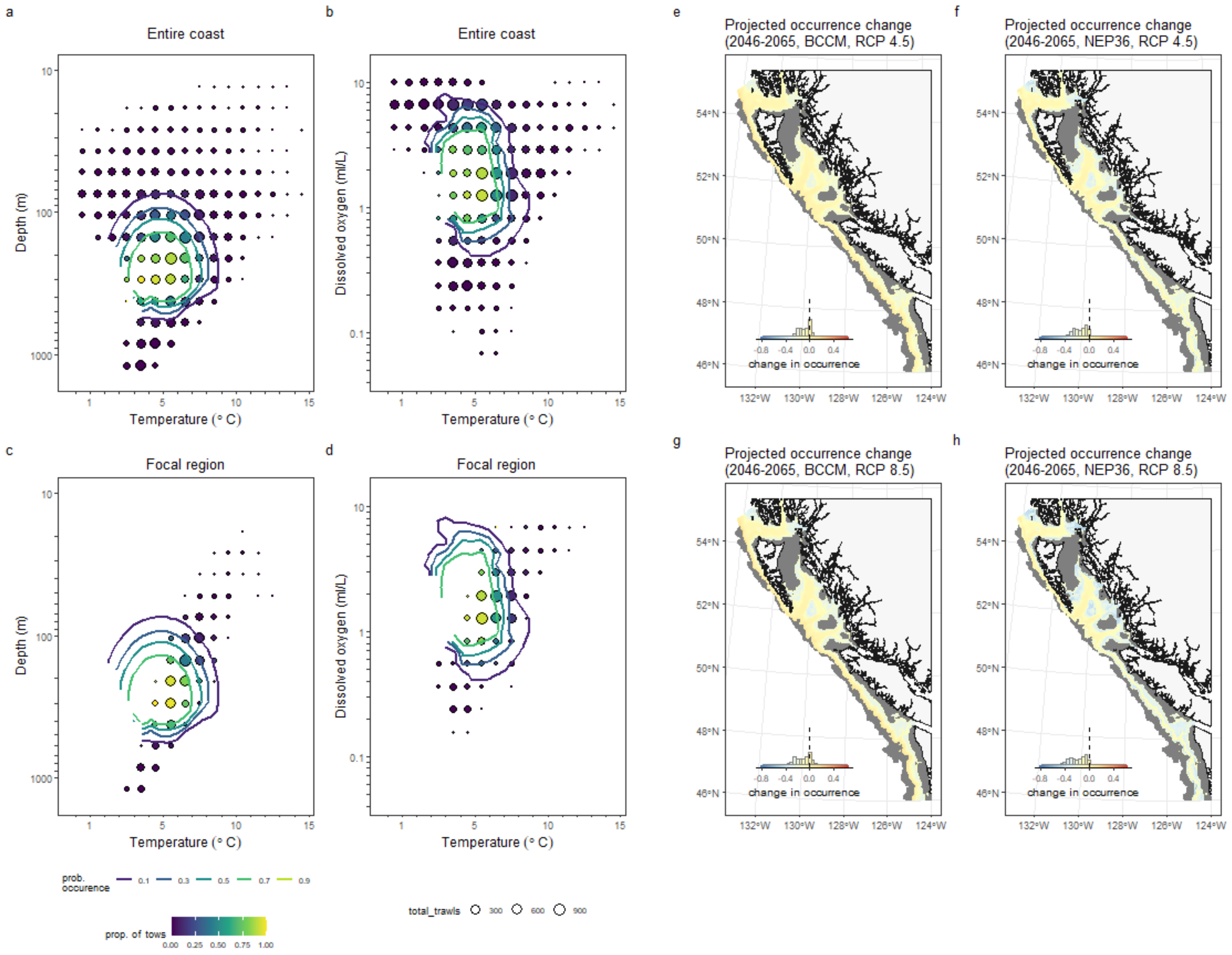

### Pacific Sanddab
*Citharichthys sordidus*

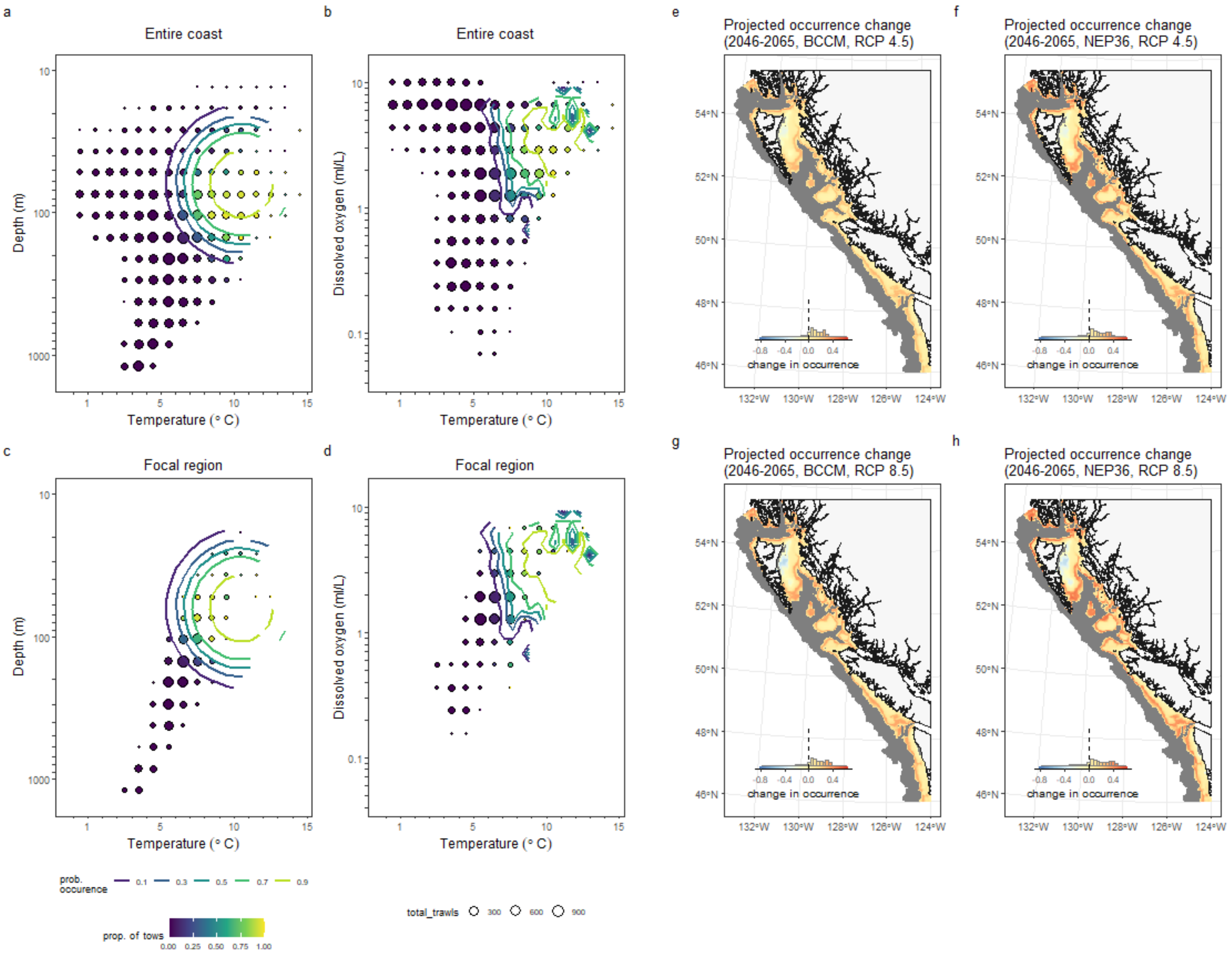

#### Pacific Tomcod

*Microgadus proximus*

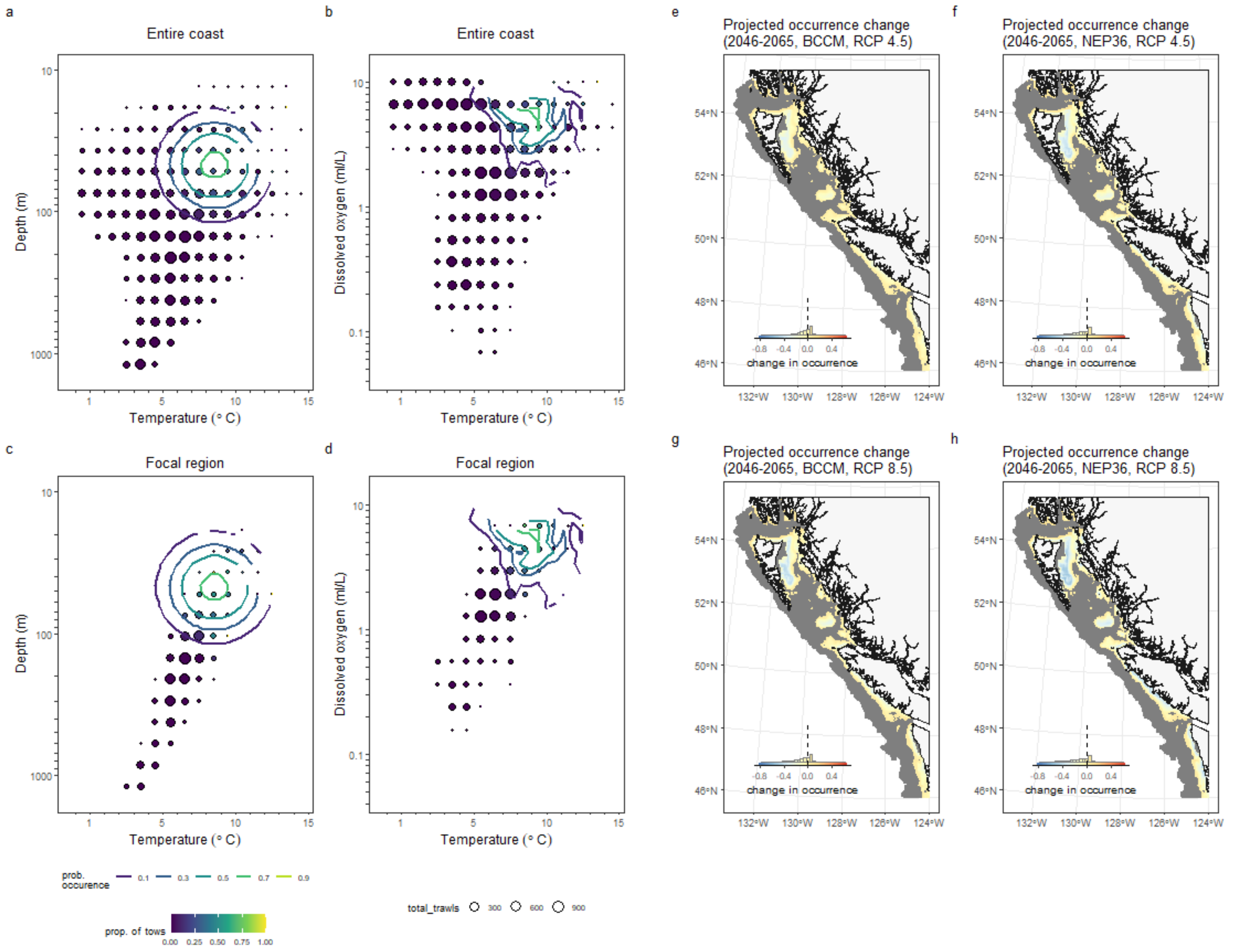

### Pacific Viperfish

*Chauliodus macouni*

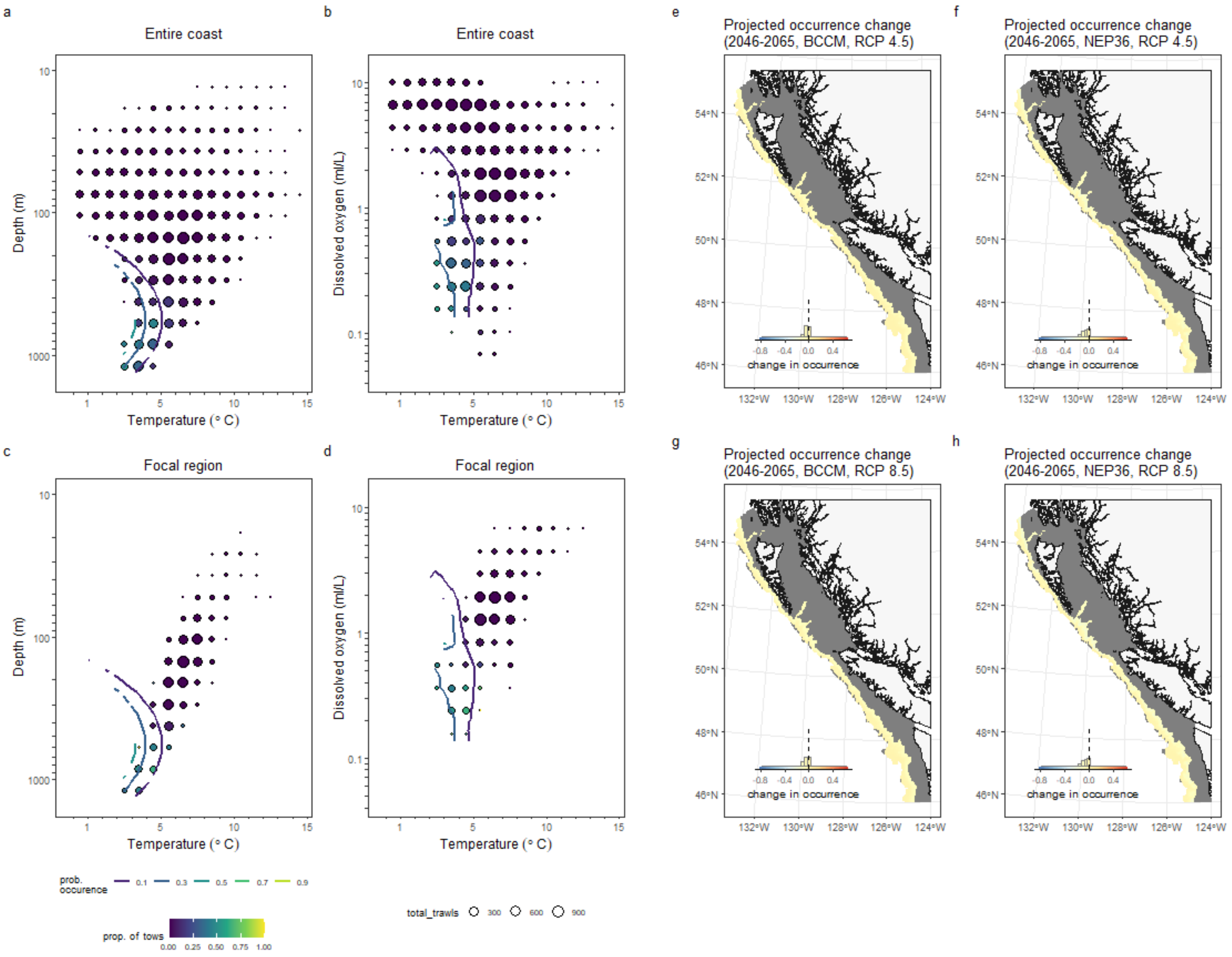

Petrale Sole

*Eopsetta jordani*

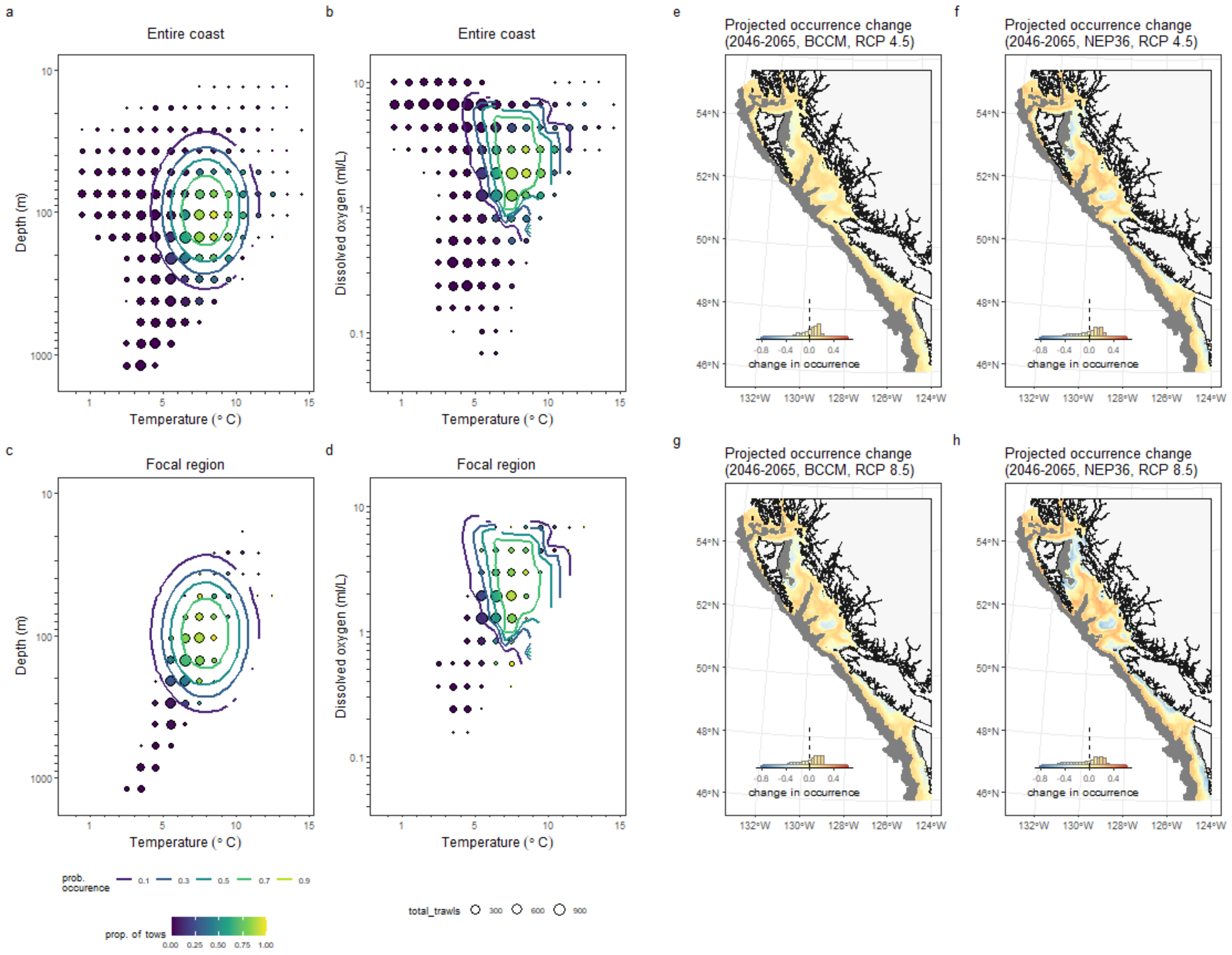

### Popeye
*Coryphaenoides cinereus*

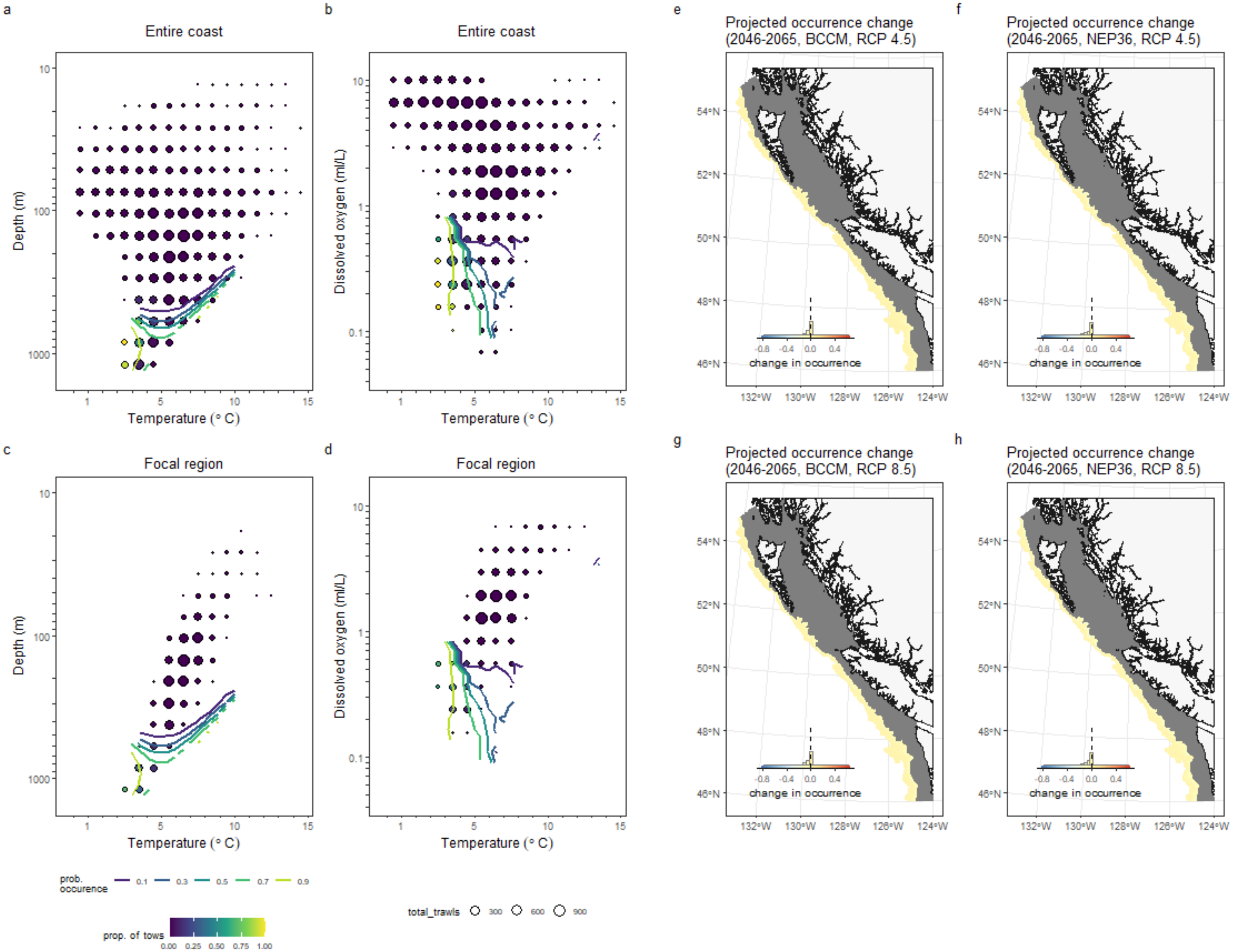

### Redbanded Rockfish

*Sebastes babcocki*

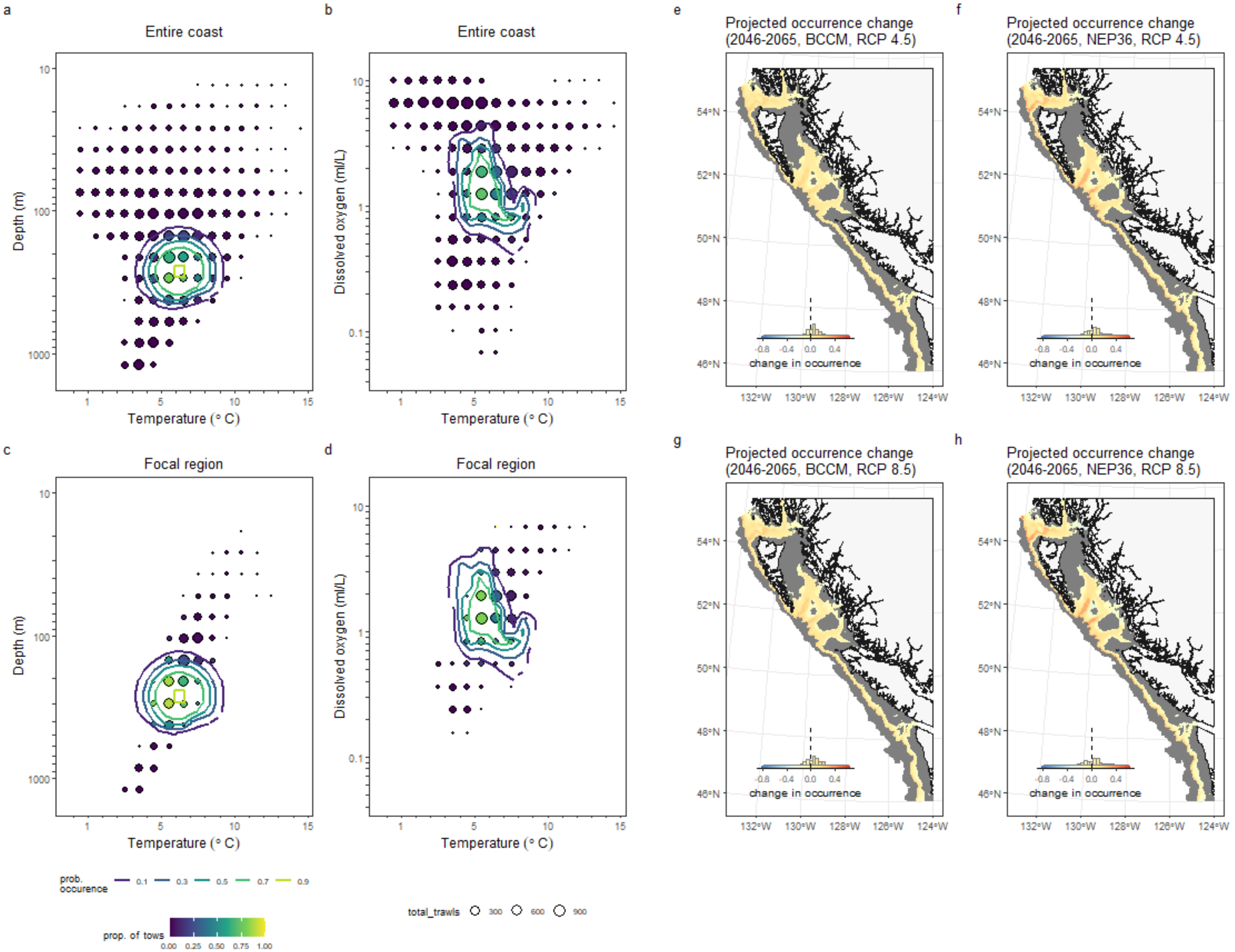

### Rex Sole

*Glyptocephalus zachirus*

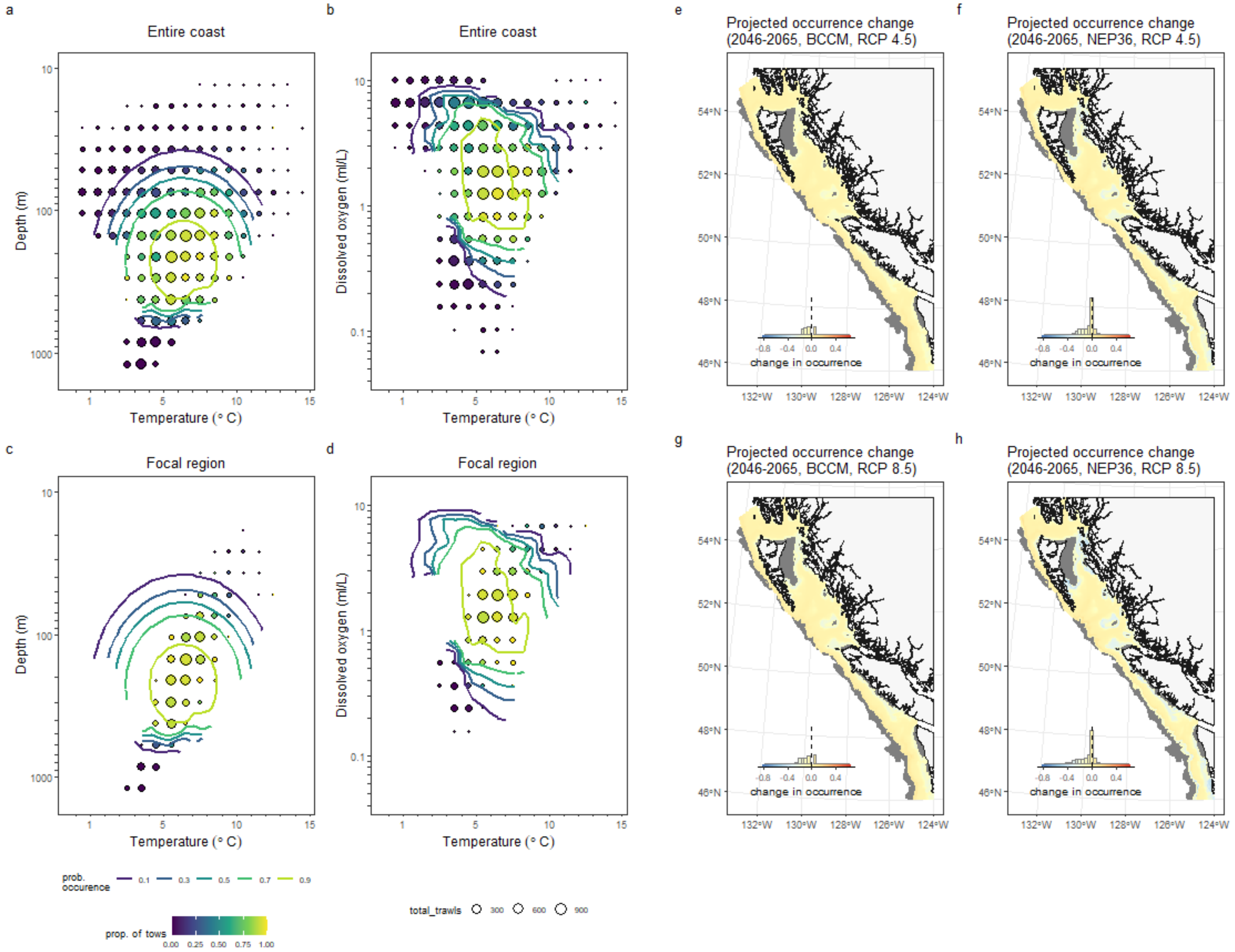

### Rougheye Rockfish Complex

*Sebastes aleutianus/melanostictus*

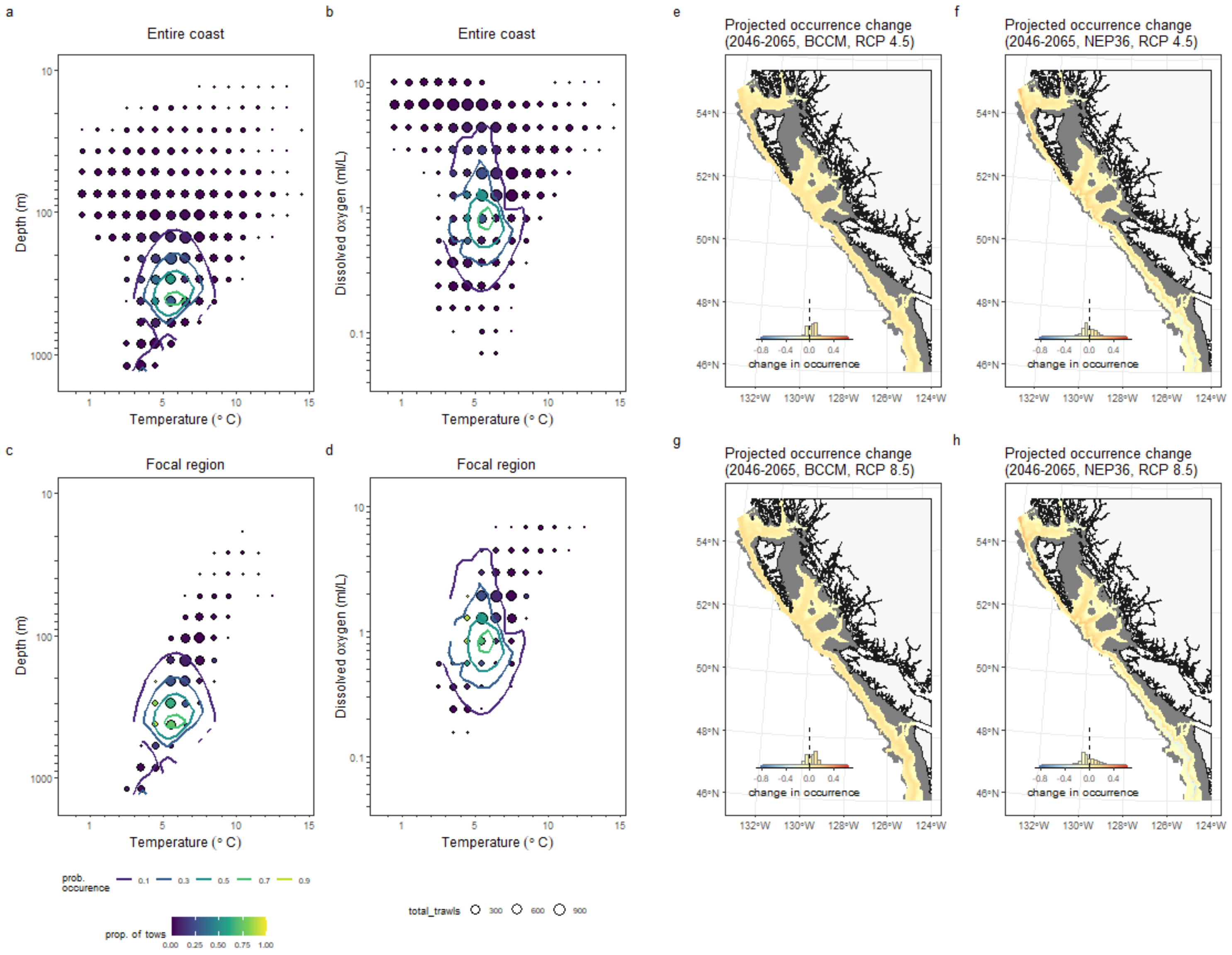

#### Roughtail Skate

*Bathyraja trachura*

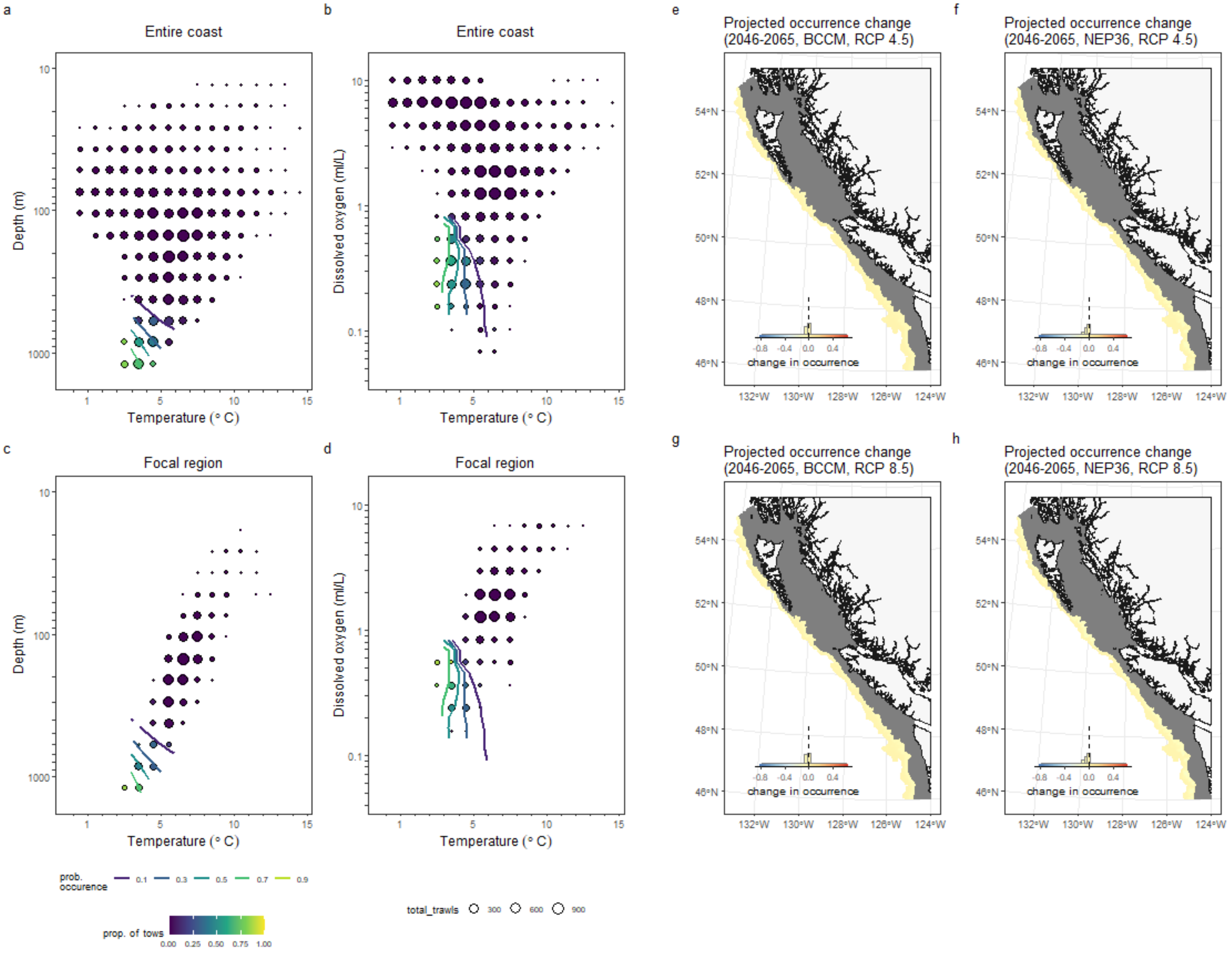

#### Sand Sole
*Psettichthys melanostictus*

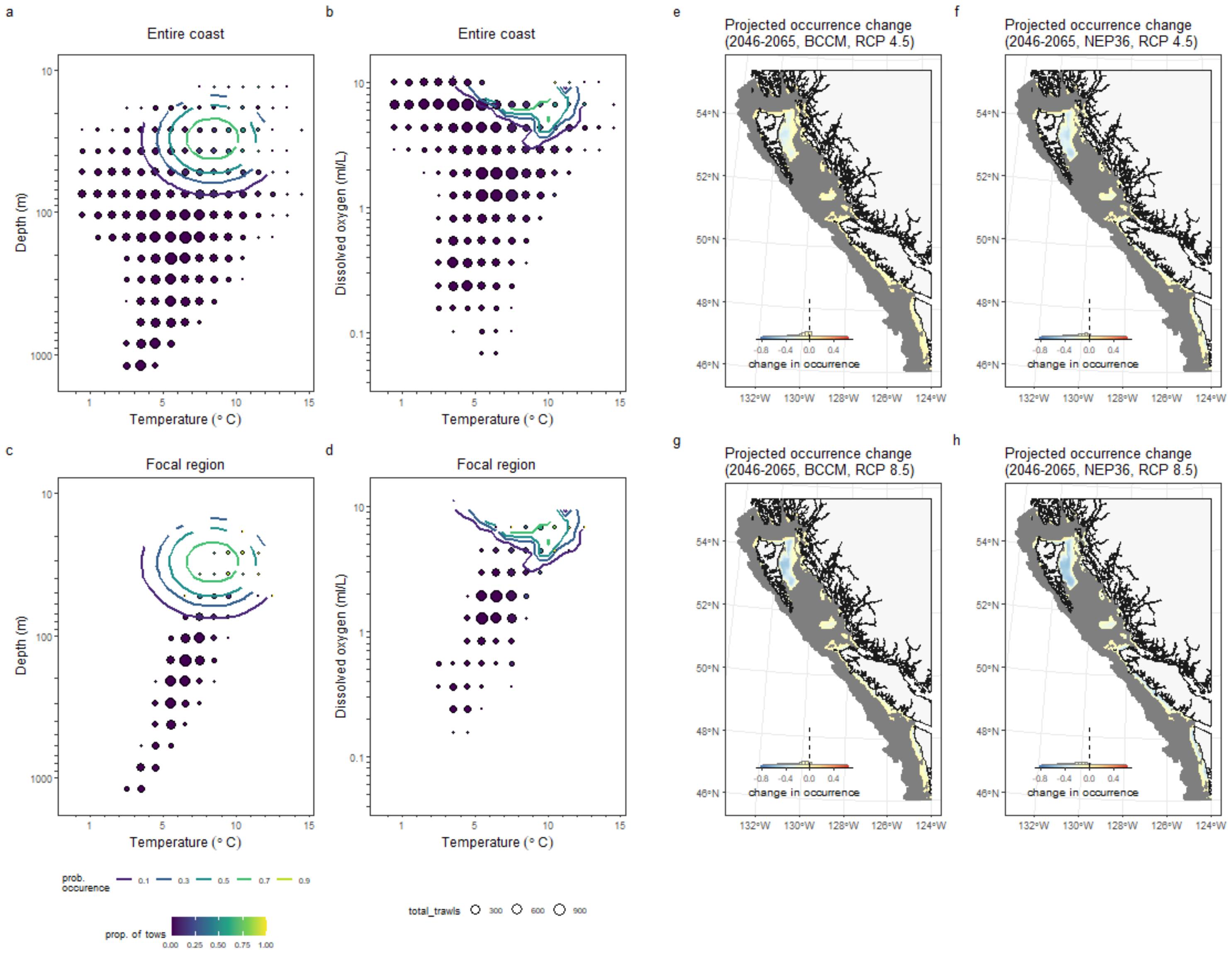

### Sandpaper Skate

*Bathyraja interrupta*

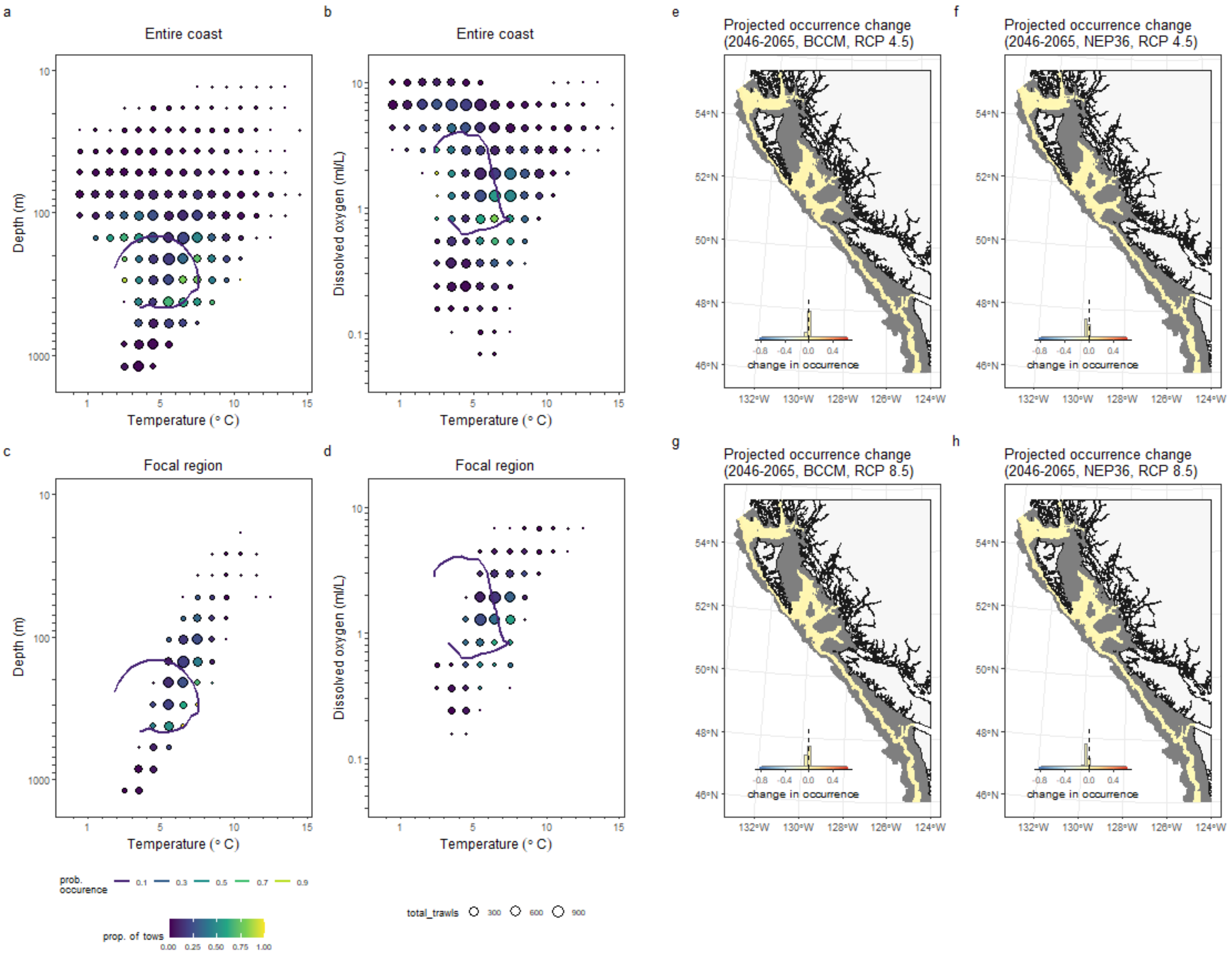

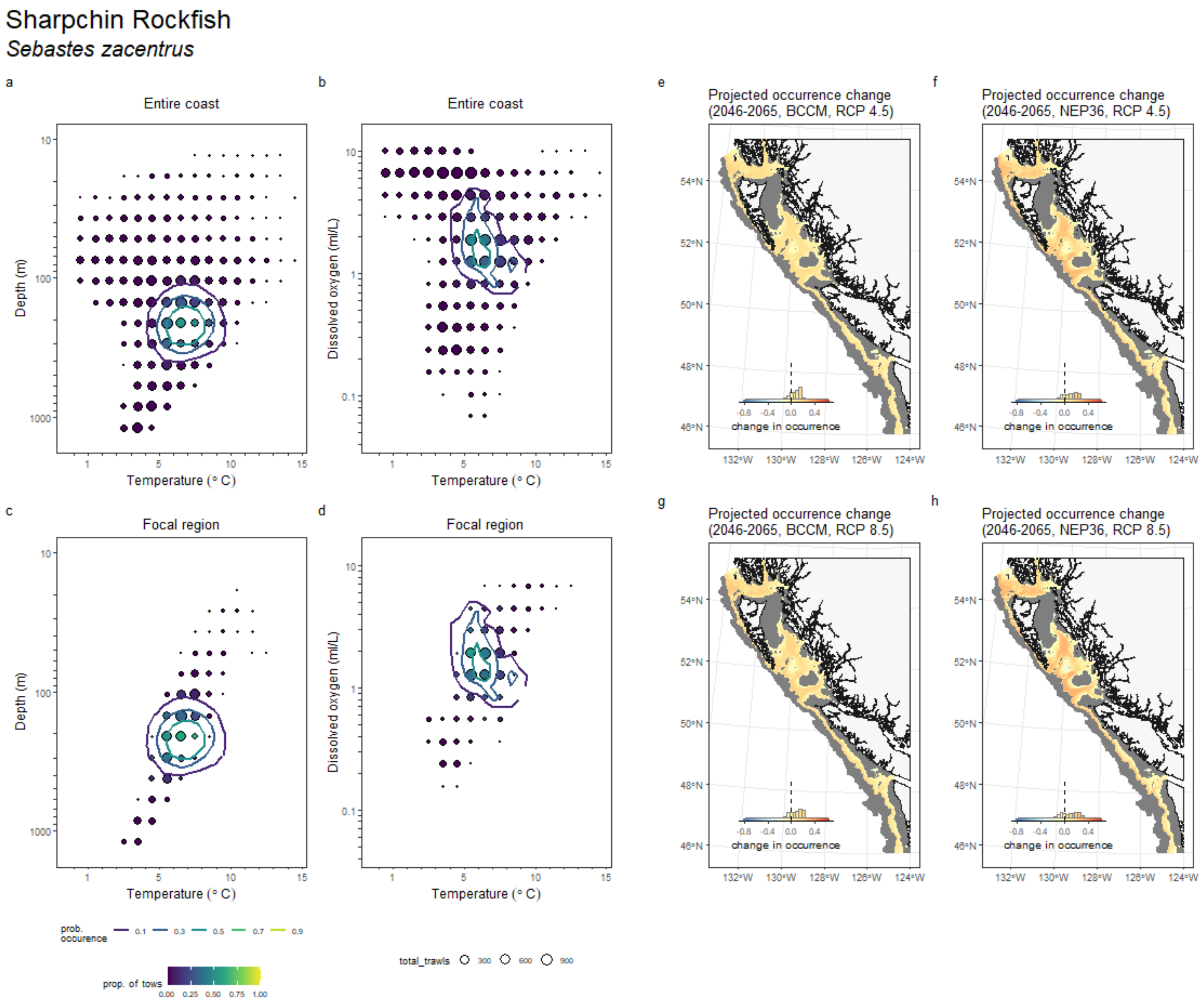
Sharpchin Rockfish
*Sebastes zacentrus*
a
Entire coast
b
Entire coast
c
Focal region
d
Focal region
Depth (m)
Dissolved oxygen (ml/L)
Temperature (° C)
e
Projected occurrence change (2046-2065, BCCM, RCP 4.5)
f
Projected occurrence change (2046-2065, NEP36, RCP 4.5)
g
Projected occurrence change (2046-2065, BCCM, RCP 8.5)
h
Projected occurrence change (2046-2065, NEP36, RCP 8.5)
change in occurrence
prob. occurence
prop. of tows
total_trawls

### Shortspine Thornyhead
*Sebastolobus alascanus*

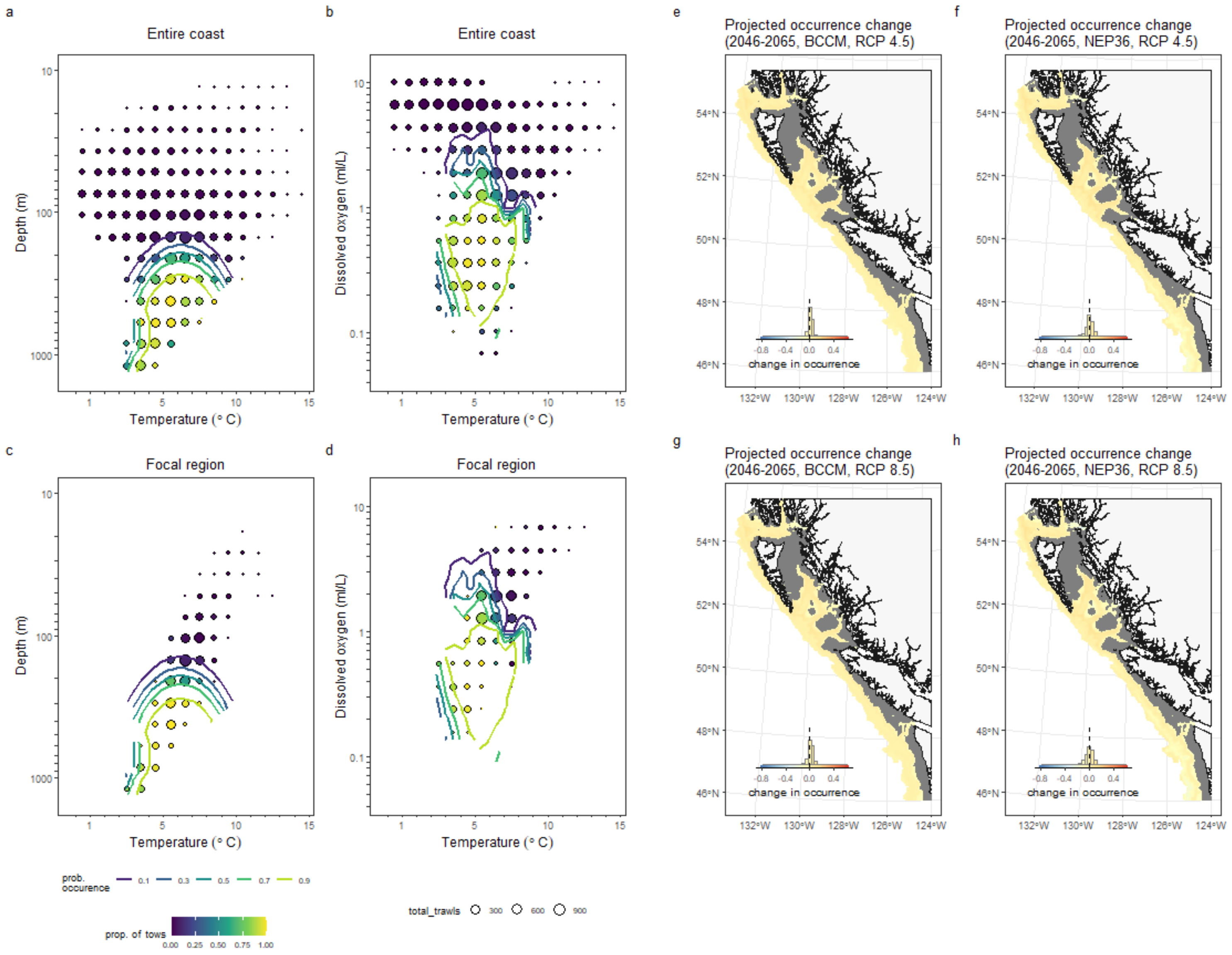

### Silvergray Rockfish

*Sebastes brevispinis*

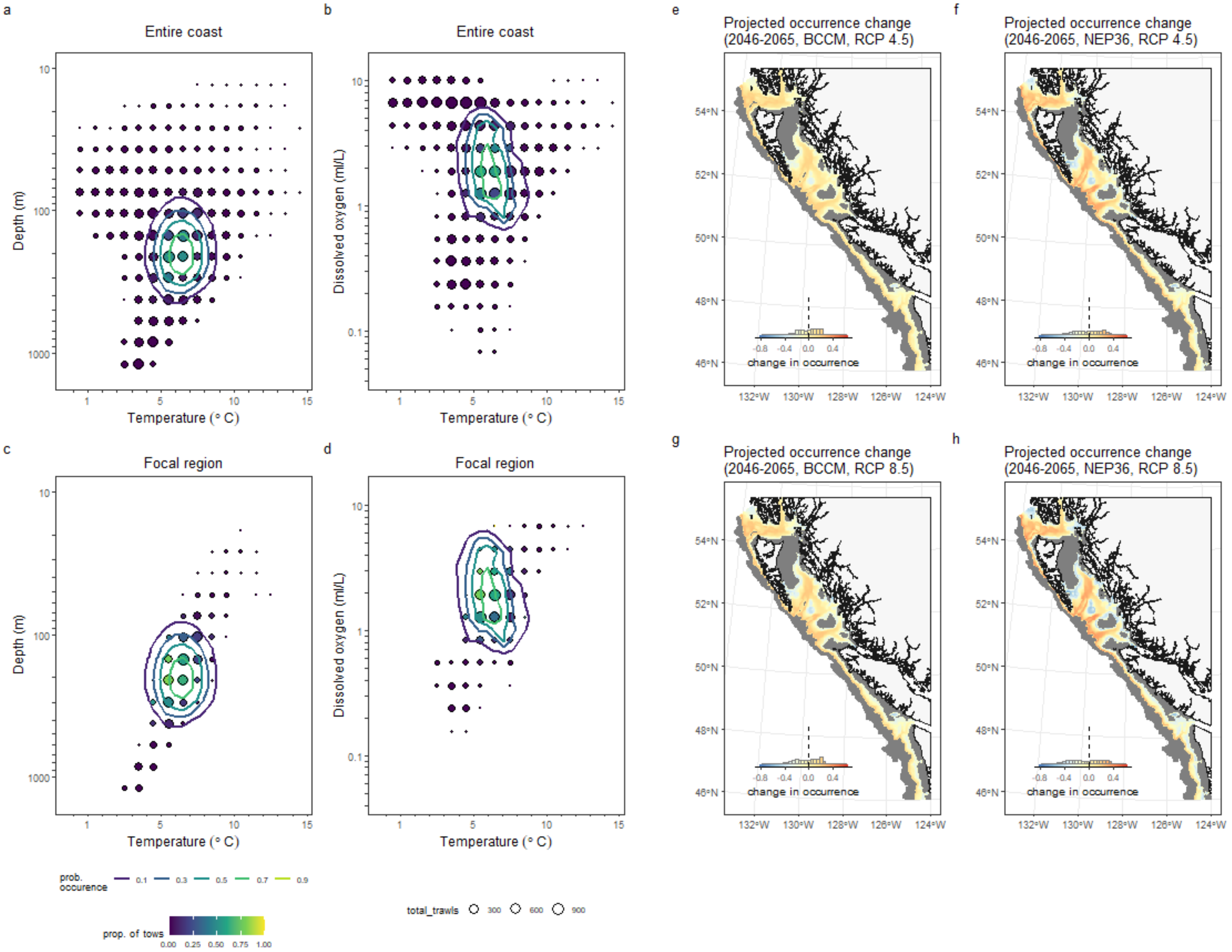

#### Slender Sole
*Lyopsetta exilis*

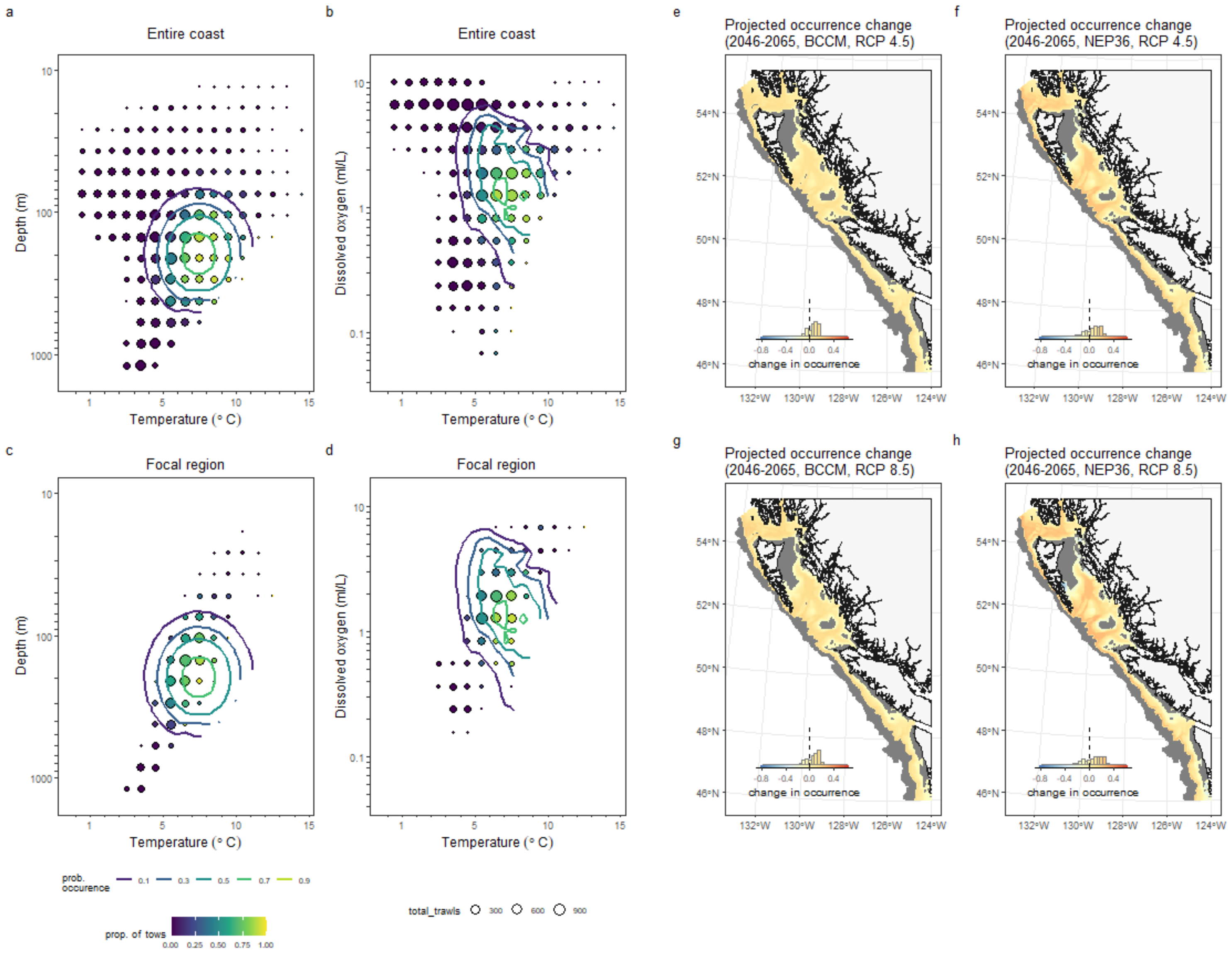

#### Southern Rock Sole
*Lepidopsetta bilineata*

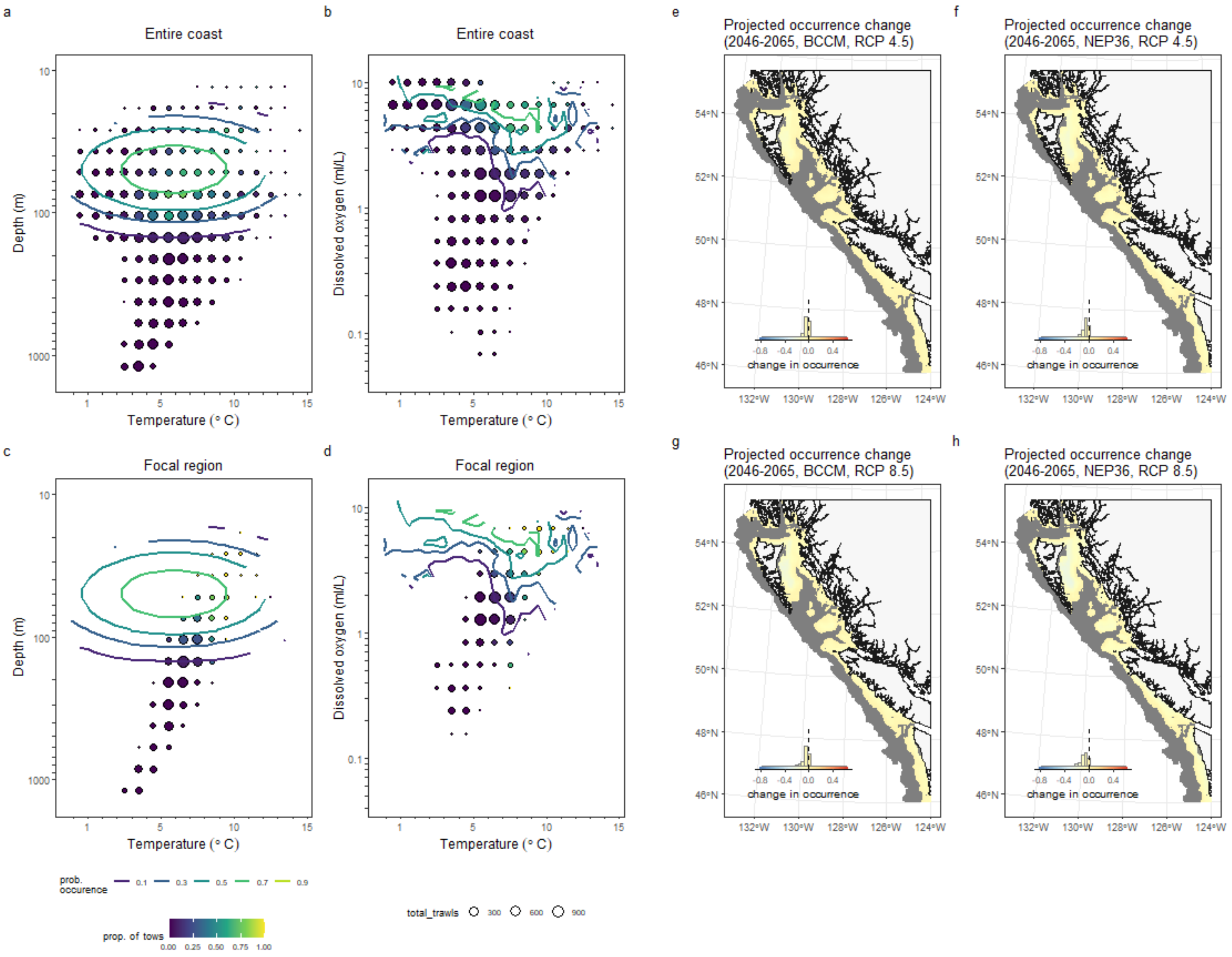

#### Splitnose Rockfish

*Sebastes diploproa*

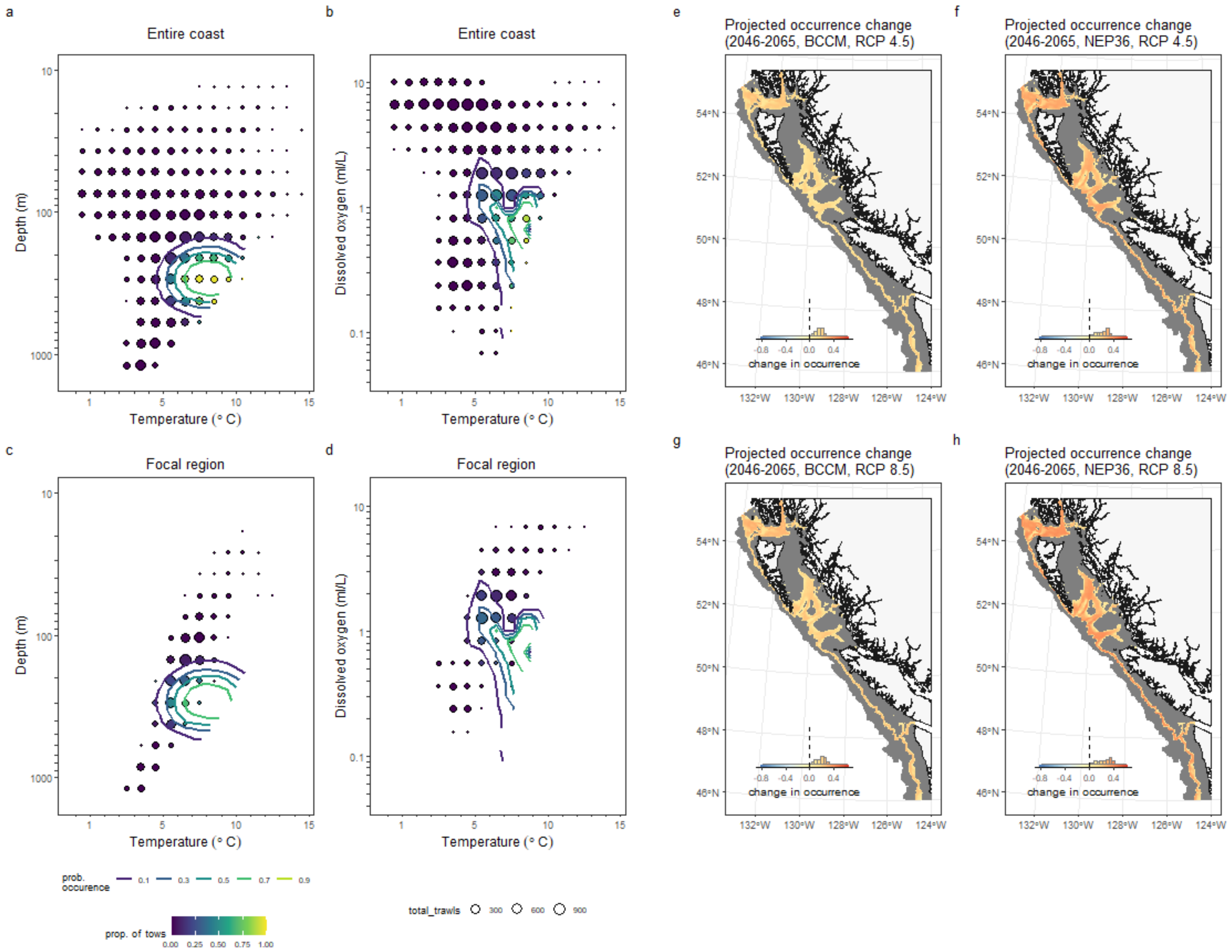

#### Spotted Ratfish

*Hydrolagus colliei*

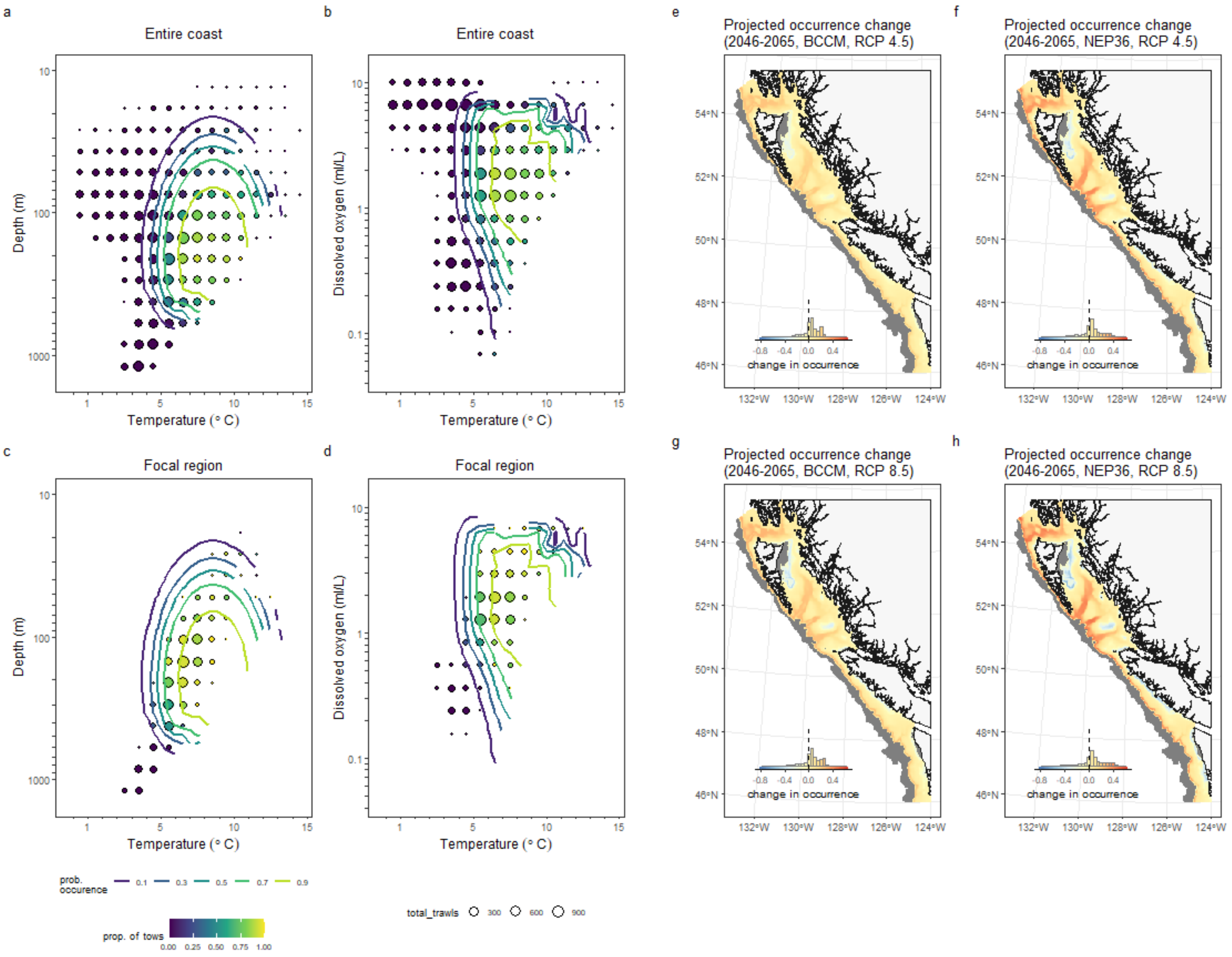


32

#### Sturgeon Poacher

*Podothecus accipenserinus*

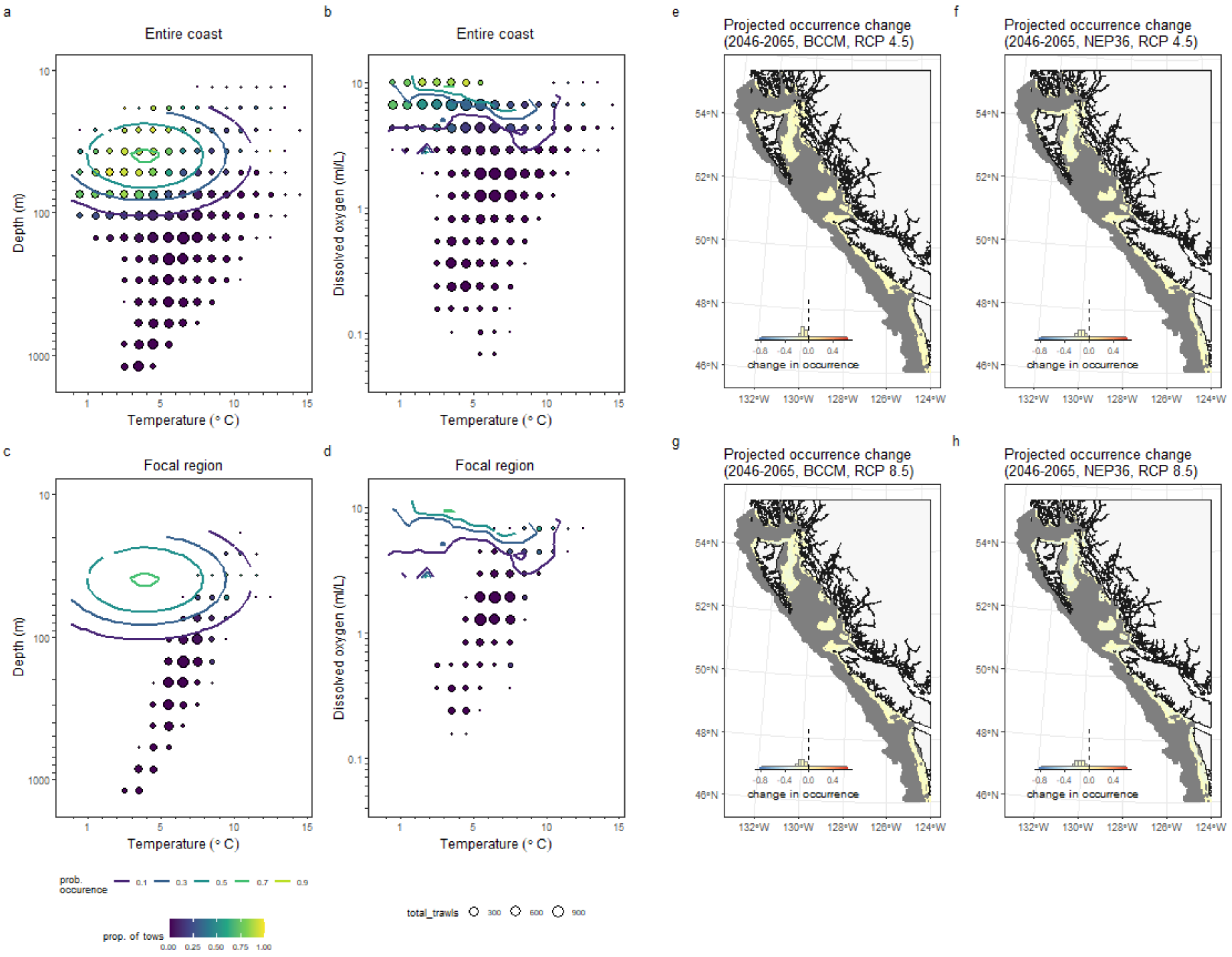

### Twoline Eelpout
*Bothrocara brunneum*

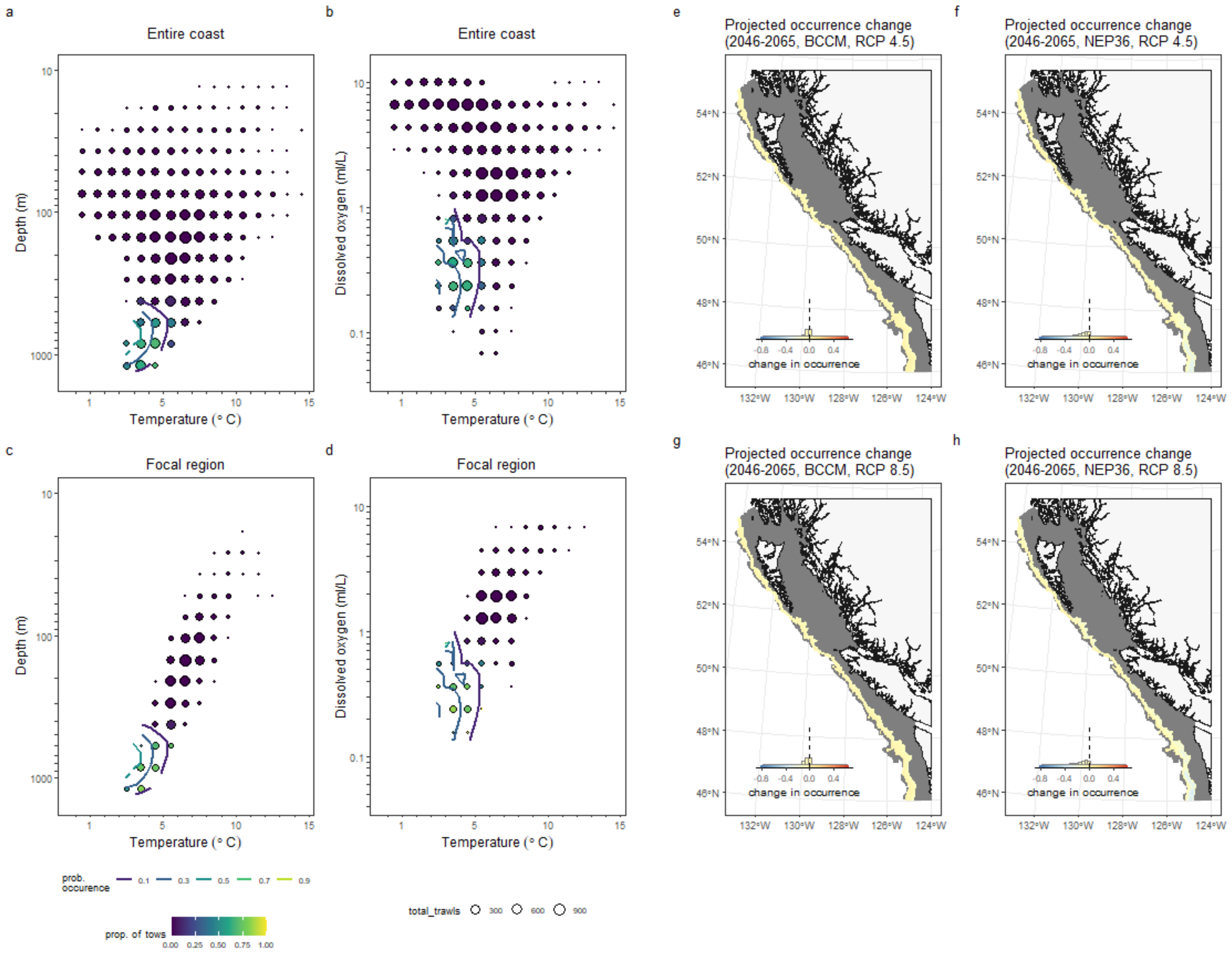

#### Walleye Pollock

*Gadus chalcogrammus*

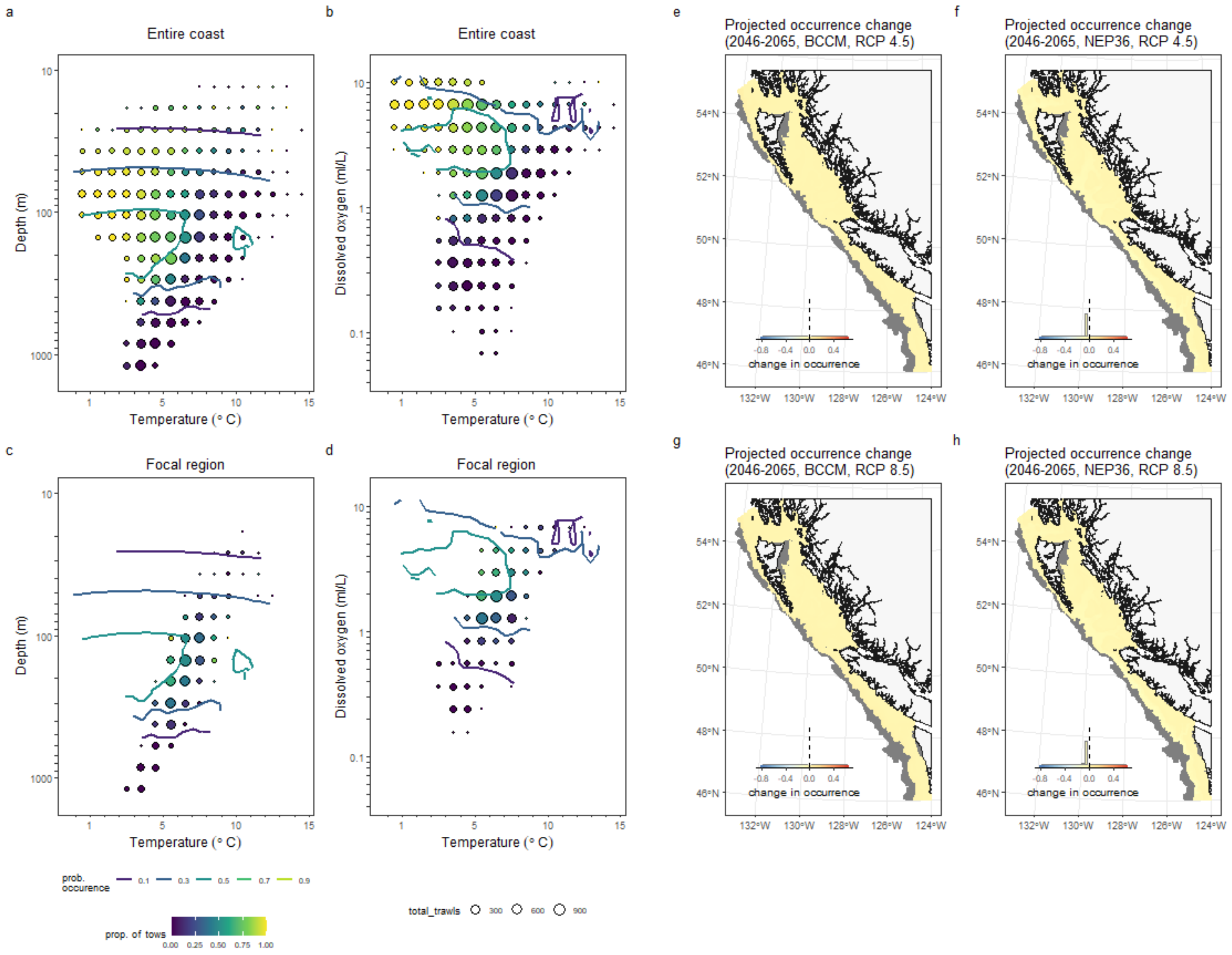

### References

Holdsworth, A.M., Zhai, L., Lu, Y. & Christian, J.R. (2021). Future Changes in Oceanography and Biogeochemistry Along the Canadian Pacific Continental Margin. *Frontiers in Marine Science*, **8**, 602991. Retrieved August 27, 2021, from https://www.frontiersin.org/articles/10.3389/fmars.2021.602991/full

Peña, M.A., Fine, I. & Callendar, W. (2019). Interannual variability in primary production and shelf-offshore transport of nutrients along the northeast Pacific Ocean margin. *Deep Sea Research Part II: Topical Studies in Oceanography*, **169–170**, 104637. Retrieved July 14, 2020, from http://www.sciencedirect.com/science/article/pii/S0967064519300220
